# Phosphoinositide-binding proteins mark, shape and functionally modulate highly-diverged endocytic compartments in the parasitic protist *Giardia lamblia*

**DOI:** 10.1101/741348

**Authors:** Lenka Cernikova, Carmen Faso, Adrian B. Hehl

**Affiliations:** Institute of Parasitology, University of Zurich, Winterthurerstrasse 266a, CH-8057 Zurich, Switzerland

**Keywords:** endocytosis, *Giardia*, phosphoinositide, lipid-binding domain, lipid-strip, peripheral vacuoles, clathrin, epsin, PX domain, STED microscopy, APEX, NECAP1, FYVE

## Abstract

Phosphorylated derivatives of phosphatidylinositol (PIPs), are key membrane lipid residues involved in clathrin-mediated endocytosis (CME). CME relies on PI(4,5)P2 to mark endocytic sites at the plasma membrane (PM) associated to clathrin-coated vesicle (CCV) formation. The highly diverged parasitic protist *Giardia lamblia* presents disordered and static clathrin assemblies at PM invaginations, contacting specialized endocytic organelles called peripheral vacuoles (PVs). The role for clathrin assemblies in fluid phase uptake and their link to internal membranes via PIP-binding adaptors is unknown.

Here we provide evidence for a robust link between clathrin assemblies and fluid-phase uptake in *G. lamblia* mediated by proteins carrying predicted PX, FYVE and NECAP1 PIP-binding modules. We show that chemical and genetic perturbation of PIP-residue binding and turnover elicits novel uptake and organelle-morphology phenotypes. A combination of co-immunoprecipitation and *in silico* annotation techniques expands the initial PIP-binding network with addition of new members. Our data indicate that, despite the partial conservation of lipid markers and protein cohorts known to play important roles in dynamic endocytic events in well-characterized model systems, the *Giardia* lineage presents a strikingly divergent clathrin-centered network. This includes several PIP-binding modules, often associated to domains of currently unknown function that shape and modulate fluid-phase uptake at PVs.

## Introduction

Phosphorylated derivatives of the minor membrane phospholipid phosphatidylinositols (PIPs) are surface molecules of most eukaryotic endomembrane compartments [1–3]. PIPs play important roles in diverse pathways including signaling cascades, autophagy and membrane remodelling [2, 4–8]. Their diverse functions are reflected in their distinct subcellular distribution. PI(4,5)P_2_ is highly enriched at the plasma membrane (PM) with PI(3,4,5)P_3_ [4, 5]. PtdIns(4)P’s largest pool is at Golgi membranes, with smaller amounts found at the the PM. PI(3)P is converted into PI(3,5)P_2_ on early endosomes during transition to multivesicular bodies and then late endosomes [6, 7]. PI(3)P is also a marker of phagosomes [8] while PI(5)P marks both the PM and endomembranes [9]. At least 14 distinct PIP-binding modules haven been identified in eukaryotes, demonstrating a wide range of selective protein-lipid interactions associated with the PM and internal membranes [10].

In addition to their structural functions in membranes, PIPs are involved in spatiotemporal organization of membrane remodeling processes such as clathrin-coated vesicles (CCV) formation during clathrin-mediated endocytosis (CME). In particular, PI(4,5)P2 marks sites of endocytosis at the PM and recruits proteins involved in the formation of CCVs [11]. The protein interactomes of mammalian PI(4,5)P2-binding proteins include the early-acting clathrin interacting partners AP2 [12–15], AP180/CALM [16, 17] and epsin [17, 18]. These factors carry specific PIP-binding domains that can discriminate between PIP variants to achieve membrane targeting specificity.

*Giardia lamblia* (syn. *intestinalis, duodenalis*) is a widespread parasitic protist that colonizes the upper small intestine of vertebrate hosts. Its life cycle is marked by the alternation between an environmentally-resistant, infectious cyst stage responsible for parasite transmission, and a trophozoite stage proliferating by binary fission. Nutrient uptake of trophozoites in the lumen of the small intestine is almost entirely routed through peripheral vacuoles (PVs). These organelles are positioned just beneath the PM and are contacted by funnel-shaped invaginations of the PM that are likely conduits for uptake of fluid-phase extracellular material [19].

A recent characterization of the PV protein interactome using the highly conserved *G. lamblia* clathrin heavy chain (*Gl*CHC) as affinity handle confirmed the endocytic nature of these organelles by highlighting the presence of giardial AP2 (*Gl*AP2) subunits, the single dynamin-like protein GlDRP and a putative clathrin light chain *Gl*4259 (*Gl*CLC; [19]). Notably absent were components for CCV uncoating and disassembly, consistent with a lack of measurable clathrin assembly turnover and in line with the observations that CCVs are missing in *G. lamblia* and clathrin assemblies are static and long-lived. Therefore, *G. lamblia* presents an unusual endocytic system, characterized by divergent endocytic compartments (PVs) associated to static clathrin assemblies that are not predicted to form ordered arrays or higher-order structures such as CCVs yet are closely membrane-associated.

Included in the giardial CHC interactome were three proteins with predicted PIP-binding domains: FYVE domain protein *Gl*16653 and two PX-domain proteins (*Gl*7723 and *Gl*16595), the latter part of a six-member protein family (Table 1; [19, 20]). In a previous study, we hypothesized that *Gl*16653 (*Gl*FYVE), *Gl*7723 (*Gl*PXD1) and *Gl*16595 (*Gl*PXD2) act as PIP-binding adaptors to link and maintain static clathrin assemblies at the PM and PV membrane interface in *G. lamblia* [19]. We further hypothesized that a perturbation of PIP-binding protein levels and/or function would lead to impaired fluid-phase uptake by affecting PV functionality. To test these hypotheses, we performed an in-depth functional characterization of all previously-identified PIP-binding proteins associated to clathrin at PVs. We assessed their lipid-binding preferences and visualized their subcellular localizations using electron microscopy and both conventional and super resolution light microscopy. By manipulating protein levels and/or function we could elicit novel fluid-phase uptake and PV morphology-related phenotypes, thereby establishing PIPs as a link between the role of clathrin as a membrane remodeling proteins and PV-based endocytosis in *G. lamblia*. Furthermore, we used a combination of co-immunoprecipitation and *in silico* annotation techniques to expand protein interactomes established previously, thereby discovering a new set of PIP-binding proteins with roles likely reaching beyond the PV compartment. Lastly, we propose an updated working model summarizing the complex networks between PIP-binding proteins and clathrin assemblies at PVs.

**Table 1:**
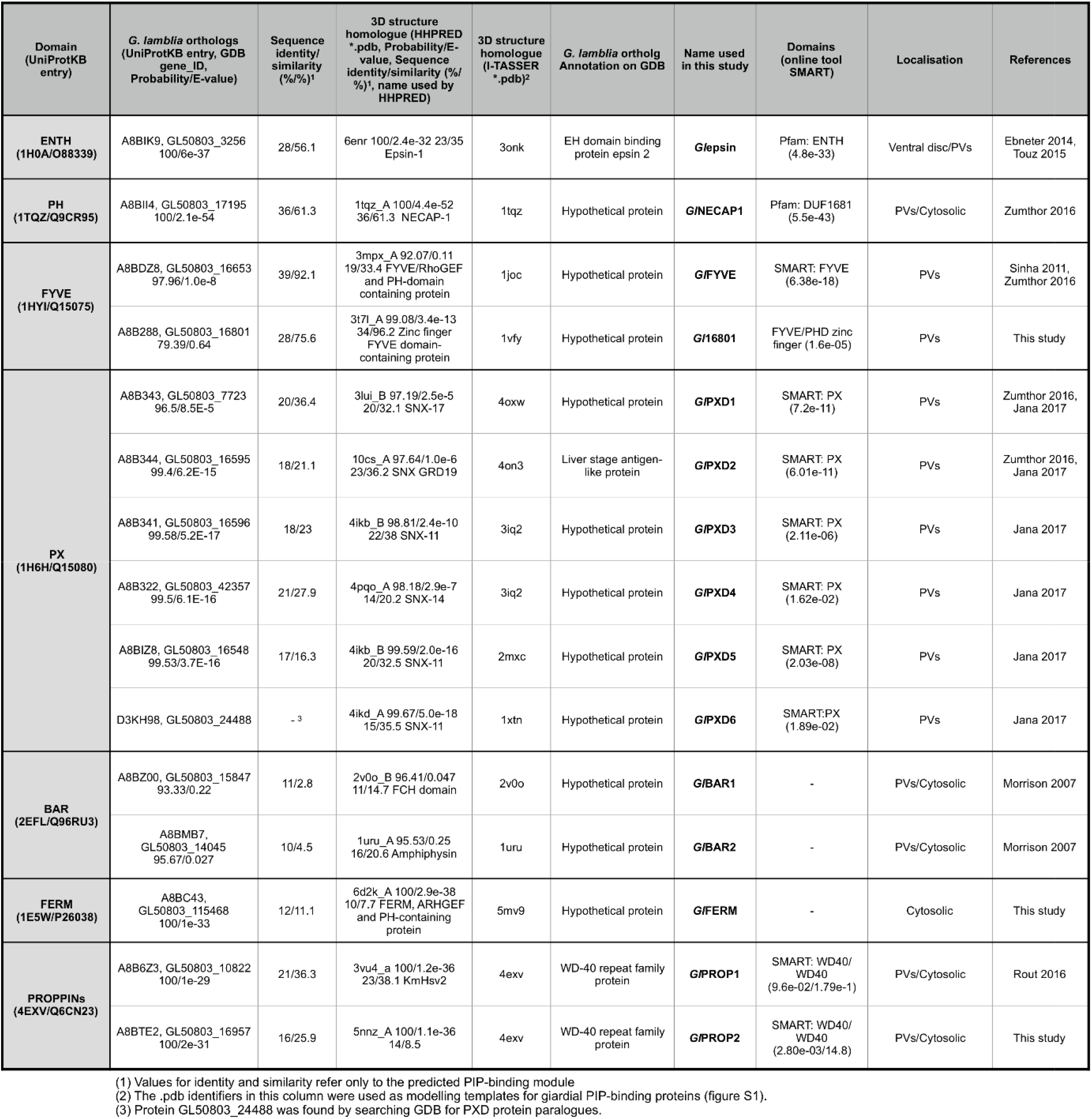
*G. lamblia* PIP-binding proteins. A compilation of all PIP-binding domains identified in the *Giardia* Genome Database (www.giardiadb.org; GDB) using previously characterized domains [24] as baits for HMM-based homology searches (column 1). Predicted *Giardia*l orthologs are present for PIP-binding domains ENTH, PH, FYVE, PX, BAR, FERM and PROPPINS (column 2) and mostly retrieve the correct domains when used as a baits for reverse HHpred searches (column 4). Except for *Gl*epsin, *Gl*PXD2 and *Gl*PROP1 and 2, all others are currently annotated on GDB as generically “hypothetical”, i.e. of unknown function (column 6). Each orthologue was assigned a name used throughout this report (column 7). Functional domain predictions using SMART (http://smart.embl-heidelberg.de/; column 8) and subcellular localization data (column 9) either previously reported or acquired in this study (column 10), are also included.

## Results

### The *G. lamblia* genome encodes at least seven distinct PIP-binding modules

Given that several types of PIP-binding modules have been identified in eukaryotes, we determined how many endocytosis-associated module types were actually represented in the *Giardia* genome, in addition to the known *G. lamblia* epsin, FYVE and PXD variants [19–23]. For this reason, we selected a total of 14 protein types from various organisms known to harbor PIP-binding domains, some of them involved in endocytosis. These are: ANTH (AP180 N-terminal homology), ENTH (epsin N-terminal homology), PH (Pleckstrin homology domain), FYVE (Fab1, YOTB, Vac1 and EEA1), PX (Phox homology), BAR (bin, amphiphysin and Rvs), FERM (4.1, ezrin, radixin, moiesin), PROPPINs (β-propellers that bind PIs), C2 (conserved region-2 of protein kinase C), GOLPH3 (Golgi phosphoprotein 3), PDZ (postsynaptic density 95, disk large, zonula occludens), PTB (phosphotyrosine binding), Tubby modules and the PH-like module of the endocytosis-associated NECAP1 protein [24]. Representatives for each module were used as bait for the HMM-based tool HHpred [25] for protein structure prediction and the detection of remotely related sequences in the *G. lamblia* predicted proteome (Table 1). Putative *Giardia* protein homologs (Table 1) were then subjected to the online tools SMART [26, 27] and InterProScan [28] to identify, conserved structural domains and sequence motifs within a query sequence (Fig 1A).

**Figure 1:**
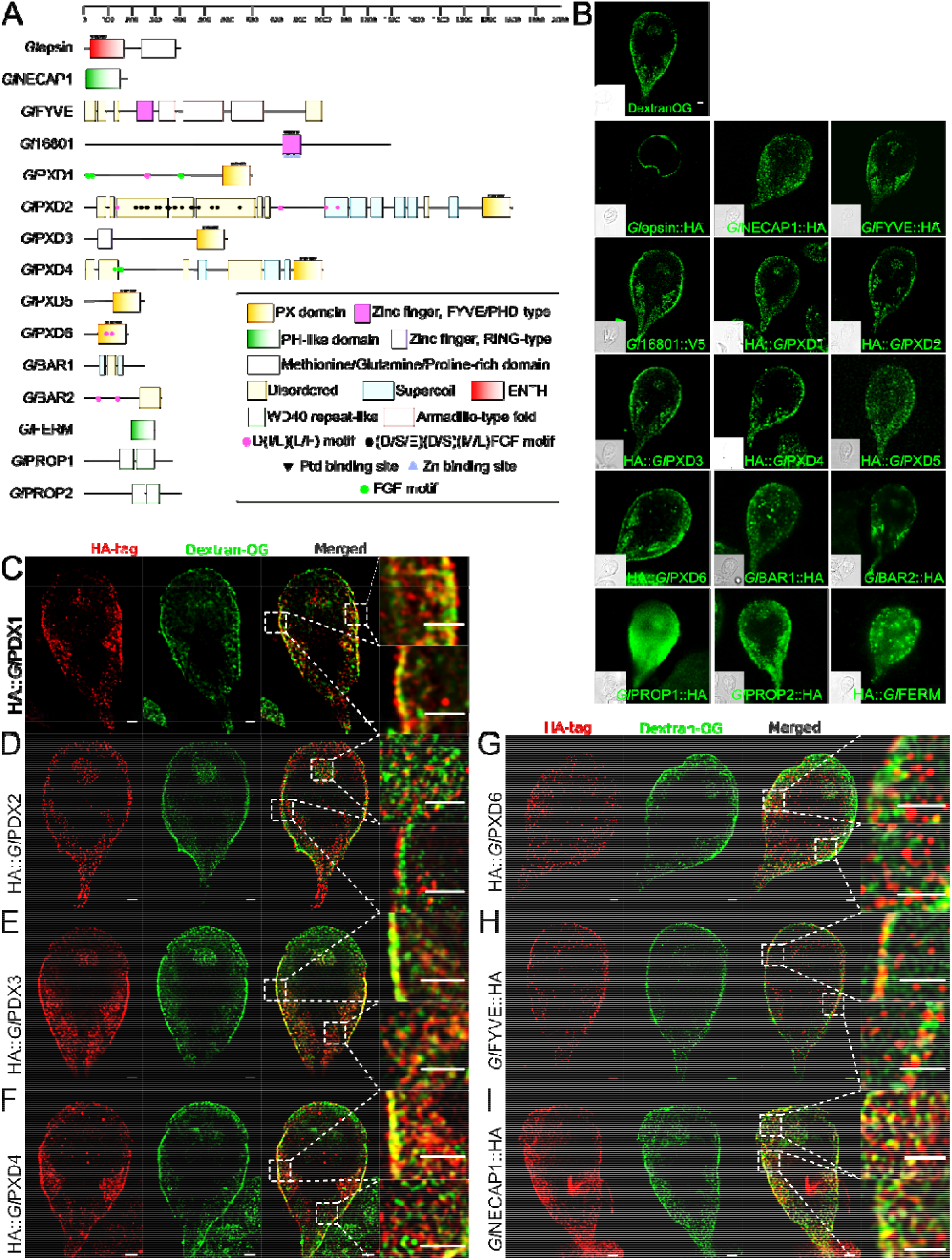
Functional domain prediction analysis and subcellular localization of *G. lamblia* PIP-binding proteins. (A) Predicted functional domains for all identified PIP-binding proteins including positions of repetitive motifs and putative lipid and Zn -binding residues using HHPRED, HMMER and InterProScan. Ptd – Phosphatidylinositol. (B) Conventional confocal light-microscopy analysis of representative non-transgenic trophozoites labelled with Dextran-OG (first panel) to mark PV lumina and of immune-labelled trophozoites expressing HA-tagged PIP-binding protein reporters. and. Except for *Gl*epsin and *Gl*FERM, all tested reporter proteins localise in close proximity to peripheral vacuoles (PVs) at the cell cortex. Epitope-tagged *Gl*NECAP1, *Gl*PXD5 and *Gl*PROP1 additionally show signal distribution throughout the cell. Cells were imaged at maximum width, where nuclei and the bare-zone are at maximum diameter. Epitope-tagged *Gl*epsin-expressing cells were imaged at maximum width of the ventral side. Insets: DIC images. Scale bar: 1 µm (C) Confocal STED microscopy analysis of trophozoites expressing clathrin assemblies-associated epitope-tagged PIP-binding reporter proteins for *Gl*PXD1-6, *Gl*FYVE and *Gl*NECAP1 (red channel) co-labelled with Dextran-OG as a marker for PV lumina (green channel). As shown in the merged insets, although all reporters are clearly PV-associated, reporters for proteins *Gl*FYVE and *Gl*PXD1 and 2 are proximal to the PM with respect to Dextran-OG, indicating they reside at the PV-PM interface. In contrast, reporters for *Gl*PXD3 and *Gl*NECAP1 appear to intercalate PVs. Scale bars: 1 µm for full cell image, 1 µm for insets.

This data mining approach detected high-confidence homologs for hitherto undiscovered *G. lamblia* proteins containing PH-like, FERM, BAR, FYVE and Proppin PIP-binding domains (Table 1, Fig 1A). No homologs could be found for the ANTH, C2, GOLPH3, PDZ, PTB, Tubby and PH PIP-binding module types.

Protein GL50803_17195 (*Gl*NECAP1) is a predicted NECAP1 homolog containing a PH-like domain. Similarly, a conserved PH-like domain found at the C-terminus of FERM proteins was correlated with high confidence to protein GL50803_115468 (*Gl*FERM). Immunofluorescence assays (IFAs) and confocal microscopy imaging of an epitope-tagged *Gl*NECAP1 reporter expressed as an extra copy under its own promoter showed localization in close proximity to PVs and in the cytosol (Fig 1B). Similar to *Gl*NECAP1, BAR domain-containing proteins GL50803_15847 and GL50803_14045 (*Gl*BAR1 and 2) localize in close proximity to PVs (Fig 1B). Tagged reporters for new FYVE and PROPPIN members GL50803_16801 (*Gl*16801) and GL50803_16957 (*Gl*PROP2), respectively, both localise in close proximity to PVs (Fig 1B). In contrast, a tagged *Gl*FERM reporter presents a diffused cytosolic subcellular distribution (Fig 1B).

To extend the initial annotation of giardial PIP-binding proteins we performed multiple sequence alignment (MSA) analyses for each giardial PIP-binding module with selected orthologs to delineate lipid-binding motifs and residues critical for PIP recognition (Fig S1). *In silico* structural analyses of the lipid-binding domains of giardial proteins and their closest homologs were performed *ab initio* using the online tool I-TASSER [29–31]. Comparative analysis of structure models generated with I-TASSER clearly demonstrated positional conservation of residues critical for PIP binding (Fig S1). Since *Gl*PXD1-2, *Gl*FYVE and *Gl*NECAP1 were experimentally shown to be associated to giardial clathrin assemblies [19], we selected these proteins and *Gl*PXD3-6 for more detailed subcellular localization experiments. Stimulated emission-depletion (STED) microscopy in co-labelling experiments with Dextran-OG as a marker for fluid-phase endocytosis, unequivocally confirmed accumulation for *Gl*PXD1-4 and 6, *Gl*FYVE and *Gl*NECAP1 epitope-tagged reporters at PVs (Fig 1C). The signal generated by *Gl*PXD5 reporters was insufficient for a conclusive localization.

Taken together, *in silico* analysis identifies seven distinct PIP-binding module types encoded in the *G. lamblia* genome, conserved on both sequence and structural levels. Subcellular localization of epitope-tagged variants by fluorescence microscopy indicates clear association to PVs with the exception of *Gl*epsin.

### PIP-binding proteins associated with clathrin assemblies present distinct lipid-binding profiles in vitro

PX domains [32] and FYVE [33–35] preferentially bind PI(3)P. Even though PH domains have rather promiscuous binding preferences, a subset of PH domains binds strongly to PtdIns(3,4,5)P3 and PtdIns(4,5)P2, as well as PtdIns(3,4)P2 [36–38]. Based on the presence of conserved residues for lipid-binding in the giardial PXD1-6, FYVE and NECAP1 proteins (Fig S1), we hypothesized that their lipid-binding preferences would also be conserved. We tested this experimentally by expressing MBP-fused, epitope-tagged *Gl*PXD1-6, *Gl*FYVE and *Gl*NECAP1 lipid-binding domains (Fig S2A and B). The recombinant fusion proteins were affinity-purified and used in lipid binding assays either for commercially-available PIP gradients as membrane-supported arrays (1.56-100 pmol/spot) (Fig 2A) or membrane strips spotted with defined amounts (100 pmol/spot) of PIPs (Fig S2C). The negative control for binding consisted of a PIP array probed with purified epitope-tagged MBP alone, whereas the positive control consisted of a PIP array probed with a commercially-available anti-PI(4,5)P_2_ antibody (Fig 2A). Quantification of the chemiluminescence signals shows a marked preference of MBP-*Gl*PXD1 for PI(4,5)P_2_ in PIP gradients (Fig 2B) which was corroborated by experiments using PIP strips (Fig S2C). Under these conditions, *Gl*PXD2, 3 and 6 show unexpectedly promiscuous binding preferences, with *Gl*PXD2 presenting a marked affinity for PI(3)P and PI(4,5)P_2,_ *Gl*PXD3 for PI(3)P and to a lesser extent PI(5)P, and *Gl*PXD6 for PI(3)P, PI(4)P and PI(5)P (Fig 2B). These data were in line with results from independent PIP strip experiments (Fig S2C and D). MBP-*Gl*PXD4 and MBP-*Gl*PXD5 binding preferences could only be probed using PIP strips (Fig S2C), showing in both cases a marked affinity for PI(3,5)P_2_ and PI(4,5)P_2_ (Fig S2C and D). Binding preferences for MBP-*Gl*FYVE could not be determined, given that no signal was ever obtained on both PIP arrays and strips (Fig 2A, Figs S2C and E). Surprisingly, testing of *Gl*NECAP1 consistently detected cardiolipin as the preferred lipid moiety (Fig 2C; Fig S2E), with no detectable preference for PIP residues (Fig S2C). Taken together, our data shows clearly distinguishable lipid binding profiles in vitro, with varying degrees of promiscuity for different PIP-binding domains.

**Figure 2:**
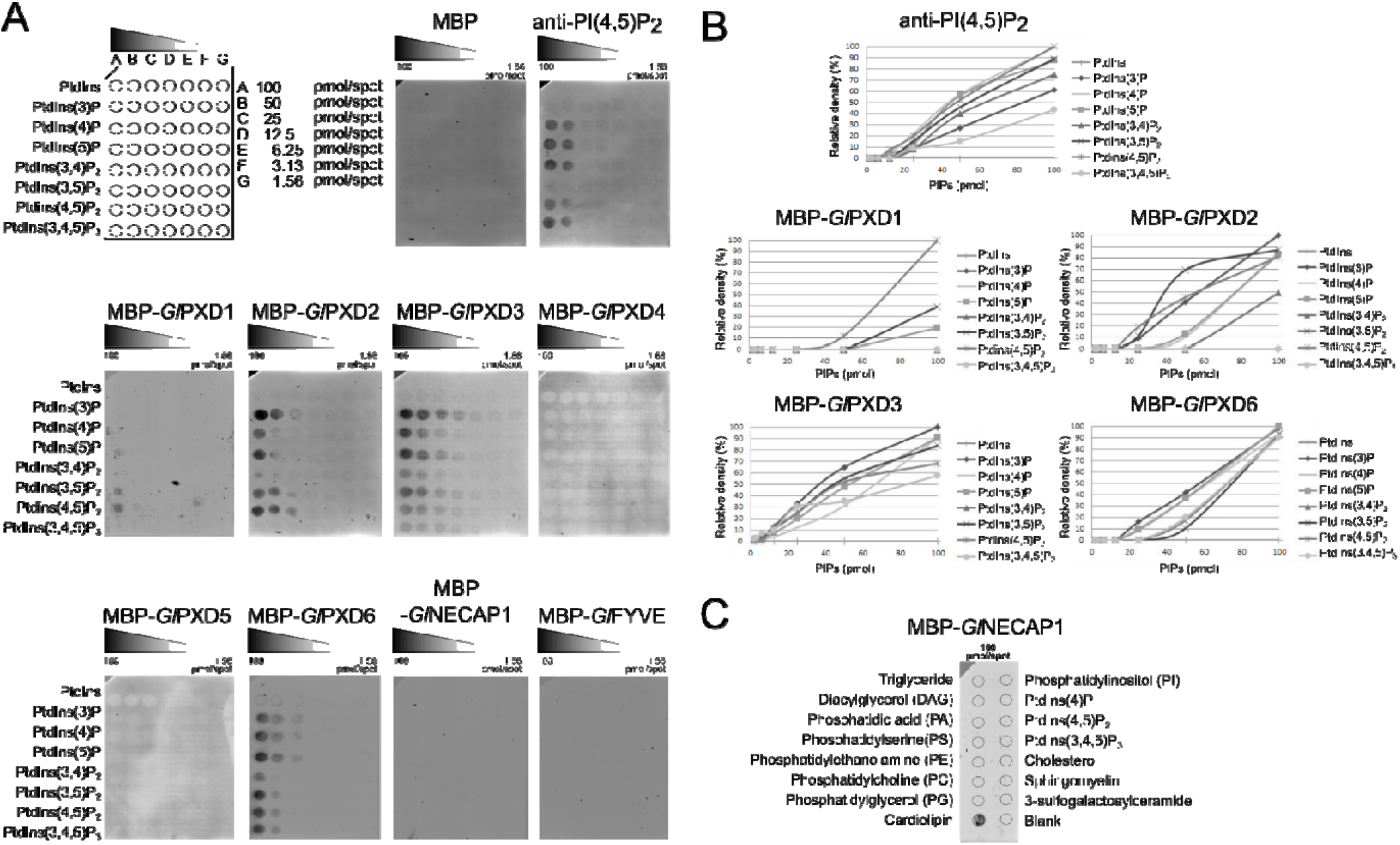
Lipid-binding properties of selected giardial PIP-binding domains. (A) Membrane-supported lipid arrays spotted with gradients of different phosphorylated variants of phosphatidylinositol (PtdIns), from 100pmol (A) to 1.56pmol/spot (G), were probed with fixed amounts (2.5 µg) of clathrin assemblies-associated epitope-tagged PIP-binding domains from *Gl*PXD1-6, *Gl*NECAP1 and *Gl*FYVE, followed by immunodetection of the epitope tag. Lipid-binding preferences for the protein fusion partner MBP (MBP alone) and for antibodies raised against PI(4,5)P2 (anti-PI(4,5)P2) were included as negative and positive controls for binding, respectively. No signal using arrays was obtained for MBP::*Gl*PXD4 and MBP::*Gl*PXD5, however, binding preferences for these fusions were determined using lipid strips (Figs S2A and B). (B) Plots of densitometric analyses using FIJI for each MBP-fused PIP-binding domain and each spotted PI/PIP residue based on array data presented in (A). (C) Testing of the binding affinity of the MBP-fused PIP-binding domain from *Gl*NECAP1 on a wider range of lipid residues detects cardiolipin as the preferred substrate.

### Saturation of PI(3)P, PI(4,5)P_2_ and PI(3,4,5)P_3,_ but not PI(4)P binding sites in vivo inhibits PV-mediated uptake of a fluid-phase marker

The marked preference of *Gl*PXD1-6 for PIP residues PI(3)P and PI(4,5)P_2_ raised the question whether their saturation of in vivo would elicit loss of function phenotypes in fluid phase uptake by *Giardia* trophozoites. Using a combination of commercially available antibodies, heterologous reporter constructs and chemical treatment we saturated sites of PI3P, PI(4,5)P_2,_ and in addition PI(3,4,5)P_3_ and PI(4)P.

Detection of PI(3)P, PI(4,5)P_2_ and PI(3,4,5)P_3_ in chemically fixed trophozoites by immunofluorescence microscopy with primary PIP-targeted antibodies highlights enrichment for all PIP moieties in the cortical region containing PVs (Fig S3).

Ectopic expression of fluorescent high-affinity reporters for PI(3)P and PI(4)P 2xFYVE::GFP and GFP::P4C [39], respectively, in transgenic *G. lamblia* trophozoites was used to both identify as well as saturate membranes enriched for PI(3)P and PI(4)P deposition (Figs 3A-D). Live microscopy of cells expressing 2xFYVE::GFP shows distinct reporter accumulation in cortical areas consistent with binding to PV membranes (Fig 3B, green panels), whereas representative cells from line GFP::P4C show a more diffused cytosolic staining pattern, with some accumulation at PVs (Fig 3D, green panels). Fluid-phase uptake of Dextran-R was assessed in cells from both transgenic lines and compared to wild-type cells using quantification of signal intensity. Wild-type control cells and transgenic cells expressing small amounts of 2xFYVE::GFP (Fig 3A) incorporated large amounts of Dextran-R (Fig 3E). Conversely, a strong 2xFYVE::GFP signal correlated with low amounts of endocytosed Dextran-R detected at the cell periphery and with noticeably enlarged cells (Fig 3B). In contrast, there was no detectable difference in either Dextran-R uptake efficiency (based on fluorescent signal intensity) or cell width between weak (Fig 3C) and strong expressors (Fig 3D) of the GFP::P4C line. Cell width (Fig 3F) and fluid-phase uptake (Fig 3G) aberrant phenotypes in 2xFYVE::GFP cells were recorded with respect to wild-type control and GFP::P4C cells and tested for significance (p>0.05) on 100 cells/line selected in an unbiased fashion. These data translate into a significant negative correlation between expression of the PI(3)P-binding 2xFYVE::GFP reporter and fluid-phase uptake (Fig 3H) whereas only a slight albeit insignificant correlation was found between Dextran uptake and GFP::P4C expression (Fig 3I).

**Figure 3:**
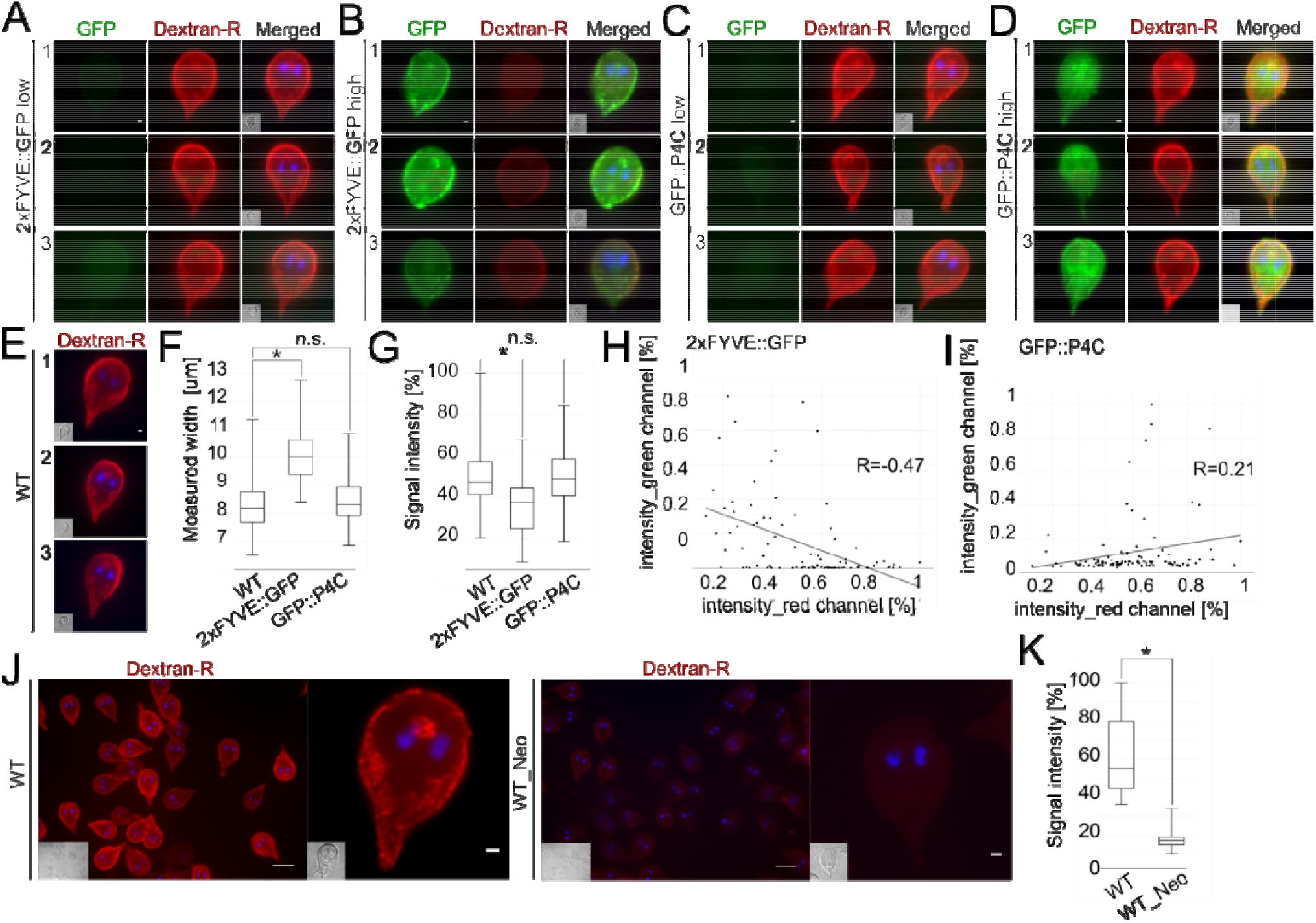
Saturation of PI(3)P, PI(4,5)P2 and PI(3,4,5)P3 binding sites in *G. lamblia* trophozoites elicits uptake and morphological phenotypes. Light microscopy-based immunofluorescence analysis of representative transgenic trophozoites expressing *Legionella*-derived PIP-binding constructs. (A-B) Compared to low 2xFYVE::GFP-expressing cells from the same population, saturation of PI(3)P binding sites in cells highly overexpressing a regulated encystation-dependent epitope-tagged construct 2xFYVE::GFP (anti-HA) inhibits uptake of fluid-phase marker Dextran-R. Scale bars: 1 µm. (C-D) Expression levels of PI(4)P-binding epitope-tagged construct GFP::P4C expression (anti-HA) have no visible impact on Dextran-R signal at PVs of transfected cells. Scale bars: 1 µm. (E) Dextran-R uptake in non-transgenic wild-type cells as negative controls for construct-induced uptake phenotypes. Scale bars: 1 µm (F) Box-plot representing the distribution of cell width (in µm) across at least 100 wild-type, 2xFYVE::GFP- and GFP::P4C-expressing cells selected in an unbiased fashion. A statistically significant (two-sided t-test assuming unequal variances, p<0.05) increase in median cell width with respect to non-transgenic cells is detected for 2xFYVE::GFP-but not GFP::P4C-expressing cells. Asterisks indicate statistical significance. n.s.: not significant. (G) Box-plot representing the distribution of measured Dextran-R signal intensity across at least 100 wild-type, 2xFYVE::GFP- and GFP::P4C-expressing cells selected in an unbiased fashion. A statistically significant (two-sided t-test assuming unequal variances, p<0.05) decrease in Dextran-R signal intensity, normalized to wild-type cells (100%), is detected for 2xFYVE::GFP-but not for GFP::P4C-expressing cells. Asterisks indicate statistical significance. n.s.: not significant. (H) A statistically significant (p<0.5) linear correlation exists between Dextran-R signal (*x*-axis, intensity_red channel [%]) and 2xFYVE::GFP expression (*y*-axis, intensity_green channel [%]) measured across 100 cells. (I) The apparent linear correlation between GFP::P4C expression (*y*-axis, intensity_green channel [%]) and Dextran-R signal (x-axis, intensity_red channel [%]) is not statistically significant (p<0.5). (J) Wide-field microscopy-based immunofluorescence analysis of the impact of Neomycin treatment on Dextran-R uptake to deplete PI(4,5)P2 and PI(3,4,5)P_3_ binding sites in non-transgenic wild-type cells. With respect to non-treated cells (WT; left panel), Dextran-R signal at PVs is visibly impacted in neomycin-treated cells (WT_Neo; right panel). Scale bars: 10 µm for full wide-field image, 1 µm for a single cell. (K) Box-plot representing the distribution of measured Dextran-R signal intensity across 100 wild-type cells, either untreated (WT) or treated with neomycin (WT_Neo). Neomycin treatment causes a statistically significant (two-sided t-test assuming unequal variances, p<0.05) decrease in Dextran-R signal. Scale bars: wide-field: 10 µm; single cells:: 1 µm. For all images, nuclei are labelled with DAPI (blue). Insets: DIC images.

The cationic antibiotic neomycin binds tightly to the headgroup of phosphoinositides with a marked preference for PI(4,5)P_2_ and PI(3,4,5)P_3_ [40, 41]. As a means to saturate PI(4,5)P_2_ and PI(3,4,5)P_3_ binding in *Giardia* trophozoites, we tested its effect on fluid-phase uptake by treating wild-type trophozoites with 2mM neomycin followed by uptake of Dextran-R. Quantitative light microscopy image analysis revealed a significantly lower level of Dextran-R in treated trophozoites (p<0.05). (Fig 3J, K). Taken together, the data indicate that saturation of PI(3)P, PI(4,5)P_2_, and PI(3,4,5)P_3_, but not PI(4)P binding significantly impacts fluid-phase endocytosis through *G. lamblia* PVs.

### Functional characterization of GlPXD1-4 and 6, GlFYVE and GlNECAP1

Manipulation of PIP residue homeostasis elicited PV-dependent fluid-phase uptake phenotypes. We hypothesized that changing expression levels of giardial PIP-binding proteins previously identified in clathrin interactomes would elicit aberrant uptake phenotypes in *Giardia* trophozoites. In addition we explored the functional boundaries of each PIP-binding module by defining their protein interactomes. To test this, we used the previously-generated epitope-tagged reporter lines for full-length *Gl*PXD1-4 and 6, *Gl*FYVE and *Gl*NECAP1 (Fig 1C) for assessing the effects of ectopic expression on fluid-phase uptake phenotypes. Furthermore, we used the same lines as baits in antibody-based affinity co-immunoprecipitation (co-IP) and identification of reporter-associated protein complexes. Further investigation of *Gl*PXD5 was abandoned at this stage due to its intractably low levels of expression.

#### The extended interactomes of GlPXD1, GlPXD4 and GlPXD6

Epitope-tagged, full-length *Gl*PXD1 is a validated *Gl*CHC interaction partner; its extended interactome confirms association to all core clathrin assembly components (*Gl*CHC, *Gl*CLC, *Gl*DRP, *Gl*AP2) (Fig 4 and Table S1) [19]. A weaker interaction with *Gl*PXD2 was also found. The *Gl*PXD4 interactome includes *Gl*CHC and *Gl*DRP and, uniquely for the *Gl*PXD protein family, a previously confirmed interaction with *Gl*PXD2 [19] albeit detected at lower stringencies (95_2_95, 2 hits) (Fig 4; Table S4). A putative SNARE protein GL50803_5785, previously identified in the *Gl*Tom40 interactome [42], was detected at lower stringencies (95_2_95, 2 hits). Similar to *Gl*PXD1, *Gl*PXD6 showed strong interaction with the β subunit of *Gl*AP2 and *Gl*CHC (Fig 4), although the reverse interaction was not detected in the previously-published clathrin-centered interactome [19]. Using lower stringency parameters (95_2_50, 3 hits), revealed interaction with *Gl*FYVE, *Gl*PXD3 and *Gl*DRP (Fig 4; Table S). The *Gl*PXD6 interactome includes *Gl*16717, a protein of unknown function predicted to carry a StAR-related lipid-transfer domain (Steroidogenic Acute Regulatory protein, START) domain [43]. Ectopic expression of epitope-tagged *Gl*PXD1, 4 and 6 elicited no discernible PV-related phenotypes.

**Figure 4:**
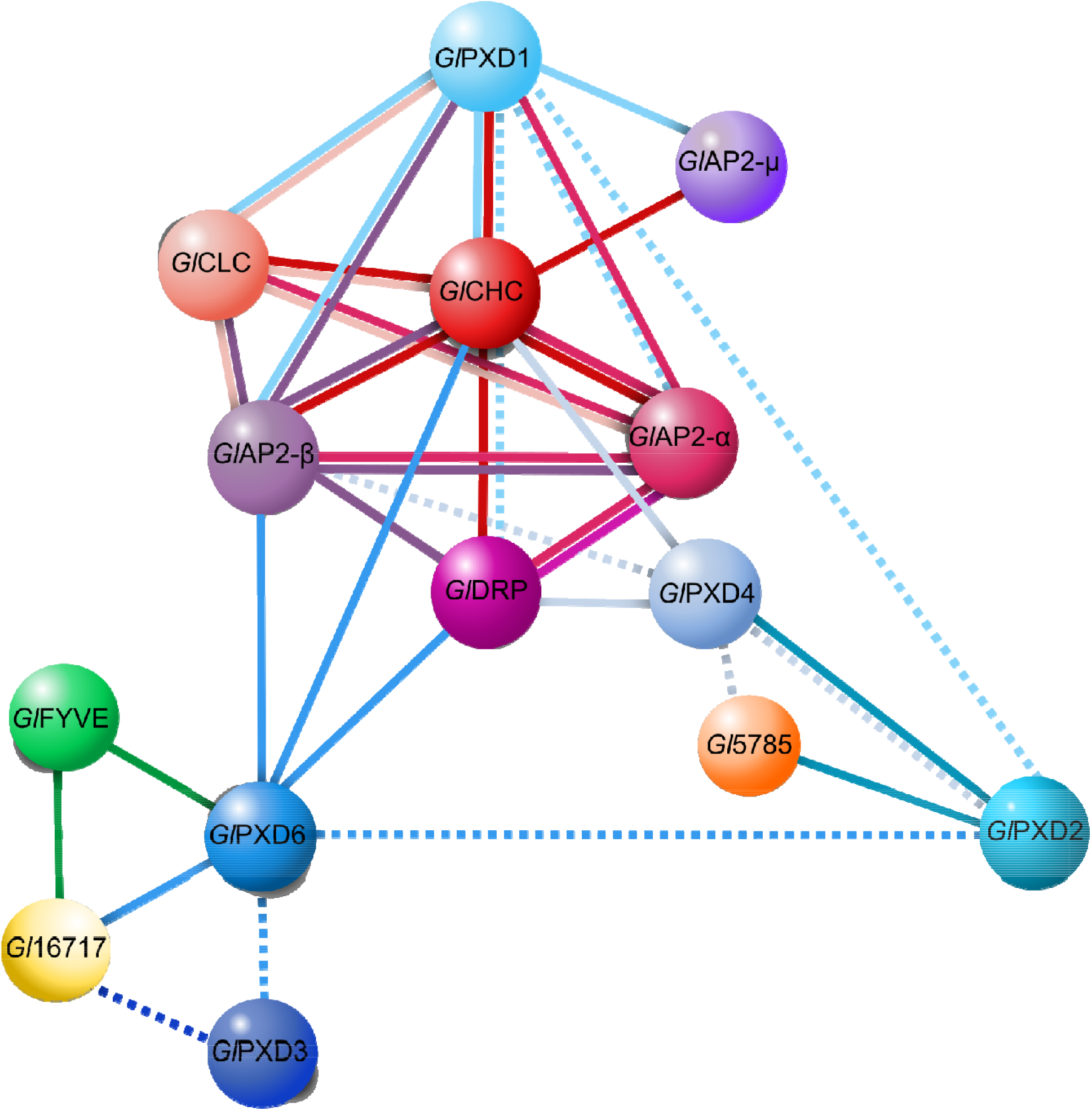
The extended interactomes of *Gl*PXD1, *Gl*PXD4 and *Gl*PXD6. Curated core interactomes for *Gl*PXD1, *Gl*PXD4 and *Gl*PXD6. All three epitope-tagged variants used as affinity handles in co-immunoprecipitation experiments identify *Gl*CHC as a strong interaction partner for *Gl*PXD1, 4 and 6. *Gl*PXD1 and 4 further interact with other known clathrin assembly components such as *Gl*CLC, *Gl*AP2 subunits α, β and µ, and *Gl*DRP. *Gl*PXD2, albeit at low stringency, is the only other PXD protein found in all three interactomes. The *Gl*PXD4 interactome includes a putative SNARE protein (5785; [42] while *Gl*PXD6 as an affinity handle pulled down another PIP residue binder, *Gl*FYVE, known to be associated to clathrin assemblies in *G. lamblia* [19]. Solid lines: interactions detected at high stringency. Dashed lines: interactions detected at low stringency. Yellow partners are currently annotated on GDB as “hypothetical protein” i.e. proteins of unknown function.

#### Ectopic expression of tagged GlPXD2 severely perturbs PV organisation

Mining the *Gl*PXD2 protein interactome dataset with high stringency parameters confirmed interactions with *Gl*CHC, *Gl*AP2 and *Gl*PXD4 (Fig 5A; Table S2). Furthermore, we identified three predicted SNARE proteins: GL5785, GL50803_14469 (*Gl*14469; at lower stringencies 95_2_50, 9 hits) and GL50803_10013 (*Gl*10013; Fig 5A) [44]. The SNARE GL5785 was detected also in the interactomes of *Gl*PXD4 *Gl*TOM40 [42]. *Gl*NECAP1 was also identified as a *Gl*PXD2 interacting partner, albeit only by applying low stringency parameters (95_2_50, represented by a dashed line, Fig 5A).

**Figure 5:**
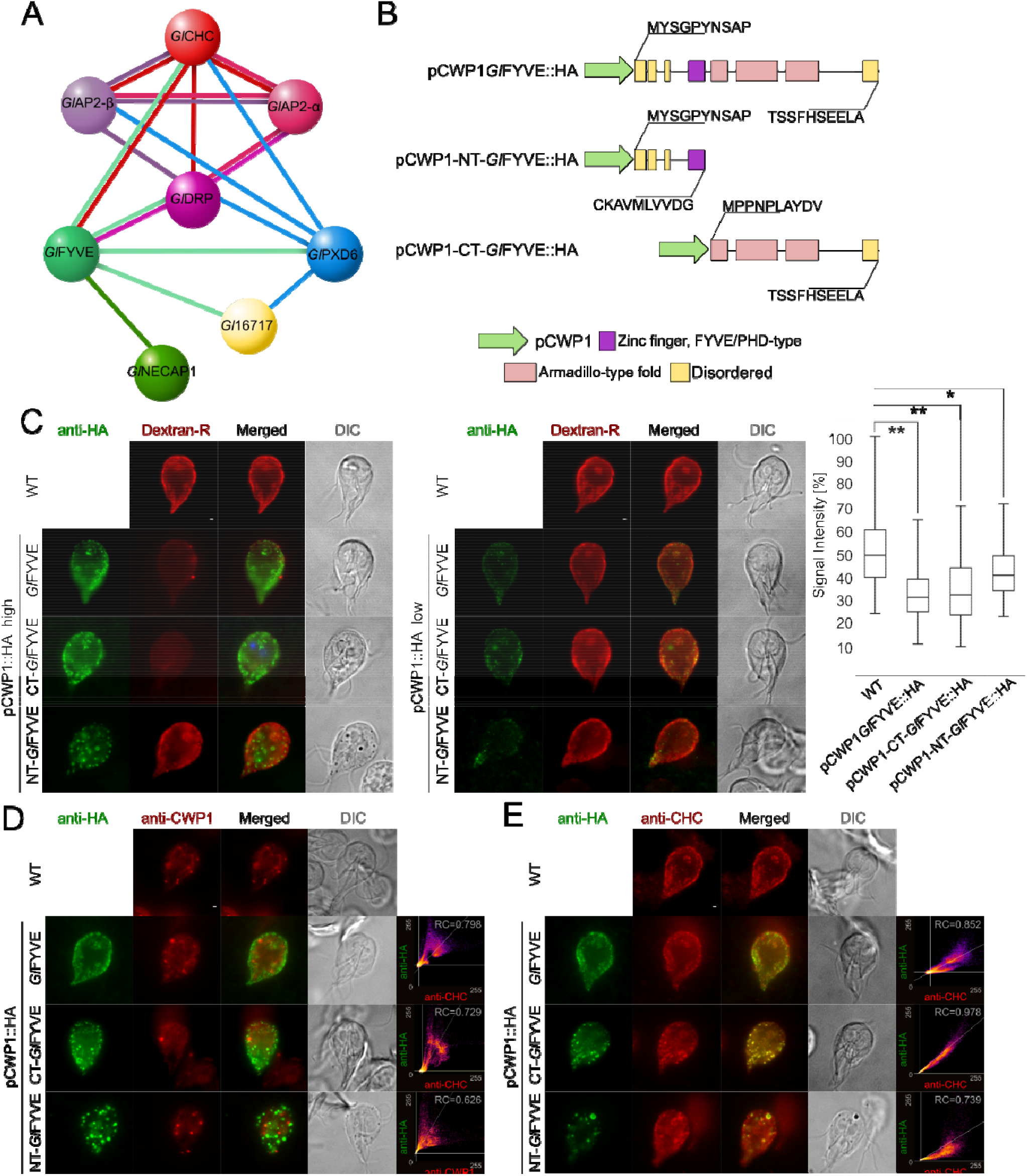
The extended interactome of *Gl*PXD2 and the impact of *Gl*PXD2 ectopic expression on PV morphology. (A) The extended interactome analysis of epitope-tagged *Gl*FYVE confirms confirms tight association to *Gl*CHC, *Gl*DRP and *Gl*PXD6. *Gl*NECAP1 as an alternative PIP-binding module was also detected. (B) C-terminally epitope-tagged full-length (top; pCWP1-*Gl*FYVE::HA), C-terminal truncated (middle; pCWP1-NT-*Gl*FYVE::HA, residues 1-300) and N-terminal truncated (bottom; pCWP1-CT-*Gl*FYVE::HA, 301-990 residues) constructs were generated for regulated expression and phenotype testing. (C) Confocal imaging and immunofluorescence analysis of non-transgenic wild-type cells and in cells overexpressing constructs *Gl*FYVE::HA, NT-*Gl*FYVE::HA or pCWP1-CT-*Gl*FYVE::HA (anti-HA) shows statistically significant (two-sided t-test assuming unequal variances, p<0.05) differences in their ability to take up Dextran-R. Cells overexpressing construct pCWP1-NT-*Gl*FYVE::HA present additional membrane-bound structures that are not detected in other lines and do not associate with Dextran-R labelling. Asterisks indicate statistical significance: * p<0.05; ** p<0.005. n.s.: not significant. DIC: differential interference contrast. Scale bars: 1 µm. (D) Confocal imaging and immunofluorescence analysis of non-transgenic wild-type cells and cells overexpressing constructs *Gl*FYVE::HA, NT-*Gl*FYVE::HA or pCWP1-CT-*Gl*FYVE::HA (anti-HA) using anti-CWP1-TxRed antibody (anti-CWP1) shows that the membrane compartments found in NT-*Gl*FYVE::HA-expressing cells are not related to ESVs. Scale bars: 1 µm. (E) Antibody-based detection and immunofluorescence analysis of *Gl*CHC deposition (anti-CHC) in non-transgenic wild-type cells and in cells overexpressing constructs *Gl*FYVE::HA, NT-*Gl*FYVE::HA or pCWP1-CT-*Gl*FYVE::HA (anti-HA) detects a significant degree of *Gl*CHC association to the CT-*Gl*FYVE::HA variant, with only partial association to NT-*Gl*FYVE::HA and *Gl*FYVE::HA constructs. Scale bars: 1 µm.

In contrast to ectopic expression of tagged *Gl*PXD1, 4, and 6, expression of an epitope-tagged reporter HA::*Gl*PXD2 elicited a distinct phenotype. In contrast to non-transgenic wild-type cells (Fig 5C) and weakly-expressing HA::*Gl*PXD2 cells (Figs 5D and E upper panels), gated STED imaging of trophozoites strongly expressing HA::*Gl*PXD2 showed large membranous clusters which also accumulated Dextran-R (Fig 5D) and were bound by both anti-*Gl*CHC (Fig 5E) and anti-PI(3)P (Fig 5F) antibodies. Transmission electron microscopy (tEM) analysis confirmed the presence of randomly distributed peripheral PV clusters in cells expressing HA::*Gl*PXD2 (Fig 5G; left panel) which were not present in representative wild-type control cells (Fig 5G; right panel).

#### The GlPXD3 interactome is connected to clathrin assemblies and includes a novel dynamin-like protein

*Gl*DRP, *Gl*CHC, and *Gl*AP2 (α/β subunits) were detected in the *Gl*PXD3 interactome, thereby establishing the association of this PX domain protein with clathrin assembly structures at the PV/PM interface (Fig 6A; Table S3). A pseudokinase (*Gl*15411 [45]) previously identified in *Gl*CHC assemblies was also found in the *Gl*PXD3 interactome (Fig 6A; [19]). Furthermore, the *Gl*PXD3 and *Gl*15411 interactomes share proteins GL50803_16811 (*Gl*16811) tentatively annotated as a ZipA protein in GDB, and proteins GL50803_87677 (*Gl*87677) and GL50803_17060 (*Gl*17060), annotated as a NEK kinase and an ankyrin-domain carrying protein, respectively (Fig 6A). Unique interaction partners for *Gl*PXD3 include the SNARE protein *Gl*7309 [44] and *Gl*NSF (GL50803_114776) [46]. In addition, protein GL50803_103709 carrying a predicted N-terminal BRO domain and protein GL50803_9605 were identified as unique *Gl*PXD3 interaction partners (Fig 6A). Furthermore, the StAR-related lipid-transfer protein *Gl*16717, already found in the *Gl*PXD6 interactome, was also found to be a low-stringency interaction partner for *Gl*PXD3 and *Gl*15411, thereby connecting the *Gl*PXD3 and *Gl*PXD6-*Gl*FYVE circuits. IFA analysis for *Gl*15411, *Gl*16811, *Gl*103709, and *Gl*7309 shows PV-associated labelling profiles for all corresponding epitope-tagged reporters whereas *Gl*NSF presents a diffused localization pattern and *Gl*9605 shows a diffused yet punctate deposition pattern (Fig 6B).

**Figure 6:**
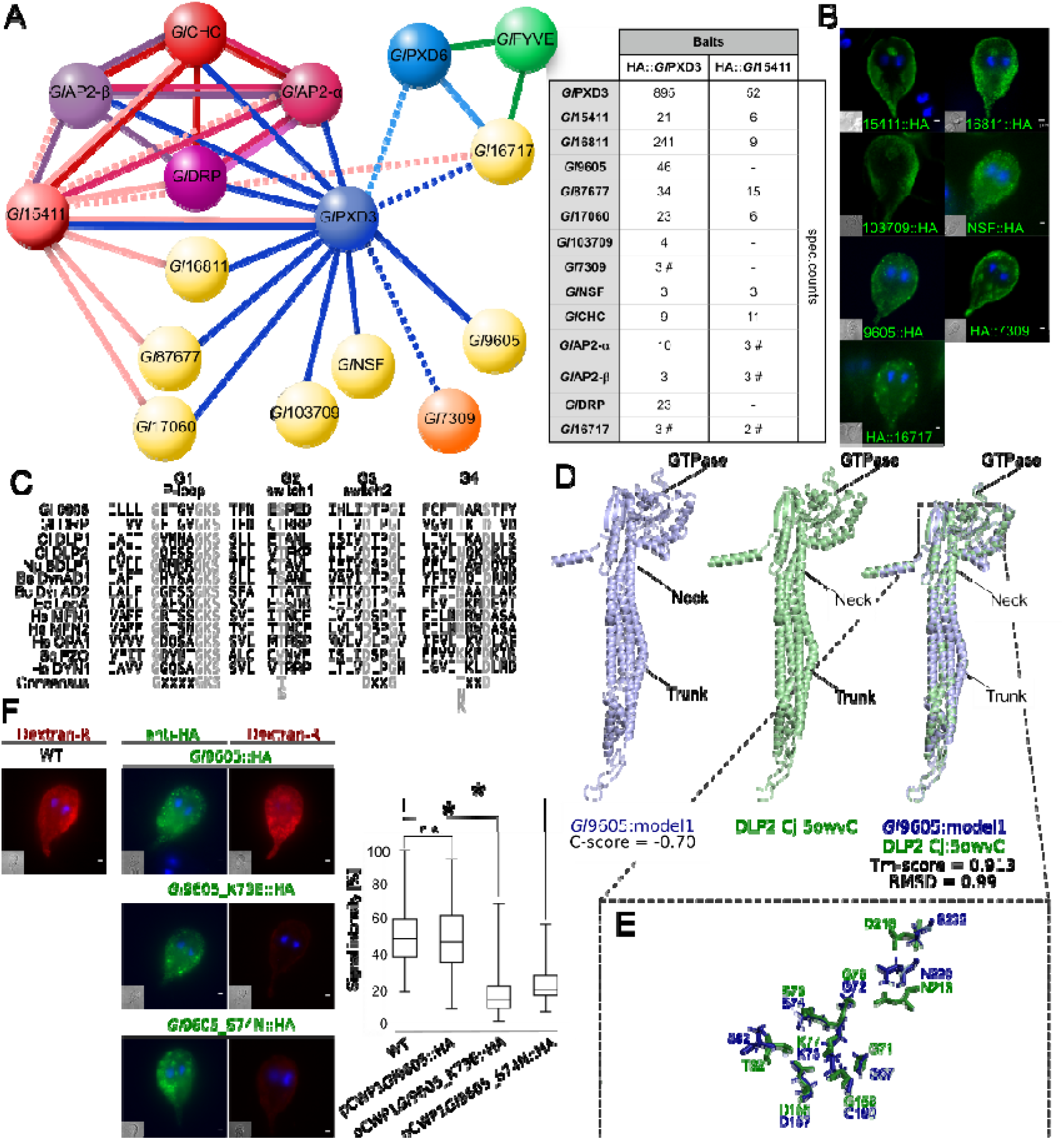
The extended *Gl*PXD3 interactome includes a novel dynamin-like protein in *G. lamblia*. (A) Left panel: Analysis of the extended *Gl*PXD3 interactome using an epitope-tagged variant as affinity handle reveals robust interactions with clathrin assembly components *Gl*CHC, α and β *Gl*AP2 subunits, and *Gl*DRP. Predicted inactive inactive NEK kinase 15411 [45] is similarly associated to clathrin assemblies [19] and further shares proteins *Gl*16811, *Gl*87677 and *Gl*17060 as interaction partners with *Gl*PXD3. Predicted SNARE protein *Gl*7309, *Gl*NSF (GL50803_1154776) and proteins *Gl*103709 and *Gl*9605 are unique *Gl*PXD3 interaction partners. The *Gl*PXD3 interactome is connected to the *Gl*PXD6 circuit both directly and through *Gl*16717. Solid lines: interactions detected at high stringency. Dashed lines: interactions detected at low stringency. Yellow partners are currently annotated on GDB as “hypothetical protein” i.e. proteins of unknown function. Right panel: Total spectral counts as a measure of relative abundance for interaction candidates in the interactomes of *Gl*PXD3 and *Gl*15411 epitope-tagged variants. Hashtag: detection at low stringency (95_2_50) and visualised with a dashed line in the interactome. (B) Light-microscopy-based immunofluorescence analysis of representative transgenic trophozoites expressing epitope-tagged reporter variants for proteins *Gl*15411, *Gl*16811, *Gl*103709, *Gl*NSF, *Gl*9605, *Gl*7309 and *Gl*16717. Cells were imaged at maximum width, where nuclei and the bare-zone are at maximum diameter. Nuclei are labelled with DAPI (blue). Insets: DIC images. Scale bars: 1 µm (C) MSA analysis G1-Ploop, G2 switch1, G3 switch 2 and G4 regions of the conserved GTPase domains of *Gl*9605, *Gl*DRP, *Campylobacter jejuni* DLP1 (Uniprot accession CJ0411) and DLP2 (CJ0412), *Nostoc punctiforme* BDLP1 (B2IZD3), *Bacillus subtilis* DynAD1 (P54159), *Bacillus cereus* DynAD2 (CUB17917), and *Escherichia coli* LeoA (E3PN25) bacterial dynamin-like proteins (BDLPs), *Homo sapiens* MFN1 (Q8IWA4), MFN2 (O95140), OPA1 (O60313) and DYN1 (Q05193), and *Saccharomyces cerevisiae* Fzo1p (P38297). Conserved positions are highlighted in grey. (D) I-TASSER *de novo* predicted 3D structure for *Gl*9605 (blue) and its closest known structural homologue, *C. jejuni* DLP2 (5owvC; green) indicating the GTPAse, neck and trunk regions that characterize BDLPs. (E) A close-up view of the overlapping structures in the GTPase domains of *Gl*9605 (blue) and *C. jejuni* DLP2 (5owvC; green) marking specific residues important for GTP binding and catalytic activity. (F) Quantitative microscopy-based immunofluorescence analysis of Dextran-R signal in cells overexpressing either a non-mutated full length epitope-tagged *Gl*9605 or mutated *Gl*9605 K73E and S74N variants. In contrast to non-transgenic wild-type controls and *Gl*9605::HA overexpressing cells, expression of *Gl*9605 K73E and S74N variants inhibited Dextran-TxR uptake in a statistically significant fashion (box-plot). Asterisks indicate statistical significance. n.s.: not significant.

Protein *Gl*9605, the sixth most abundant hit in the *Gl*PXD3 interactome (Fig 6A), and currently annotated as having an unknown function, was identified as a highly-diverged dynamin-like protein (Fig 6C). In support of this, the predicted GTPase domain in *Gl*9605 contains signature motifs in the P-loop (G1), switch 1 (G2) and switch 2 (G3) regions [47–49]. Conserved motifs in the G4 region are only partially maintained (Fig 6D). To test residue conservation on a structural level, *Gl*9605 was subjected to *ab initio* modelling using I-TASSER and the resulting tertiary structure was superimposed on that of a dynamin-like 2 (DLP2 Cj:5ovW) [50], *Gl*9605’s closest structural homologue (Fig 6D). A structural overlap TM-score of 0.913 suggests an almostperfect structural match, with clear chemical and positional conservation of key residues involved in GTPase activity (Fig 6E). We sought to elicit a dominant-negative phenotype by engineering *Gl*9605 K73E and S74N mutants [51]. In contrast to either wild-type cells or cells overexpressing a wild-type epitope-tagged *Gl*9605 control, expression of *Gl*9605 K73E and S74N mutant reporters inhibited fluid-phase uptake of Dextran-R in a statistically significant manner (p<0.05; Fig 6F).

#### Regulated ectopic expression of GlFYVE variants inhibits fluid-phase uptake and induces the emergence of novel membrane-bound compartments

*Gl*FYVE is a confirmed interactor of clathrin assemblies (Zumthor et al., 2016) through specific association to *Gl*CHC and *Gl*DRP (Fig 7A) Table S6). *Gl*FYVE’s extended interactome includes *Gl*PXD6, *Gl*NECAP1 and protein GL50803_16717 (*Gl*16717). The latter was also found in the *Gl*PXD6 interactome (Fig 4) and partially localizes to PVs as an epitope-tagged reporter (Fig 7A).

**Figure 7:**
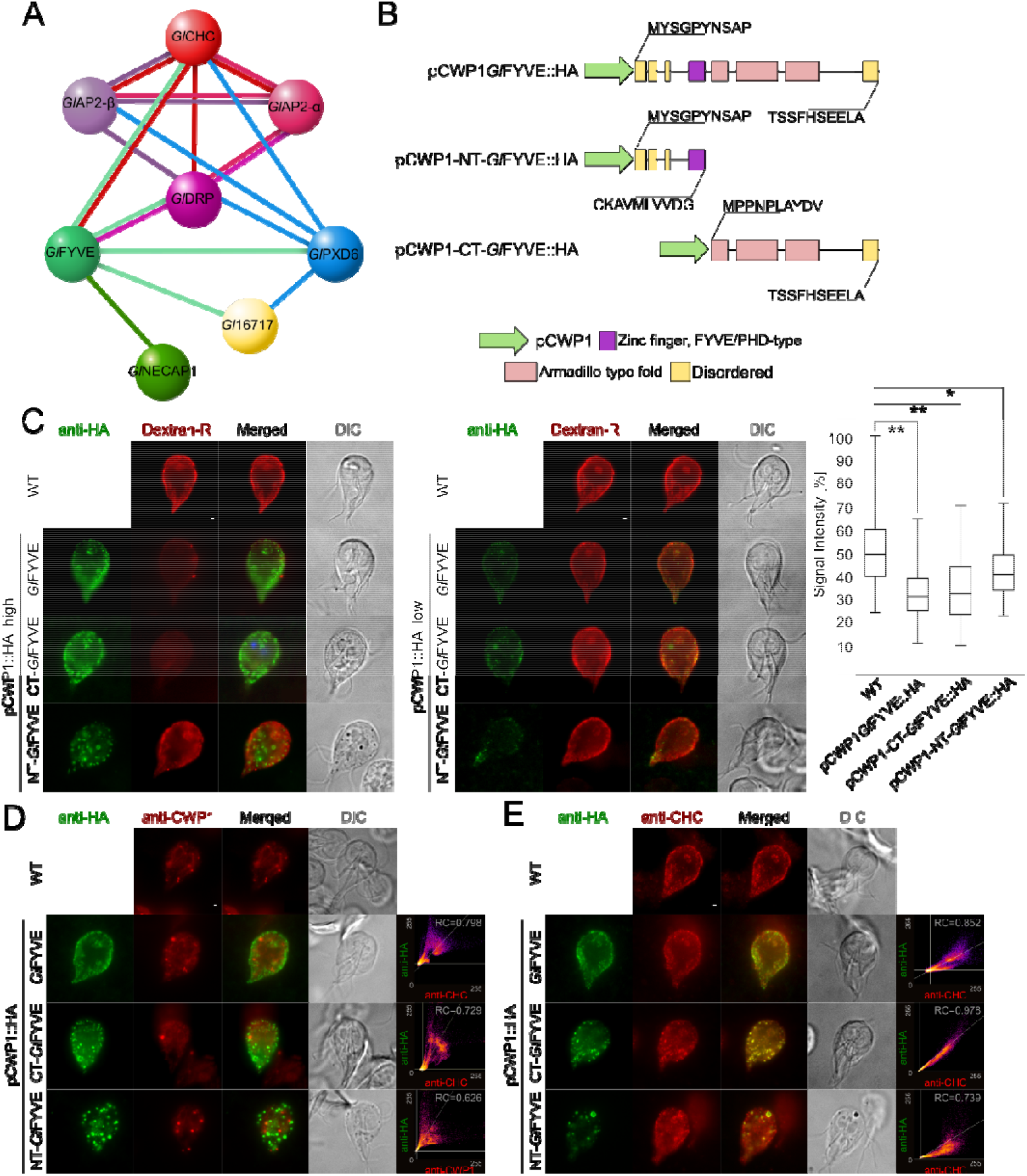
Regulated ectopic expression of *Gl*FYVE variants inhibits fluid-phase uptake and induces novel membrane-bound compartments. (A) The extended interactome analysis of epitope-tagged *Gl*FYVE confirms confirms tight association to *Gl*CHC, *Gl*DRP and *Gl*PXD6. *Gl*NECAP1 as an alternative PIP-binding module was also detected. (B) C-terminally epitope-tagged full-length (top; pCWP1-*Gl*FYVE::HA), C-terminal truncated (middle; pCWP1-NT-*Gl*FYVE::HA, residues 1-300) and N-terminal truncated (bottom; pCWP1-CT-*Gl*FYVE::HA, 301-990 residues) constructs were generated for regulated expression and phenotype testing. (C) Confocal imaging and immunofluorescence analysis of non-transgenic wild-type cells and in cells overexpressing constructs *Gl*FYVE::HA, NT-*Gl*FYVE::HA or pCWP1-CT-*Gl*FYVE::HA (anti-HA) shows statistically significant (two-sided t-test assuming unequal variances, p<0.05) differences in their ability to take up Dextran-R. Cells overexpressing construct pCWP1-NT-*Gl*FYVE::HA present additional membrane-bound structures that are not detected in other lines and do not associate with Dextran-R labelling. Asterisks indicate statistical significance: * p<0.05; ** p<0.005. n.s.: not significant. DIC: differential interference contrast. Scale bars: 1 µm. (D) Confocal imaging and immunofluorescence analysis of non-transgenic wild-type cells and cells overexpressing constructs *Gl*FYVE::HA, NT-*Gl*FYVE::HA or pCWP1-CT-*Gl*FYVE::HA (anti-HA) using anti-CWP1-TxRed antibody (anti-CWP1) shows that the membrane compartments found in NT-*Gl*FYVE::HA-expressing cells are not related to ESVs. Scale bars: 1 µm. (E) Antibody-based detection and immunofluorescence analysis of *Gl*CHC deposition (anti-CHC) in non-transgenic wild-type cells and in cells overexpressing constructs *Gl*FYVE::HA, NT-*Gl*FYVE::HA or pCWP1-CT-*Gl*FYVE::HA (anti-HA) detects a significant degree of *Gl*CHC association to the CT-*Gl*FYVE::HA variant, with only partial association to NT-*Gl*FYVE::HA and *Gl*FYVE::HA constructs. Scale bars: 1 µm.

To characterize the function of *Gl*FYVE and to test whether a dominant-negative effect on uptake could be elicited, we performed a deletion analysis by generating epitope-tagged C-terminal (pCWP1-NT-*Gl*FYVE::HA) and N-terminal (pCWP1-CT-*Gl*FYVE::HA) truncation constructs, consisting of either the disordered region followed by the FYVE domain (Fig 7B), residues 1-300) or the armadillo repeat-rich (ARM repeats) domains (Fig 7B), residues 301-990), respectively. Expression of both constructs is regulated by an inducible promoter which is de-repressed during induction of encystation [52]. After a short (6hrs) induction pulse, transfected cells were subjected to Dextran-R uptake. Both in cells expressing the full-length pCWP1-*Gl*FYVE::HA and truncated variants the amount of Dextran-R accumulated in PVs was significantly (p<0,05) lower (Fig 7C, box plot). Furthermore, IFA analysis of pCWP1-NT-*Gl*FYVE::HA cells revealed the presence of membrane-bound compartments which overlapped neither with Dextran-R-labelled PVs (Fig 7C) nor with encystation specific vesicles (ESVs) labeled with the anti-CWP1 antibody (Fig 7D). In contrast, CT-*Gl*FYVE::HA and full length *Gl*FYVE::HA localized predominantly to PVs as (Fig 7C and D). The subcellular localization of *Gl*CHC in these lines and in a wild-type control overlapped with the truncated CT-*Gl*FYVE::HA variant, but only partially with NT-*Gl*FYVE::HA and *Gl*FYVE::HA (Fig 7E).

#### Ectopic expression of GlNECAP1 significantly impairs fluorescent Dextran uptake

Co-IP using epitope-tagged *Gl*NECAP1 (Figs 1B and C) confirmed interaction with clathrin assembly components *Gl*AP2-β, µ and α subunits, *Gl*CHC and *Gl*DRP (Table S7). Interaction with *Gl*FYVE (Fig 7A) and, at lower stringency also for *Gl*PXD2 (Fig 5A) could be confirmed (Fig 8A). Three putative conserved AP2-interacting motifs were identified using multi-sequence alignment; the high affinity WxxF motif at the N-terminus, two residues being invariant throughout evolution, K147 and G149, and AP2-beta linker interacting residues binding sites (Fig 8B) [53]. *De novo* 3D modelling confirms overall structural conservation of all key residues in *Gl*NECAP1 (Fig 8B) when compared with mammalian NECAP1 (Fig 8C). Furthermore, the interacting interface of NECAP1 with the β-linker region of AP2 was also identified in the structural model for *Gl*NECAP1 (Fig 8C).

**Figure 8:**
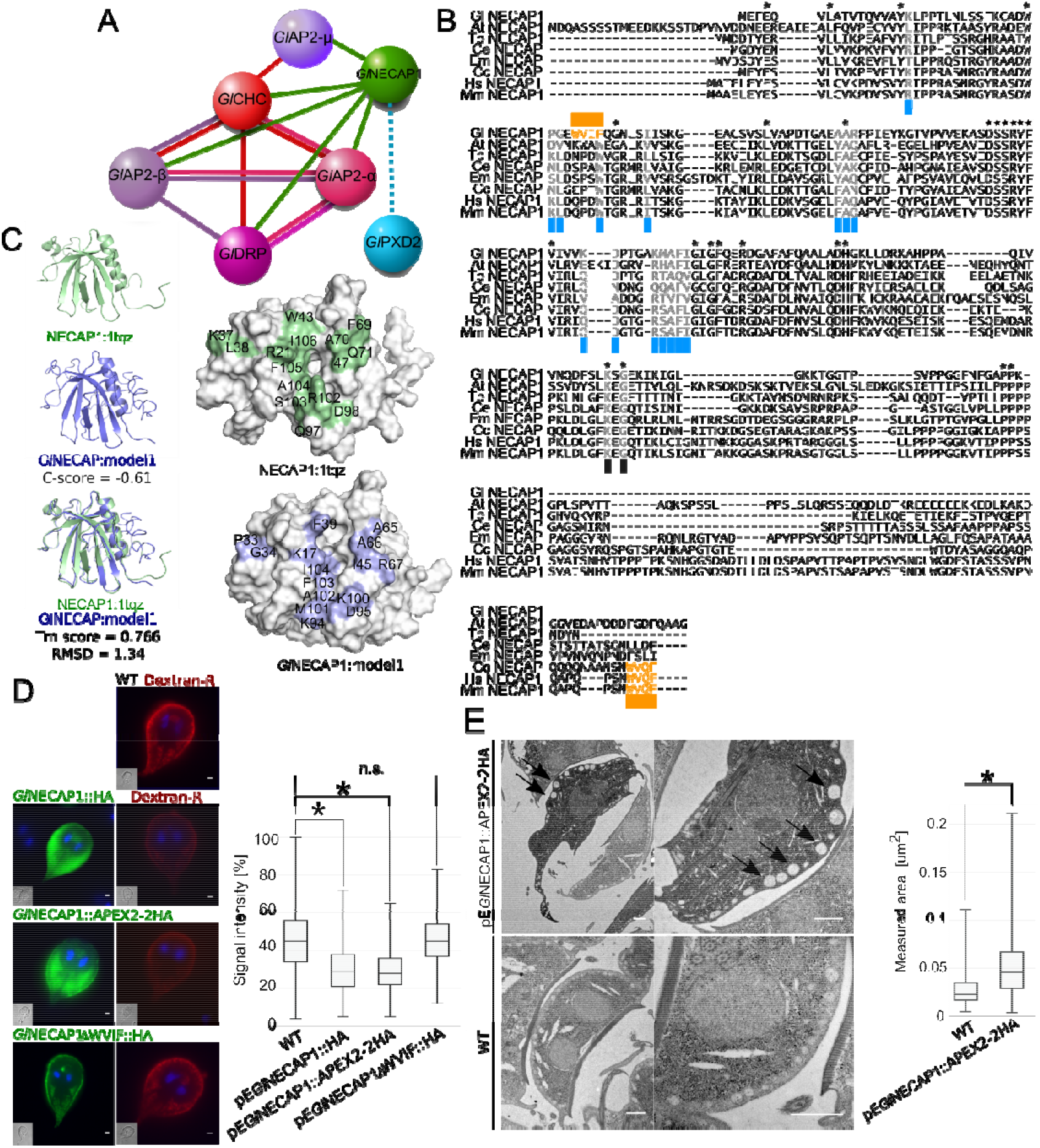
PV morphology and functionality phenotypes caused by *Gl*NECAP1 ectopic expression. (A) A *Gl*NECAP1-centered interactome highlights association to clathrin assembly components and to additional PIP-residue binders *Gl*FYVE and *Gl*PXD2. (B) Multiple sequence alignment analysis of *Gl*NECAP1 and NECAP1 orthologues from *Arabidopsis thaliana* (Uniprot accession Q84WV7), *Trichinella pseudospiralis* (A0A0V1JQ20), *Caenorhabditis elegans* (Q9N489), *Echinococcus multilocularis* (A0A087VZS0), *Ceratitis capitata* (W8CD89), *Homo sapiens* (Q8NC96) and *Mus musculus* (Q9CR95) identifies conserved motifs and residues for interaction with AP2. *Gl*NECAP1 presents partial conservation, with a WXXF motif (orange) shifted to the N-terminus with respect to other orthologues. (C) *Ab initio* template-based 3D modelling of *G. lamblia* and *H. sapiens* NECAP1 (1tqz) homologues predicts similar structures, with conservation of key residues involved in the interaction between NECAP1 proteins and AP2 complexes (shaded in blue and green). (D) Wild-type non-transgenic control cells (WT) and cells overexpressing either epitope-tagged *Gl*NECAP1 reporters *Gl*NECAP1::HA, *Gl*NECAP1::APEX2-2HA or the ΔWVIF deletion construct *Gl*NECAP1∆WVIF::HA (green) were tested for Dextran-R (red). Dextran-R signal intensity was significantly (p<0.05) decreased in *Gl*NECAP1::HA- and *Gl*NECAP1::APEX2-2HA-expressing cells compared to wild-type controls and *Gl*NECAP1∆WVIF::HA-expressing cells (box-plot). (E) Quantitative tEM analysis of *Gl*NECAP1::APEX2-2HA-expressing cells (upper panels) and wild-type non-transgenic cells (WT; lower panels) shows visibly enlarged PVs in *Gl*NECAP1::APEX2-2HA-expressing cells, with a statistically significant (p<0,05) increase in median PV area (in µm^2^; box-plot).

To test whether expression of a *Gl*NECAP1 variant lacking the putative high-affinity motif WVIF could elicit a dominant-negative uptake effect, a deletion construct *Gl*NECAP1∆WVIF::HA lacking this motif (Fig 8B) for conditional expression in induced trophozoites. Accumulation of Dextran-R into PVs detected by microscopy was significantly lower (p<0.05) in induced cells expressing *Gl*NECAP1::HA or an APEX- and epitope-tagged variant *Gl*NECAP1::APEX2-2HA compared with wild type controls (Fig 8D, box plot). Conversely, inducible expression of a deletion construct *Gl*NECAP1∆WVIF::HA (Fig 8D, *Gl*NECAP1∆WVIF::HA) had no discernible effect on accumulation of Dextran-R in PVs (Fig 8D, box plot). Inducible expression of the genetically encoded enzymatic reporter [54, 55] *Gl*NECAP1::APEX2-2HA showed subcellular distribution of APEX-derived deposits around significantly enlarged PVs in tEM compared to wild type controls (Fig 8E; Fig S4).

### GlPXD3 associates specifically with PVs as membrane coat

Co-localization studies with Dextran-OG and ectopically expressed HA:: *Gl*PXD3 show apparent coating of the entire PV membrane on the cytoplasmic side by the reporter construct (Fig 9A; Fig S2C) This provided us with an opportunity to generate measurements of PV organelles in optical sections using 3D STED microscopy followed by reconstruction and rendering with IMARIS. Rendered images show hive-like *Gl*PXD3-labelled structures predominantly in the cortical area of the cell underneath the PM that clearly surround the entire PV membrane (Fig 9B). The major and minor principal axes of these structures measured 437 +/− 93 nm and 271 +/− 60 nm. Consistent with the subcellular localization of this marker on the cytoplasmic side of PV membranes, these values were significantly higher (p≤ 0.05) than those obtained from PVs labeled with Dextran-OG (371 +-79 nm and 221 +/− 49 nm) (Fig 9C, D). Signal overlap of epitope-tagged *Gl*PXD3 with endogenous GlCHC as a marker for the PM-PV interface [19] in fluorescence microscopy is low. The image data indicate that both labels have distinct distributions but may spatially overlap at focal clathrin assemblies in small areas at the PV-PM interface (Fig 9E). Similarly, labelling for both PI(3)P and a reporter *Gl*PXD3 variant showed minimal signal overlap (Fig 9F), despite the strong affinity of the latter for this lipid in in vitro lipid-array binding experiments (Fig 2A and B).

**Figure 9:**
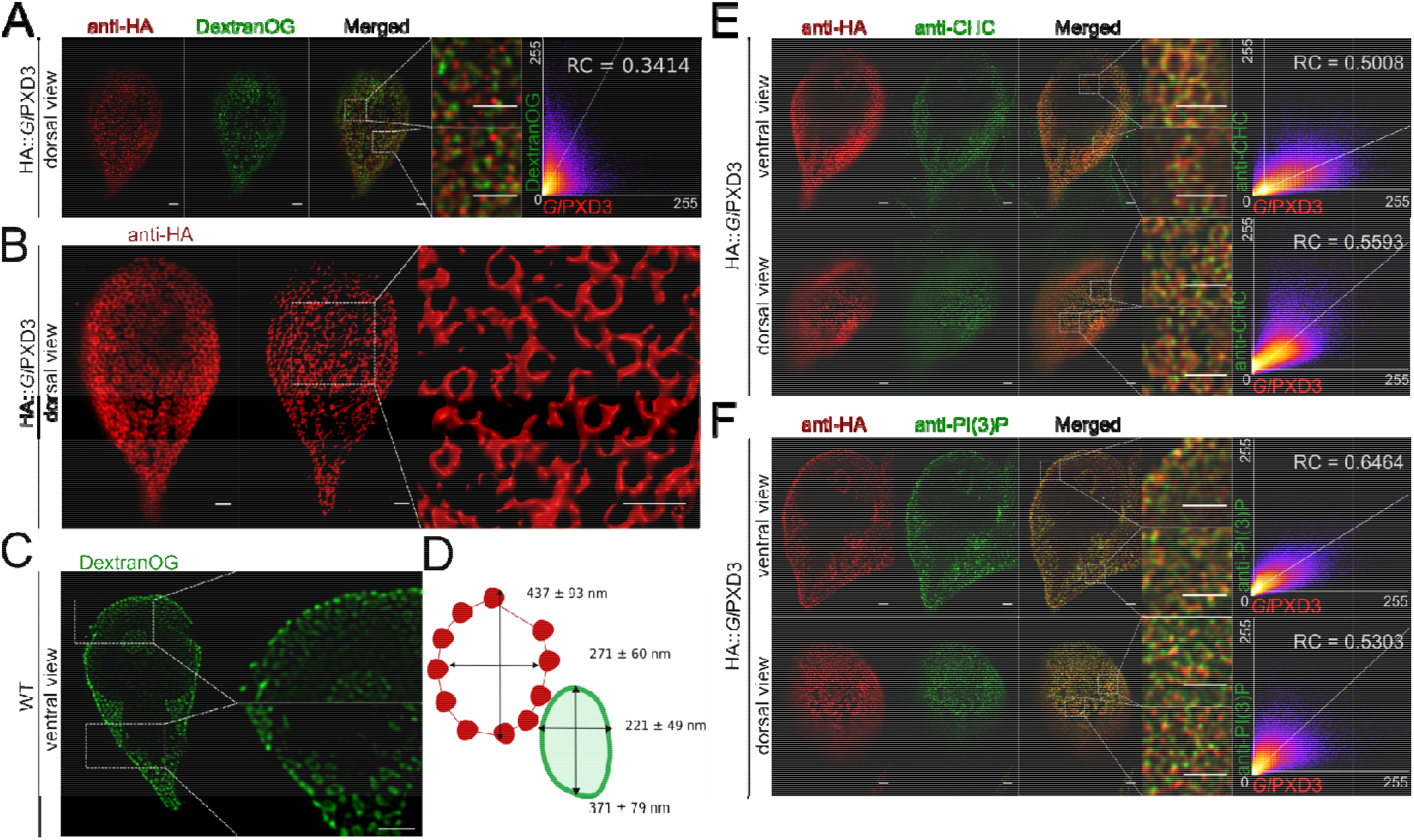
*Gl*PXD3 membrane coats as a tool to probe PV size and organization. (A) A dorsal view of representative cells expressing an epitope-tagged *Gl*PXD3 reporter (red) and co-labelled for Dextran-OG (green). STED confocal imaging followed by signal overlap analysis (scatter plot) shows proximal yet distinct deposition patterns, with *Gl*PXD3 reporters closely associated to Dextran-OG-illuminated PVs. Scale bars: whole cell 1 µm; close-ups 1 µm. (B) 3D STED microscopy (left panel) followed by reconstruction using IMARIS (middle panel) of a representative cell expressing an epitope-tagged *Gl*PXD3 reporter reveals fenestrated *Gl*PXD3-delimited areas distributed under the PM and throughout the whole cell (close-up view of inset in the right panel). Scale bars: whole cell 1 µm; close-ups 1 µm. (C) STED microscopy analysis of PVs in a representative non-transgenic wild-type cell labelled with Dextran-OG. Scale bars: whole cell 1 µm; close-ups 1 µm. (D) Average length of the major and minor principle axes of *Gl*PXD3-delimited fenestrated structures (in red) and Dextran-labelled PV organelles in wild-type non-transgenic cells (in green) measured across at least 100 structures/organelles. (E) STED confocal microscopy analysis of ventral and dorsal views of a representative cell expressing an epitope-tagged *Gl*PXD3 reporter (anti-HA) and co-labelled for *Gl*CHC (anti-CHC) shows how fenestrated *Gl*PXD3-delimited structures are decorated with *Gl*CHC foci. Scatter plots are included for signal overlap analysis. Scale bars: whole cell 1 µm; close-ups 1 µm. (F) Similar to *Gl*CHC, anti-PI(3)P antibodies (anti-PI(3)P) detect foci of PI(3)P accumulation in close proximity to *Gl*PXD3 epitope-tagged reporters (anti-HA) in HA::*Gl*PXD3-expressing cells analysed with STED microscopy. Scatter plots are included for signal overlap analysis. Scale bars: whole cell 1 µm; close-ups 1 µm.

## Discussion

### PIPs and PIP binders in *G. lamblia*

PIPs are recognized spatiotemporal organizers and decorate the surface of the eukaryotic cell’s plasma and endo –membrane system [1–3]. *G. lamblia* is no exception; despite its significant reduction in endomembrane complexity, this species maintains a variety of PIP residues, mostly located at the cell periphery. We identified 13 novel proteins, in most cases of unknown function, that carry predicted PIP-binding modules and primarily localize in close proximity to PVs.

All hitherto identified PIP-binding proteins in *G. lamblia* can be loosely grouped in two categories; they are either relatively small proteins (up to 400 amino acid residues) consisting almost entirely of the PIP-binding module (e.g. *Gl*PXD6 and *Gl*NECAP1) or they are large proteins consisting of domains of unknown function associated to a single predicted domain for PIP-binding (e.g. *Gl*PXD2 and *Gl*FYVE). A full functional characterization of the latter is a challenge given the level of genomic sequence divergence in *G. lamblia*. This makes it currently difficult to determine whether sequences are lineage-specific or so diverged as to be unrecognizable orthologues of previously-characterized proteins. Hence, structural annotation of large *G. lamblia* proteins carrying PIP-binding modules such as *Gl*PXD2 or *Gl*FYVE is limited to the lipid binding domain.

Eight out of 14 identified PIP-binding modules are either directly or indirectly associated to clathrin assemblies. Their PIP binding preferences, as measured using *in vitro* lipid-binding assays, are clearly distinct despite showing a varying degree of promiscuity, consistent with previously published data [20]. In contrast to previous reports, we could not measure PIP residue binding activity for *Gl*FYVE using *in vitro* lipid-binding assays [22]. Furthermore, *Gl*NECAP1 showed a distinctive and highly-specific binding preference for cardiolipin. This is a surprising finding since cardiolipin is an abundant phospholipid of the inner mitochondrial membrane [56] whose presence in *Giardia* is controversial [57, 58]. Although *Gl*NECAP1 lacks canonical motifs for cardiolipin binding [59], previous reports on the identification of cardiolipin-binding PH domains [60, 61] lend support to the observation that the PH-like domain in *Gl*NECAP1 could bind cardiolipin, at least *in vitro*. The evolutionary implications for the presence of cardiolipin in an organism with “bare-bones” mitochondrial remnants i.e. mitosomes, with no maintenance of membrane potential nor ATP synthesis activity [62], provide for an exciting research direction worth pursuing.

### An interactome-based model for PIP-binding proteins and clathrin assemblies at PVs

Unpublished data derived from APEX-mediated tEM experiments on transgenic trophozoites expressing APEX-tagged clathrin assembly components (*Gl*CHC and *Gl*CLC; [19]) show how larger PVs are associated to more than one PM-derived clathrin-marked invagination (Fig. 10A). This is supported by data from IFA and STED microscopy analysis of trophozoites loaded with Dextran-OG and labelled with anti-*Gl*CHC antibodies (Fig. 10B). By combining APEX-derived tEM data with STED microscopy data for both Dextran-OG and *Gl*PXD3 labelling, a quantified suborganellar model for PV organization can be built which takes into account organelle size and relative distribution of clathrin assemblies (Fig. 10C). In this model, *Gl*PXD3 clearly emerges as a membrane coat that surrounds individual PV organelles (Fig 10C, upper panel) on the cytoplasmic side of clathrin assemblies at the PV-PM interface (Fig 10C, lower panel).

**Figure 10:**
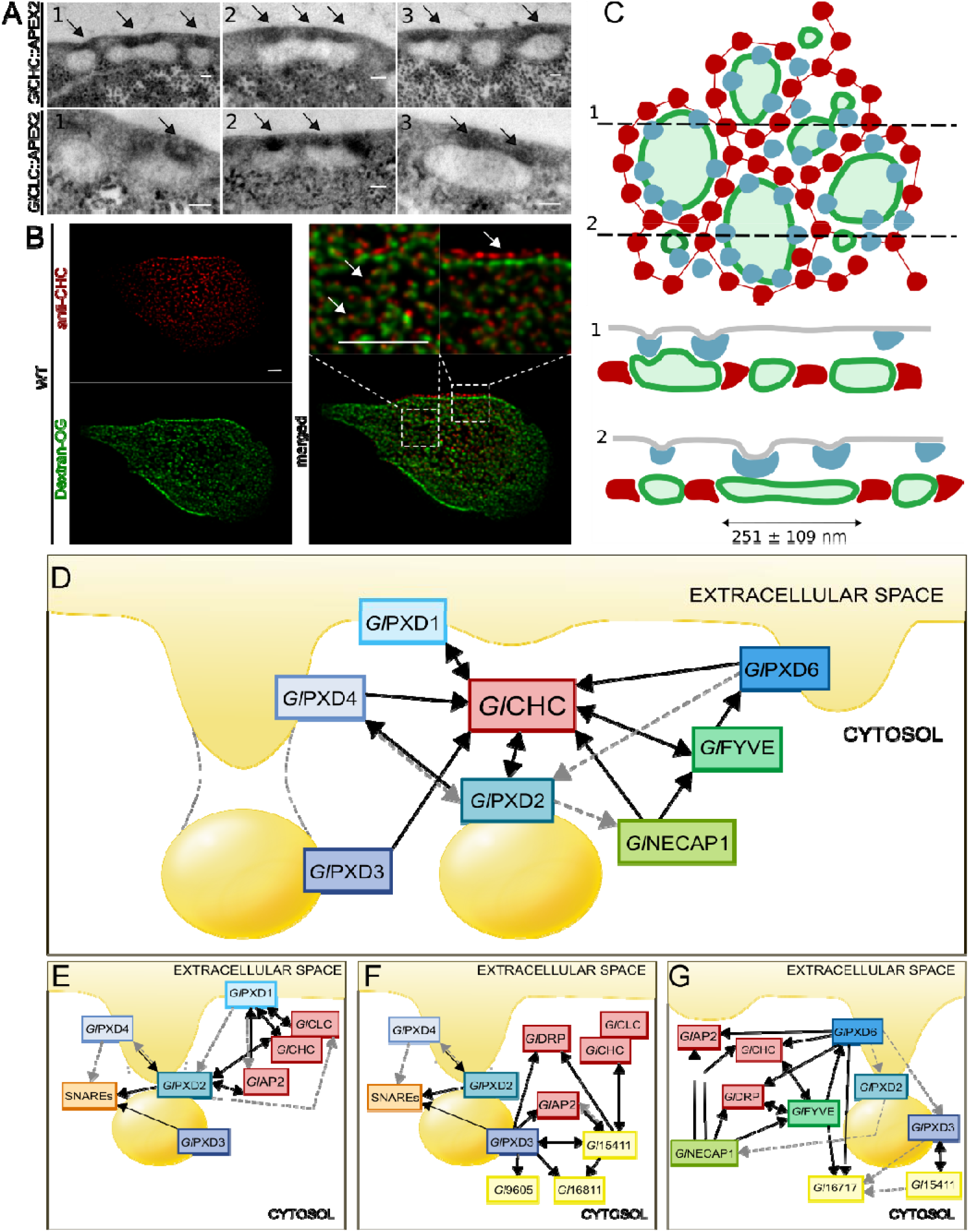
A working model for PV-associated nanoenvironments defined by clathrin assemblies and PIP-binding proteins. (A) Electron microscopy images of *G. lamblia* cells expressing an APEX2-tagged *Gl*CHC (upper panels) or *Gl*CLC (lower panels) reporter show darker APEX2-derived deposits at the PM-PV interface (arrows). Scale bar: 0.1 µm. (B) IFA analysis of a representative non-transgenic wild-type cell labelled with Dextran-OG and anti-*Gl*CHC antibodies to illuminate PV lumina and the PV-PM interface, respectively. Scale bar: 1 µm. (C) Schematic reconstruction of a surface view (left panel) of the PV system associated to clathrin assemblies (blue) and *Gl*PXD3 coats (red), based on data presented in this report and in [13]. PV membranes and lumina are represented in dark and light green, respectively. Cross-sections at (1) and (2) yield views in the right panel, highlighting foci of clathrin assemblies beneath the PM, above *Gl*PXD3’s coat-like deposition pattern surrounding PVs. (D) An overview of the *G. lamblia* PIP-binding interactome associated to PVs. All represented PIP-binding proteins were found to contact clathrin assemblies (*Gl*CHC) in either reciprocal (double-headed arrows) or one-way (single-headed arrows) modes of interaction, following filtering of co-IP data either at high (black solid lines) or low (grey dashed lines) stringency. (E-G) Nanoenvironments defined by specific sets of interaction partners including clathrin assemblies, PIP-binding proteins, SNARES and proteins of currently unknown function.

The PV-associated PIP-binding protein interactome appears as a tightly knit molecular network with *Gl*CHC at its center (Fig 10D and S5). Despite the high level of interconnectivity of distinct PIP-binder interactomes (Fig S5), specific molecular circuits such as the ones defined by the SNARE quartet (Fig 10E), pseudokinase *Gl*15411 and novel DLP *Gl*9605 (Fig 10F), as well as StAR-related lipid-transfer protein *Gl*16717 (Fig 10G), can be recognized. Notably, *Gl*PXD1 and 2 are the only PIP-binders whose extended interactomes include the *G. lamblia* clathrin light chain (Fig 10E and S5), arguably *Gl*CHC’s closest binding partner. The *Gl*PXD1 interactome further stands out for enrichment of proteasome-associated components (Table S1), invoking scenarios concerning clathrin assembly turnover in *G. lamblia*. Although previous data show that clathrin assemblies are long-lived stable complexes [19], they would still require remodeling, degradation, and substitution with new components. In the absence of classical components as well as C-terminal motifs on *Gl*CHC for ordered disassembly of clathrin coats, *Gl*PXD1’s proteasome-enriched interactome points to proteasome-mediated degradation of *Gl*CHC assemblies as an alternative process to achieve turnover albeit without recycling of coat components.

In the context of clathrin assembly dynamics, *Gl*NECAP1 once again comes to the forefront. NECAP1 is characterized as an AP2 interactor and an important component of CCVs in the assembly phase [53]. Given that CCVs have not been detected in *Giardia*, this begs the question of the functional role of a NECAP1 cardiolipin-binding orthologue in *G. lamblia* which was found to interact with *G. lamblia* AP2 subunits and *Gl*FYVE. Recent developments in gene knock-out [63] and CRISPR-Cas9-based knock-down [64, 65] methodologies tailored to *G. lamblia* will be instrumental towards a full functional characterization of *Gl*NECAP1’s function(s)

### Perturbation of PIP binding homeostasis affects fluid-phase uptake

We initially hypothesized that perturbation of either PIP saturation or PIP-binding activity would elicit fluid-phase uptake phenotypes by impacting PV functionality. The hypothesis tested positive for the saturation of PI3P, PI(3,4,5)P_3,_ and PI(4,5)P_2_. A significant effect on cell width was also detected when PI(3)P binding sites were saturated by overexpressing 2xFYVE::GFP (Figs 3B and F), linking PIP residues to both endocytic homeostasis and overall maintenance of cell size, possibly in connection to membrane turnover. Complementing these data, ectopic expression of both *Gl*FYVE and *Gl*NECAP1 significantly impacted fluid-phase uptake. Furthermore, ectopic expression of *Gl*NECAP1 induced an enlarged PV phenotype similar to that induced by expression of a predicted GTP-locked *Gl*DRP mutant [66]. Ectopic expression of a truncated *Gl*FYVE deprived of its ARM repeats induced the formation of membrane-bounded compartments of undefined origins. ARM folds are superhelical structures mostly involved in protein-protein interactions [67], suggesting that a loss of these domains may impact *Gl*FYVE function and protein complex formation. In line with this hypothesis, the NT-*Gl*FYVE epitope-tagged reporter loses association to PVs. In contrast to the *Gl*FYVE-induced uptake phenotype and despite a severe PV clustering phenotype, HA::*Gl*PXD2-expressing cells still appear to perform fluid-phase uptake comparably to wild-type cells. This suggests that PV morphology can be decoupled from effective PV-mediated uptake. Taken together, these data link PIPs to clathrin assemblies and fluid-phase PV-mediated uptake, providing new insights on clathrin’s hitherto unclear role in *Giardia* endocytosis.

### Beyond clathrin assemblies

Investigation of the molecular milieu within which clathrin-associated PIP-binding proteins operate in *G. lamblia* revealed two protein sets of special interest. Four predicted SNARE proteins were detected in both the *Gl*PXD2 and *Gl*PXD3 interactomes. Further investigations will be necessary to determine whether the function of this SNARE quartet is indeed fusing PM and PV membranes at contact sites, thereby allowing entry of fluid-phase material into PV organelles.

Another finding of special interest concerns *Gl*9605, a hitherto unrecognized DLP found in the interactome of *Gl*PXD3 with similarity to bacterial DLPs (BDLPs; Fig S6). Similar to their eukaryotic counterparts, BDLPs are capable of helical self-assembly and tubulation of lipid bilayers, and were shown to be most closely related to the mitofusins FZO and OPA (Fig S6) [24, 25], but only distantly related to classical dynamins [26]. BDLPs were also found in the Archaea class Methanomicrobia [68], making the family ubiquitously distributed across all kingdoms. These data show how the DLP/DRP family in *G. lamblia* has now expanded to include the previously unidentified endocytosis-associated *Gl*9605 BDLP homologue. *Gl*DRP plays a role in the regulation of PV and encystation-specific vesicle (ESV) size [66]. Although its role in fluid-phase uptake has not been determined, expression of a GTP-locked *Gl*DRP mutant inhibited endocytosis of biotinylated surface proteins [66]. On the other hand, a similar mutational analysis of *Gl*9605 shows that this DLP variant can elicit a dominant-negative fluid-phase uptake phenotype. Although we did not test *Gl*9605 involvement in surface protein uptake, the data so far suggest that two distinct DLPs play independent albeit complementary roles in the regulation of PV-mediated fluid-phase uptake and organelle homeostasis.

In this work, we report on the detailed functional characterization of PIP-binding proteins in *G. lamblia* that associate to clathrin assemblies. Our data reveals a previously unappreciated level of complex interplay between lipid residues and their protein binders in marking and shaping endocytic compartments in this parasite. However, several identified PIP-binding modules appear to associate to PVs *independently* of clathrin. Their extended interactomes and their involvement in fluid-phase uptake have yet to be investigated but current data point towards a complex network of PIP binders of varying binding preference and affinity, all working in the same subcellular environment, yet, in some cases (*Gl*FERM, *Gl*BAR1 and 2, *Gl*PROP1 and 2, Gl16801), not directly linked to clathrin assemblies. The only known exception is *Gl*epsin whose localization remains controversial due to conflicting reports [21, 69]. We systematically did not detect *Gl*epsin in any of the interactomes for clathrin-associated PIP binders, in line with its localization at the ventral disk [21]. Altogether, the variety of PIP residues and PIP-binding modules in the *G. lamblia* cortical area containing endocytic PVs underscores their necessity for correct functioning of membrane traffic even in a protist so clearly marked by reduction in endomembrane complexity.

## Materials and Methods

### Giardia lamblia cell culture, induction of encystation and transfection

*G. lamblia* WBC6 (ATCC catalog number 50803) trophozoites were cultured and harvested applying standardized protocols [52]. Encystation was induced by the two-step method as previously described [70, 71]. Transgenic cell lines were generated using established protocols by electroporation of linearized or circular pPacV-Integ-based plasmid vectors prepared from *E.coli* as described in [72]. Transgenic lines were then selected for puromycin resistance (final conc. 50 µg ml ^−1^). After selection, transgenic trophozoites carrying integrated or episomal reporter constructs were further cultured with or without puromycin, respectively.

### Construction of expression vectors

Oligonucleotide sequences used for cloning in this work are listed in Table S8. pPacV-Integ-based [34] expression of epitope tagged reporter constructs was driven using either putative endogenous (pE) or encystation-dependent (pCWP1) promoters. Constructs 2xFYVE::GFP and GFP::P4C [39] were kindly provided by Dr. H. Hilbi (University of Zurich).

### PV labelling using fluid-phase markers

Fluid-phase uptake assay in *G. lamblia* was performed as described previously [26] using dextran coupled to either Oregon Green 488 (Dextran-OG) (Cat. Nr. D-7171, Thermo Fisher Scientific) or Texas-Red (Dextran-R) (Cat. Nr. D-1863, Thermo Fisher Scientific) fluorophores, both at 1mg/ml final concentration.

### Co-immunoprecipitation with limited cross-linking

Co-immunoprecipitation *Gl*PXD1-6, *Gl*NECAP1, and *Gl*FYVE was done as previously reported [19, 42]. Protein input was standardized to 0.8mg/ml total protein.

### Protein analysis and sample preparation for mass spectrometry (MS)-based protein identification

Protein analysis was performed on 4%/10% polyacrylamide gels under reducing conditions (molecular weight marker Cat. Nr. 26616, Thermo Scientific, Lithuania). Immunoblotting was done as described in [73]. Gels for mass spectrometry (MS) analysis were stained using Instant Blue (Expedeon, Prod. # iSB1L) and destained with ultra-pure water.

### Mass Spectrometry, protein identification and data storage

MS-based protein identification was performed as described in [19]. Free access to raw MS data is provided through the ProteomeXchange Consortium on the PRIDE platform [74]. Accession numbers for datasets derived from bait-specific and corresponding control co-IP MS analyses are the following: PXD013890 for *Gl*PXD1, 3 and 6, PXD013897 for *Gl*FYVE, PXD013896 for *Gl*NECAP and PXD013899 for *Gl*PXD2 and 4.

### In silico co-immunoprecipitation dataset analysis

Analysis of primary structure and domain architecture of putative components of giardial PIP--binding proteins was performed using the following online tools and databases: SMART for prediction of patterns and functional domains (http://smart.embl-heidelberg.de/), pBLAST for protein homology detection (https://blast.ncbi.nlm.nih.gov/Blast.cgi?PAGE=Proteins), HHPRED for protein homology detection based on Hidden Markov Model (HMM-HMM) comparison (https://toolkit.tuebingen.mpg.de/#/tools/hhpred), PSORTII for sub-cellular localisation prediction (https://psort.hgc.jp/form2.html), TMHMM for transmembrane helix prediction (http://www.cbs.dtu.dk/services/TMHMM/), RCSB for 3D structure of homologues (https://www.rcsb.org/), and the *Giardia* Genome Database to extract organism-specific information such as protein expression levels, predicted molecular sizes and nucleotide/protein sequences (www.giardiaDB.org). The generated co-IP datasets were filtered using a dedicated control-co-IP dataset generated using non-transgenic wild-type parasites. Filtration of the bait-specific co-IP and control-co-IP datasets was done using Scaffold4 (http://www.proteomesoftware.com/products/) with high stringency parameters (95_2_95, FDR 0%) and low stringency parameters (95_2_50, FDR 0%). Furthermore, exclusive hits for bait-specific datasets were manually curated using the following criteria for inclusion into the interactome model: i) exclusive detection with > 3 spectral counts in bait-specific datasets or ii) an enrichment of peptide counts >3 with respect to the ctrl. co-IP dataset. Data presented in Tables S1-7 show exclusive and non-exclusive protein hits filtered using both stringency levels.

### Immunofluorescence analysis (IFA) and light-microscopy

Samples for immunofluorescence and analysis of subcellular distribution of reporter proteins by wide-field and laser scanning confocal microscopy (LSCM) were prepared as described previously [33, 35]. Nuclear DNA was labelled with 4’, 6-diamidino-2-phenylindole (DAPI). The HA epitope tag was detected with either the anti-HA antibody (1:50 or 1:100; Anti HA high affinity 3F10, Cat. Nr. 11867423001, Roche), anti-V5 (1:50 or 1:100; V5 Tag Monoclonal Antibody, Cat. Nr. R960-25, Thermo Fisher Scientific) or self-made antibodies raised against *Gl*CHC (dilution 1:1000) followed by an anti-rat antibody coupled to fluorochrome in case of wide-field or confocal microscopy (1:200; Goat anti-Rat IgG (H+L) Cross-Adsorbed Secondary Antibody, Alexa Fluor 488, Cat. Nr. #A11006, Invitrogen) and for STED microscopy (Goat anti-Rat IgG (H+L) Cross-Adsorbed Secondary Antibody, Alexa Fluor 594, Cat. Nr. A11007, Invitrogen). Specific PIP residues were detected using anti-PI(3)P (1:100; Purified anti-PI(3)P IgG, Z-P003 Echelon Biosciencies), antiPI(4,5)P_2_ (1:100; Purified anti-PI(4,5)P_2_ IgM, Z-P003 Echelon Biosciencies) and anti-PI(3,4,5)P_3_ (1:100; Purified anti-PI(3,4,5)P_3_ IgM, Z-P045 Echelon Biosciencies) followed by an anti-mouse antibody coupled to fluorochrome in all three cases (Goat anti-Mouse IgG (H+L) Cross-Adsorbed Secondary Antibody, Alexa flour 594, Cat. Nr. A-11005, Thermo Fischer Scientific or Goat anti-Mouse IgG (H+L) Cross-Adsorbed Secondary Antibody, Alexa flour 488, Cat. Nr. A-11017, Thermo Fischer Scientific). Cells were generally imaged at maximum width, with nuclei and the bare-zone at maximum diameter. Deconvolution was performed with Huygens Professional (Scientific Volume Imaging). Three-dimensional reconstructions and signal overlap quantification (Mander’s coefficient) in volume images of reconstructed stacks were performed using IMARIS x64 version 7.7.2 software suite (Bitplane AG) or FIJI [75], respectively.

### Super resolution (gSTED) microscopy

Sample preparation was done as described for wide field microscopy and LSCM. For imaging, samples were mounted in ProLong Gold antifade reagent (Cat. Nr. P36934, Thermo Fisher Scientific). Super resolution microscopy was performed on a LSCM SP8 gSTED 3x Leica (Leica Microsystems) at the Center for Microscopy and Image Analysis, University of Zurich, Switzerland. Nuclear labelling was omitted due to possible interference with the STED laser. Further data processing and three dimensional reconstructions of image stacks were done as described for LSCM.

### Sample preparation for transmission electron microscopy

Transgenic trophozoites expressing *Gl*PXD2 (GL50803_16595) and non-transgenic line were harvested and analysed by transmission electron microscopy (tEM) as described previously [66].

### DAB staining in APEX2 expressing cells

Transgenic trophozoites expressing *Gl*NECAP1::APEX2-2HA, *Gl*CHC::APEX2-2HAand *Gl*CLC::APEX2-2HA were harvested and washed with PBS followed by fixation in 2.5% EM grade glutaraldehyde in cacodylate buffer (100 mM cacodylate (Cat. Nr. 20838), 2mM CaCl_2_ (Cat. Nr. 21097, Fluka) in PBS) for 1h at RT. Samples were washed twice before and after quenching for 5 min in 20mM glycine/cacodylate buffer. For staining, cells were resuspended in 500ul substrate solution containing 1.4mM DAB tetrahydrodhloride (Cat. Nr. D5637, Sigma) with 0.3mM H_2_O_2_ (Cat. Nr. H1009, Sigma) in cacodylate buffer and incubated for 15 min. The reaction was terminated by washing thrice in cacodylate buffer and prepared as described for tEM.

### Chemical fixation of DAB-stained cells

DAB stained cell suspicions were post-fixed with 1% aqueous OsO4 for 1 hour on ice, subsequently rinsed three rimes with pure water and dehydrated in a sequence of ethanol solutions (70% up to 100%), followed by incubation in 100% propylene oxide and embedding in Epon/Araldite (Sigma-Aldrich, Buchs, Switzerland). Samples were polymerised at 60°C for 24h. Thin sections were imaged pre- and post-staining with aqueous uranyl acetate (2%) and Reynolds lead citrate.

### Expression and Purification of Bacterial Fusion Proteins

For each candidate PIP-binding protein, corresponding nucleotide stretches coding for selected amino acid residues (Table S9) were modified by including an HA-coding sequence either at the 5’ end or the 3’ end and then subcloned into the pMal-2Cx *E. coli* expression vector (New England Biolabs). The resulting recombinant variants were expressed as maltose-binding protein (MBP) fusions in *E.coli* (strain Bl21) and grown in LB medium either at 37°C (MBP-*Gl*PXD1, MBP-*Gl*PXD2, MBP-*Gl*PXD3, MBP-*Gl*PXD6, MBP-*Gl*NECAP1 and MBP-*Gl*FYVE) or 30°C (MBP-*Gl*PXD4 and MBP-*Gl*PXD5) to an OD_600_=0.4. Induction of expression was performed by adding0.2 mM IPTG (Isopropyl β-D-1-thiogalactopyranoside, Cat. Nr. 15529019, Thermo Fischer Scientific) to the cultures and incubating for a further 4 hours. Cells were harvested at 4°C (4,000 x g) and bacterial pellets were resuspended in 5 ml of cold column buffer with 1x PIC (Protease inhibitor cocktail set I; Cat. Nr. 539131-10VL, Merck) and 200 mM PMSF (Cat. Nr. 329-98-6, Sigma Aldrich). Cells were lysed by sonication and centrifuged (20 min, 9,000 × g, 4°C). Cleared supernatant was incubated with amylose resin slurry (Amylose Resin High Flow, Cat. Nr. E8022L, BioLabs) for 4 hours at 4°C on a turning wheel, washed with column buffer and then transferred to an empty column (BioRad 15 ml). Unbound protein was washed using until background OD_280_ reached ~0.06. Protein fractions were eluted using 10mM maltose solution and pooled for overnight dialysis in a dialysis cassette (Slide-a-Lyzer, Cat. Nr. 66380, Thermo Fischer Scientific) against 25mM NH_4_Ac at 4°C and later lyophilized. Protein fractions were stored at −80°C.

### Protein lipid overlay (PLO) assay

E. coli-derived lyophilized proteins were reconstituted in 1x PBS and protein concentration was measured using the Bradford assay. PIP strips (PIP strips, Cat. Nr. P-6001 and P-6002, Echelon) or PIP arrays (PIP strips, Cat. Nr. P-6100, Echelon) were first floated on ultrapure water for 5 min before incubation in blocking buffer (1xPBS containing 0.1%v/v Tween-20 and 3% fatty-acid free BSA (Sigma A7030)) at RT for 1h. Thereafter 0.5 μg/ml of protein in PBS containing 3% fatty acid free BSA were incubated for 1h at RT with gentle agitation. After washing with 1xPBS containing 0.1% v/v Tween-20, PIP–strips were incubated (1h, RT, agitated) with a monoclonal anti-HA antibody (clone 3F10, monoclonal antibody from Roche) at a dilution of 1:500 in blocking buffer. Subsequently strips were washed and incubated (1h, RT, agitated) with a goat-derived polyclonal anti-rat antibody conjugated to HRP at a dilution of 1:2000 in blocking buffer (Cat. Nr. 3050-05, Southern Biotech). After further washing, strips were developed using a chemiluminescent substrate (WesternBright ECL HRP Substrate, Cat. Nr. K-12045-D50).

### Densitometric analysis of lipid strips and arrays

Relative quantification of immunoblotting signal intensity on PIP strips and arrays overlaid with PIP-binding proteins was performed using FIJI [75]. For each strip or array, the spot with the highest pixel number was set as a reference for 100% binding; signals coming from all other spots were normalized against it. The data were visualized as bar charts of relative signal intensity as a measure of lipid-binding preference for each PIP-module.

### Identification of giardial orthologues of known PIP-binding domains

PIP-binding domain representatives were used as bait for *in silico* searches within the *Giardia* genome database (GDB) (http://giardiadb.org/) using the online tool HHpred (https://toolkit.tuebingen.mpg.de/) to detect remote giardial homologues using hidden Markov models (HMMs; Table 1) [25]. Outputs were firstly evaluated based on the calculated probability and the corresponding E-value for the prediction, with cut-offs for probability and e-value set to 90 and 1e-10, respectively. Sequence identity and similarity were also considered. To validate the prediction, candidate giardial PIP-binding proteins were then utilized as baits to search PDB databases using HHpred to retrieve orthologous PIP-binding proteins/modules. For additional validation, I-TASSER [29–31]was also used to predict hypothetical structures of putative giardial PIP-binding domains next step validation.

### Multiple sequence alignment analysis

Multiple sequence alignment using two or more sequences was performed with the Clustal Omega sequence alignment algorithm [76, 77]. The sequences used to compile the alignments shown in supplementary figure 1 were chosen based on representative members for each PIP-binding domain type [1, 10, 78]. Alignments for figures 6 and 8 were based on previously characterized G1-G4 GTP binding motifs [50] and NECAP1 proteins [53], respectively.

### De novo structural modeling and analysis

Ab-initio prediction of hypothetical 3D models presented in supplementary figure 1 was done using I-TASSER [29–31]. The best model was chosen based on the C-score predicted by the algorithm. A C-score is a measure of confidence for a model based on the significance of threading template alignments and the convergence parameters of the structure assembly simulations. It ranges from −5-to 2, with higher C-scores indicating higher confidence. The final 3D structures were displayed using PyMOL (The PyMOL Molecular Graphics System, Version 2.0 Schrödinger, LLC.). The superimposition of *Giardia* PIP-binding proteins with their closest structural orthologue are based on I-TASSER predictions, with structural similarities expressed by TM-score and RMSD^a^ values. The TM-score is computed based on the C-score and protein length. It ranges from 0 to 1, where 1 indicates a perfect match between two structures. RMSD^a^ is the root mean square deviation between residues that are structurally aligned by TM-align [79]. Specifically for *Gl*BAR1 and 2, the structural overlap analysis was performed by selecting positively-charged residues from previously characterized BAR domains shown to play a role in lipid binding [80]. These were manually superimposed on corresponding residues in the predicted *Gl*BAR1 and 2 structures.

### Phylogenetic analysis

Subjected sequences of GTPase domains were aligned using Clustal Omega tool. The tree construction was submitted to a PHYLogeny Inference Package (PHYLIP) program [81, 82] using random number generator seed set to 111 and number of bootstrap trials set to 10000. The tree was visualised using the on-line tool iTOL and includes branch lengths as a measure of evolutionary distance [83].

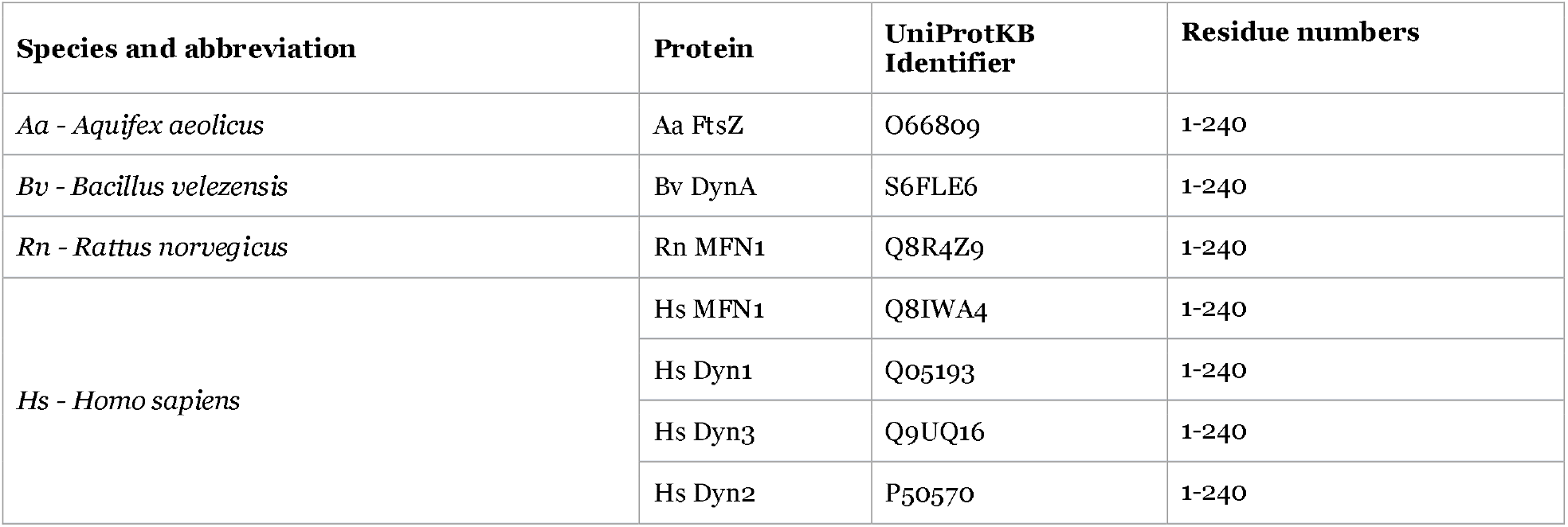
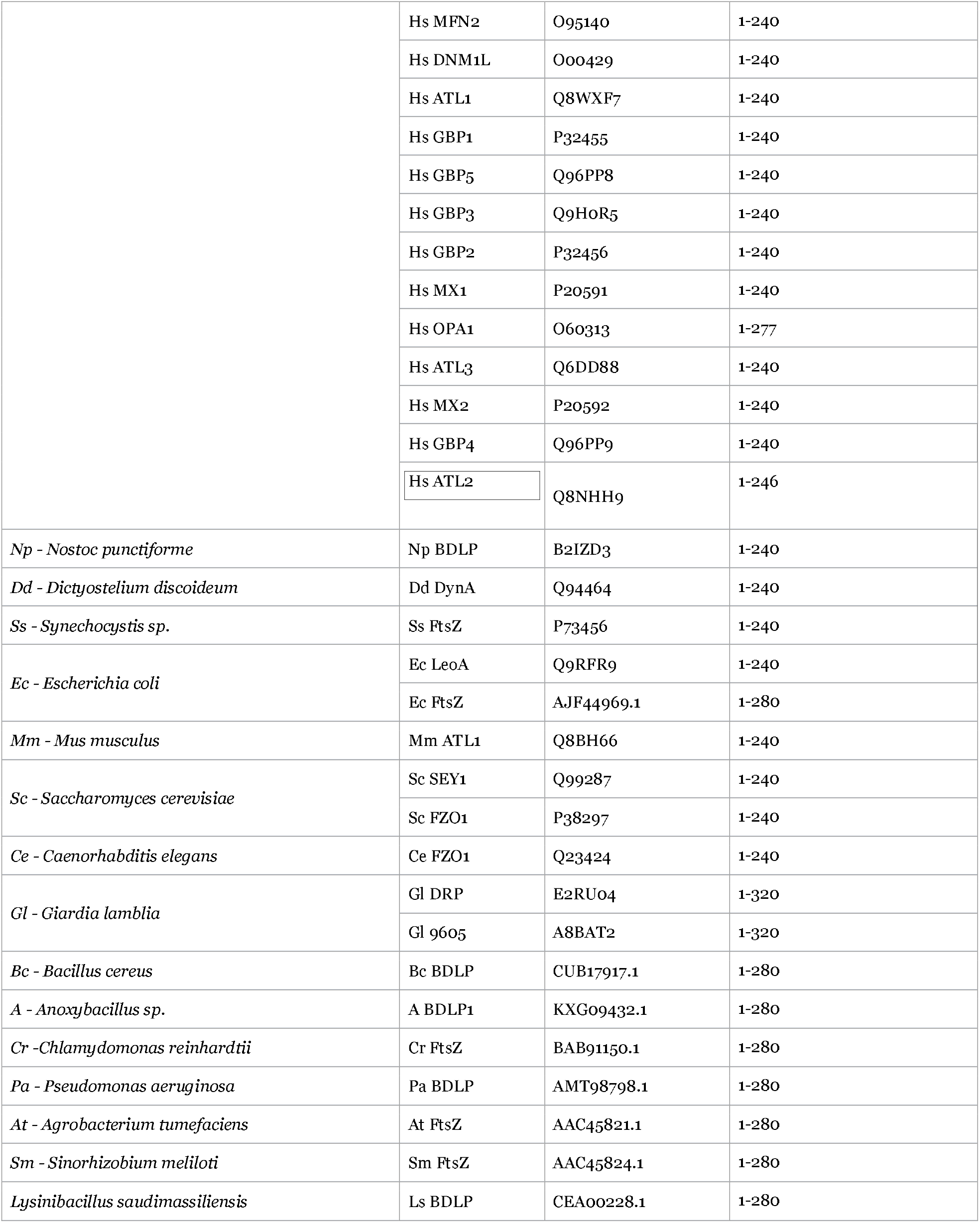

Supplemental Tables 1-7: Proteins identified in the interactomes of *Gl*PXD1-4 and 6, *Gl*FYVE and *Gl*NECAP1

Supplemental Table 8: List of oligonucleotide names and sequences for construct synthesis

Supplemental Table 9: Amino acid sequences of lipid-binding modules used in vitro for protein lipid-overlay assay

Supplemental Figure 1: Multiple sequence alignment and structural prediction analysis of *G. lamblia* PIP-binding domains.

Supplemental Figure 2: Lipid-binding properties of *Giardia*-lipid binding domains.

Supplemental Figure 3: Subcellular distribution of PI(3)P, PI(4,5)P2 and PI(3,4,5)P3 in *G. lamblia* trophozoites

Supplemental Figure 4: APEX-mediated electron microscopy analysis of *Gl*NECAP1 subcellular deposition.

Supplemental Figure 5: Overview of core protein interactomes determined from co-IP analyses

Supplemental Figure 6: Phylogenetic analysis and tree reconstruction for the predicted GTPase domain of the novel dynamin-like protein *Gl*9605.

## Acknowledgements

ABH was supported by Swiss National Science Foundation grants 140803 and 125389. Florian Schmidt and Ritta Rabbat are acknowledged for their technical assistance. Dr. Hubert Hilbi (University of Zurich) is gratefully acknowledged for sharing *Legionella*-derived constructs 2xFYVE::GFP and GFP::P4C.

