## supplemental figure 1 for "Phosphoinositide-binding proteins mark, shape and functionally modulate highly-diverged endocytic compartments in the parasitic protist *Giardia lamblia*"

**Supplemental Figure 1: Multiple sequence alignment and structural prediction analysis of *G. lamblia* PIP-binding domains.**

For all PIP-binding modules except *Gl*NECAP1, the sequence of the lipid-binding domain was aligned to its respective homologous domains. Each domain was structurally modelled using I-TASSER (blue) and superimposed on its closest structural homolog (green). For each structural overlay a TM-score and RMSD value are reported, followed by a blow-up of the calculated location of known and predicted PIP-binding motifs. (**A-B**) *Gl*epsin, (**C-E**) *Gl*FYVE and *Gl*16801, (**F-L**) *Gl*PXD1-6, (**M-O**) *Gl*BAR1 and *Gl*BAR2, (**P-Q**) *Gl*FERM and (**R-T**) *Gl*PROP1 and *Gl*PROP2. (**U**) Legend to color code for conserved/similar residues. (**V**) Closest structural homologues, including their origin and identifiers, for structural overlay analysis of PIP-binding modules in *G. lamblia*. (**W**) Selected orthologues, including their origin and identifiers, for each *G. lamblia* PIP-binding module used in the MSA analysis to highlight conserved/similar residues for lipid-binding.

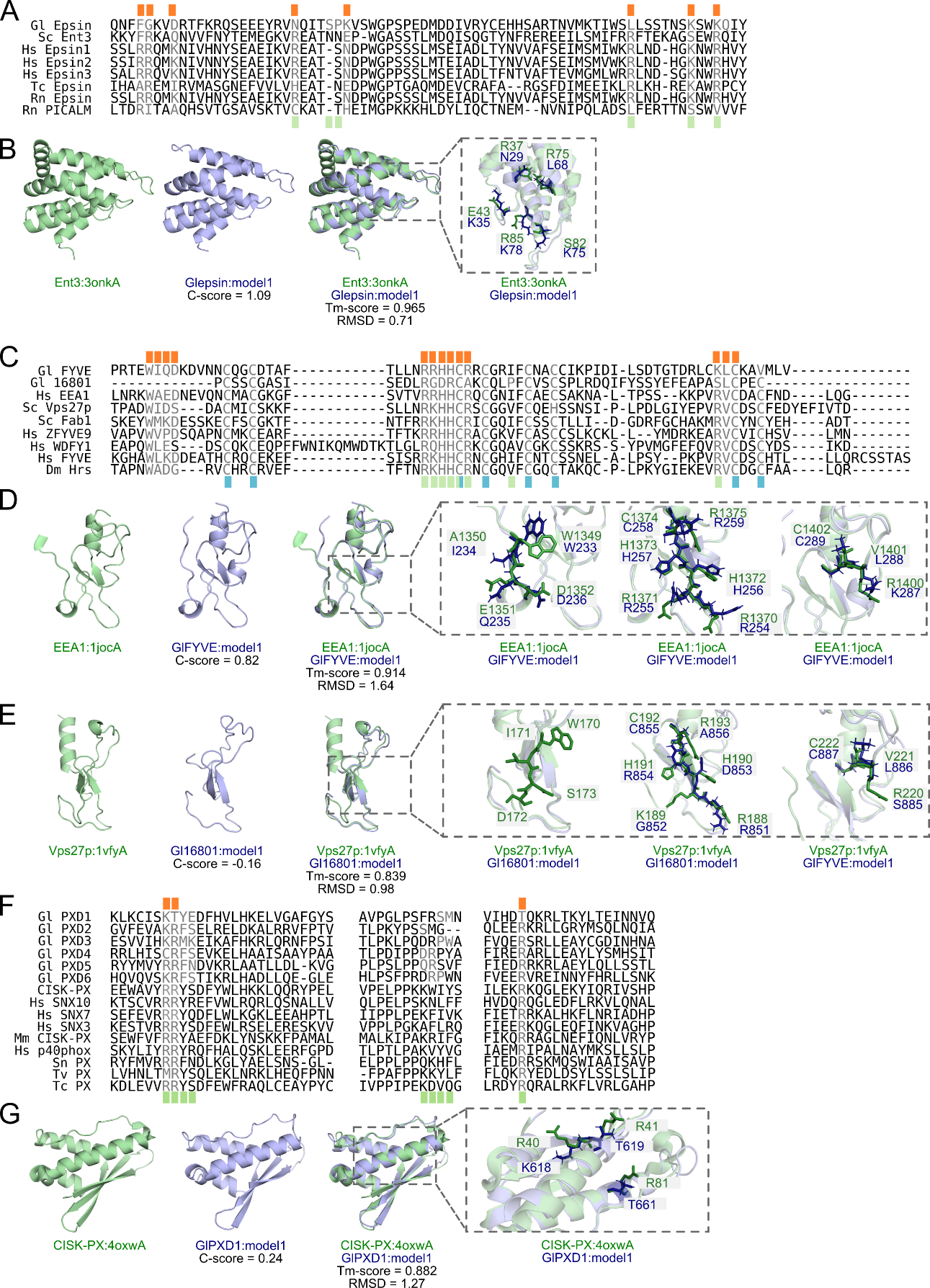

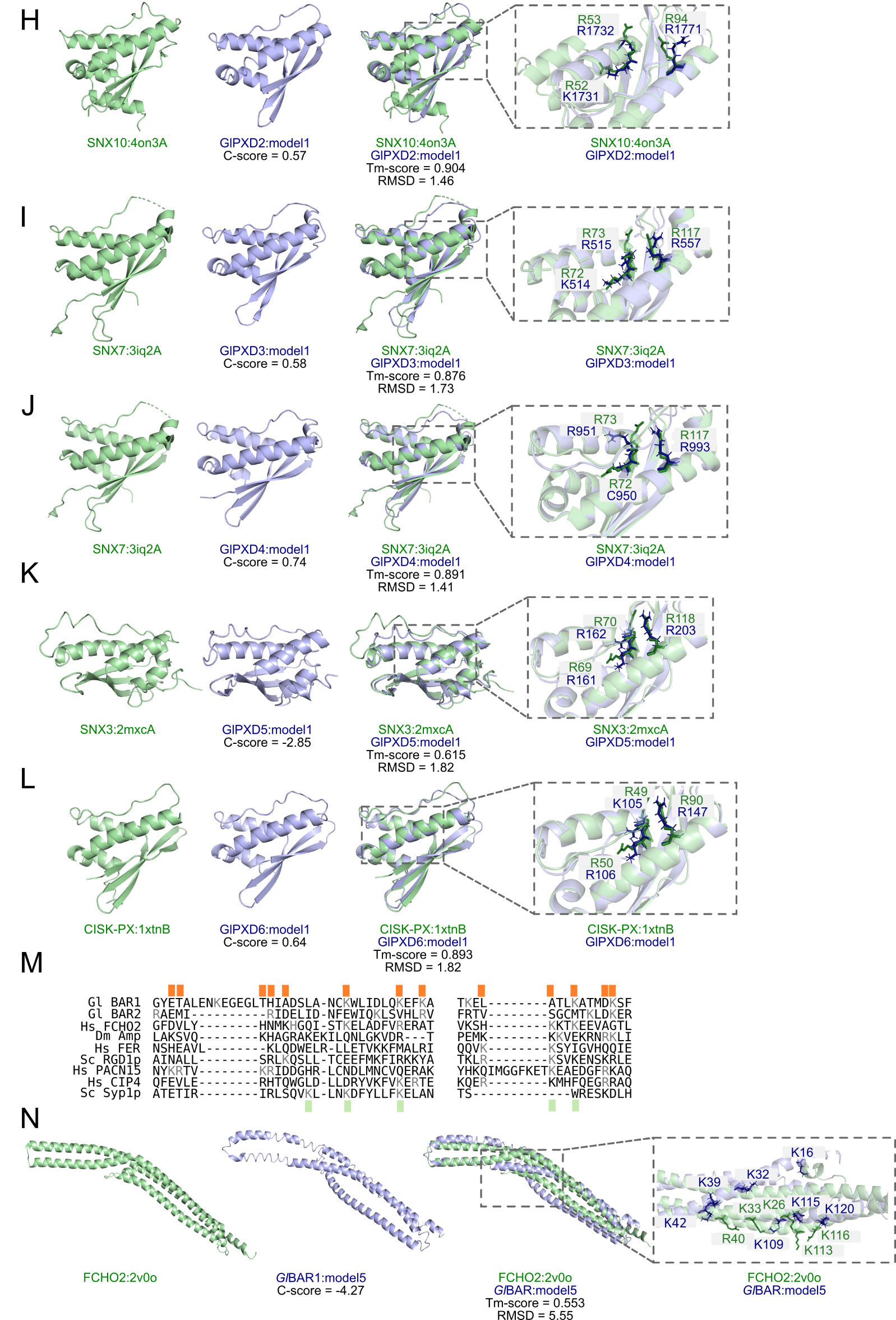

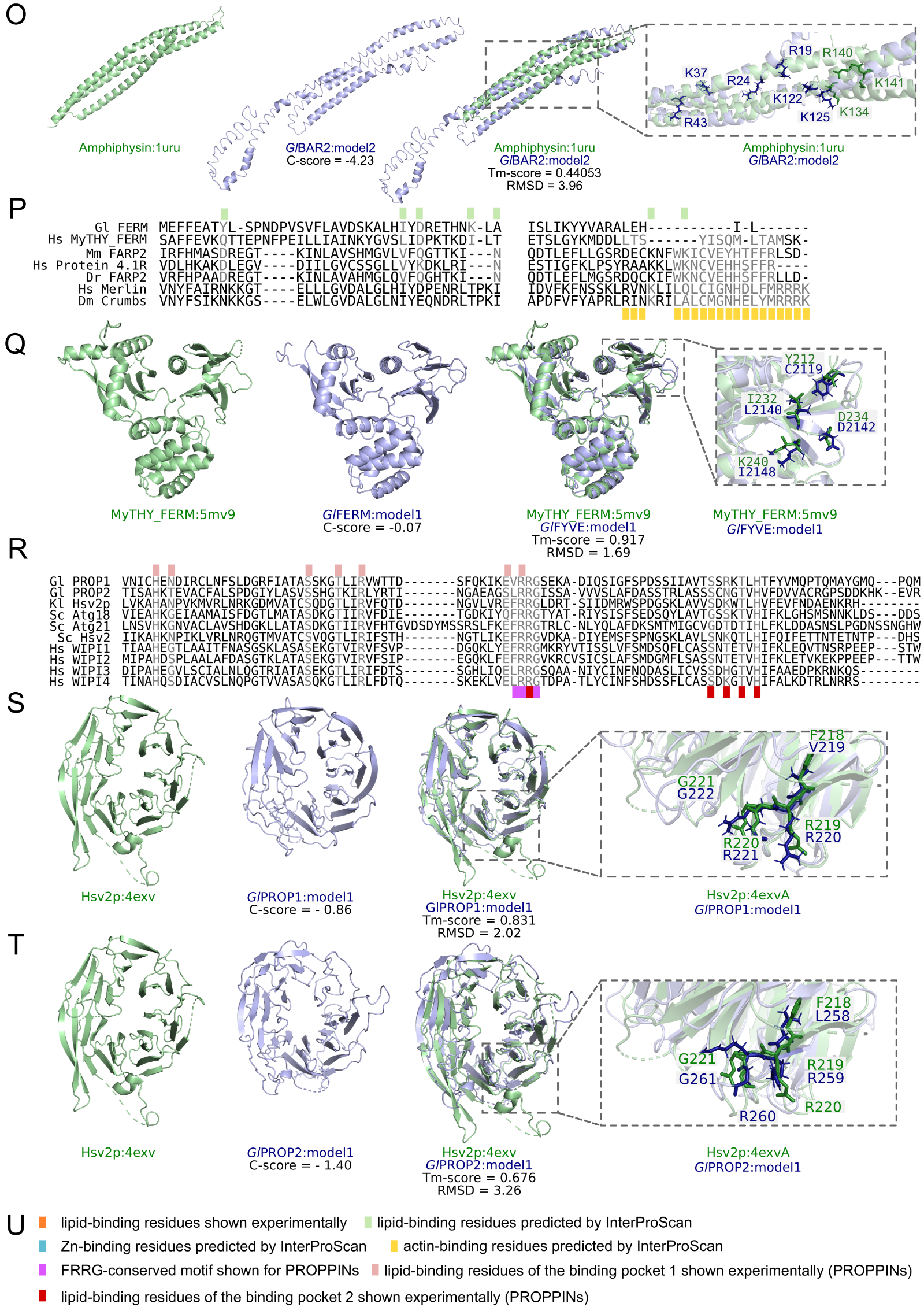

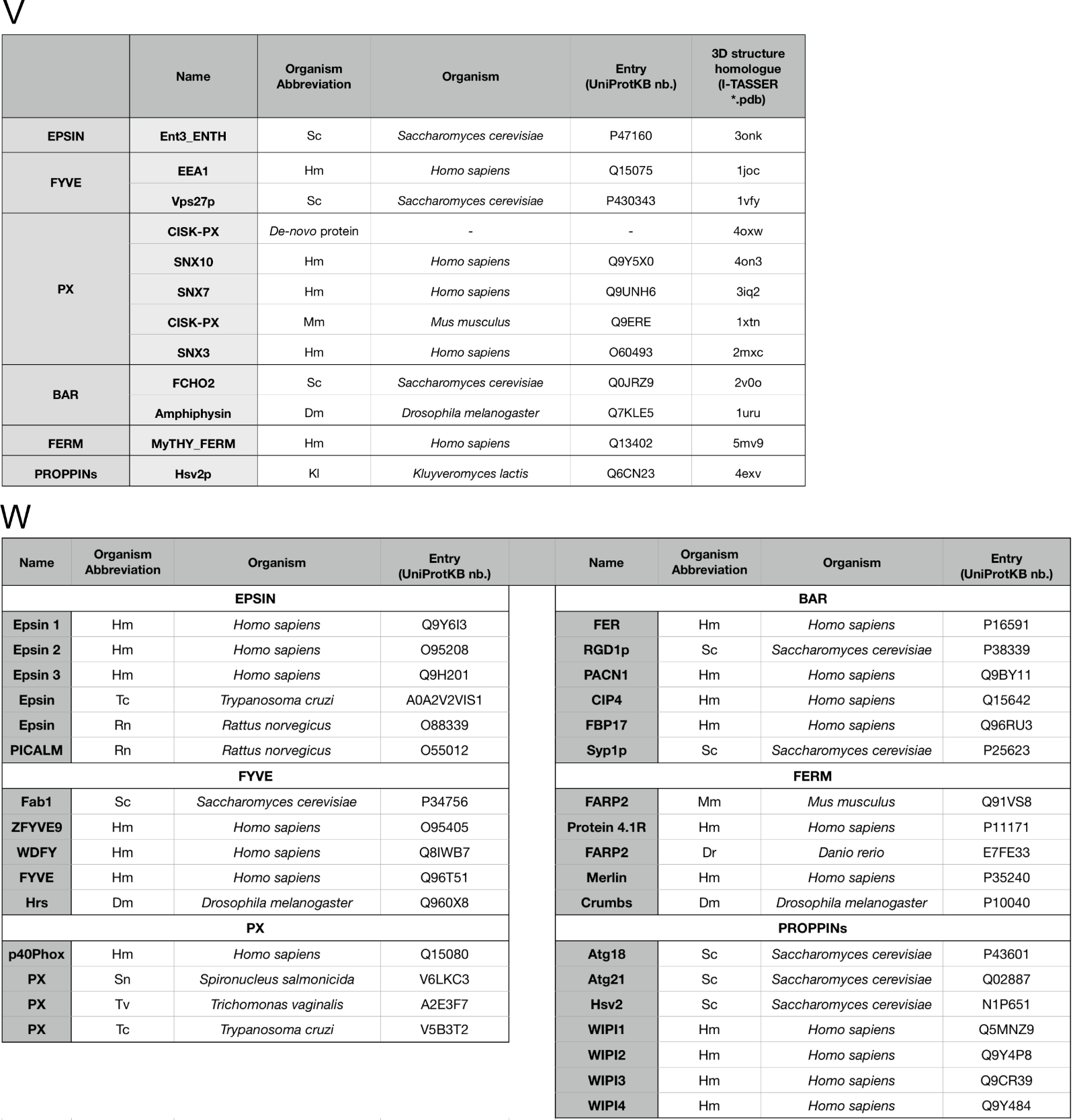
