## supplemental figure 2 for "Phosphoinositide-binding proteins mark, shape and functionally modulate highly-diverged endocytic compartments in the parasitic protist *Giardia lamblia*"

**Supplemental Figure 2: Lipid-binding properties of *Giardia*-lipid binding domains.**

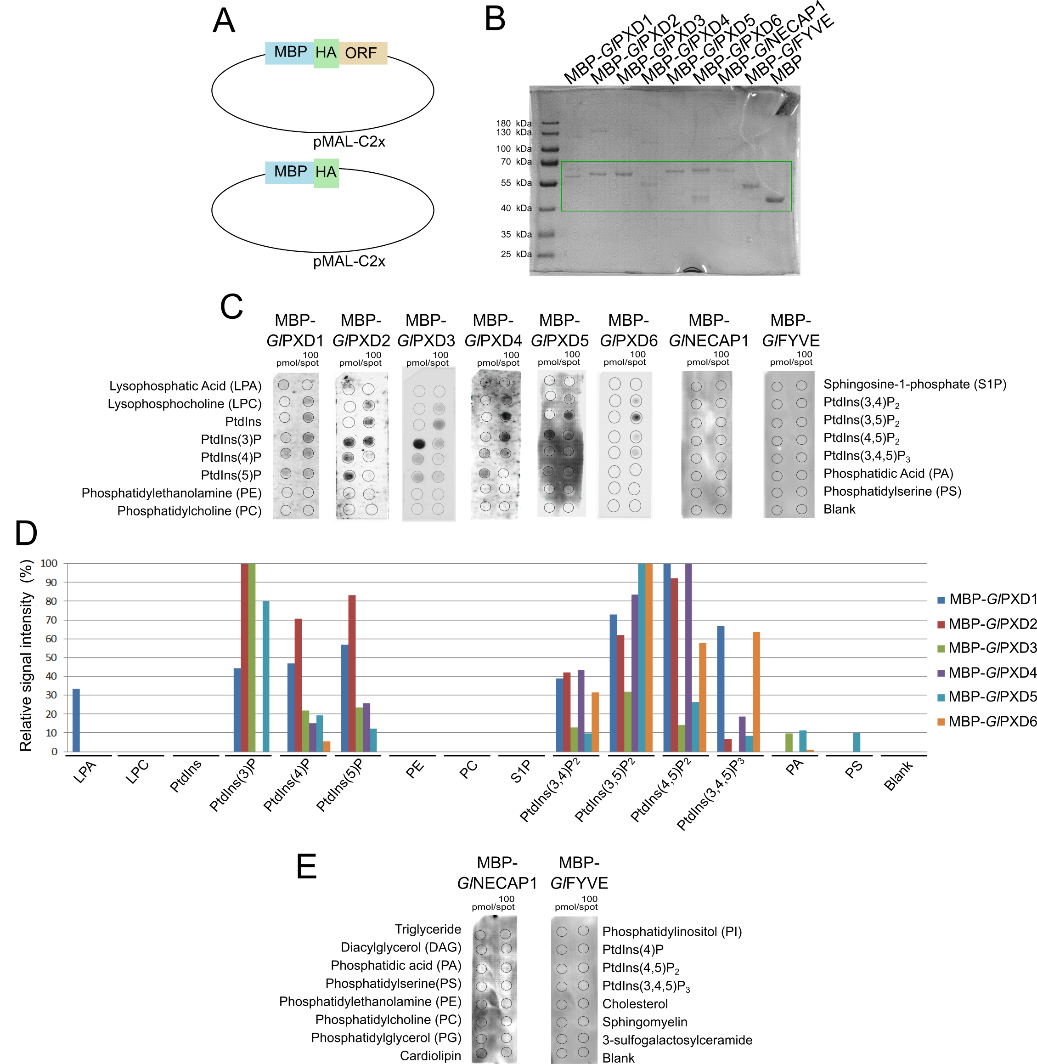

Lipid-binding and immuno-detection analysis of *G. lamblia* PIP-binding domains from proteins *Gl*PXD1-6, *Gl*FYVE and *Gl*NECAP1 using lipid strips. (**A**) Schematic diagram of the pMAL-p2Cx vector used for heterologous expression of individual PIP-binding domains in *E.coli*. (**B**) SDS-PAGE analysis of recombinant epitope-tagged MBP-PIP binding domain fusions normalised to 1µg total protein. Protein ladder sizes are included in the first lane. (**C**) Immuno-detection of epitope-tagged MBP-fusions for each PIP-binding domain overlayed on lipid strips carrying spotted lipid residues at 100pmol/spot and visualized by chemiluminescence. (**D**) Lipid binding preferences for all tested MBP-domain fusions, measured using FIJI and visualized as plots of relative signal intensity for each probed lipid residue. Values were normalized to those of lipid residues presenting strongest signal intensity. (**E**) Lipid binding preferences for *Gl*NECAP1 and *Gl*FYVE investigated using a different set of spotted lipid residues revealed *Gl*NECAP1’s exclusive affinity for cardiolipin.
