## supplemental figure 3 for "Phosphoinositide-binding proteins mark, shape and functionally modulate highly-diverged endocytic compartments in the parasitic protist *Giardia lamblia*"

**Supplemental Figure 3: Subcellular distribution of PI(3)P, PI(4,5)P2 and PI(3,4,5)P3 in *G. lamblia* trophozoites**

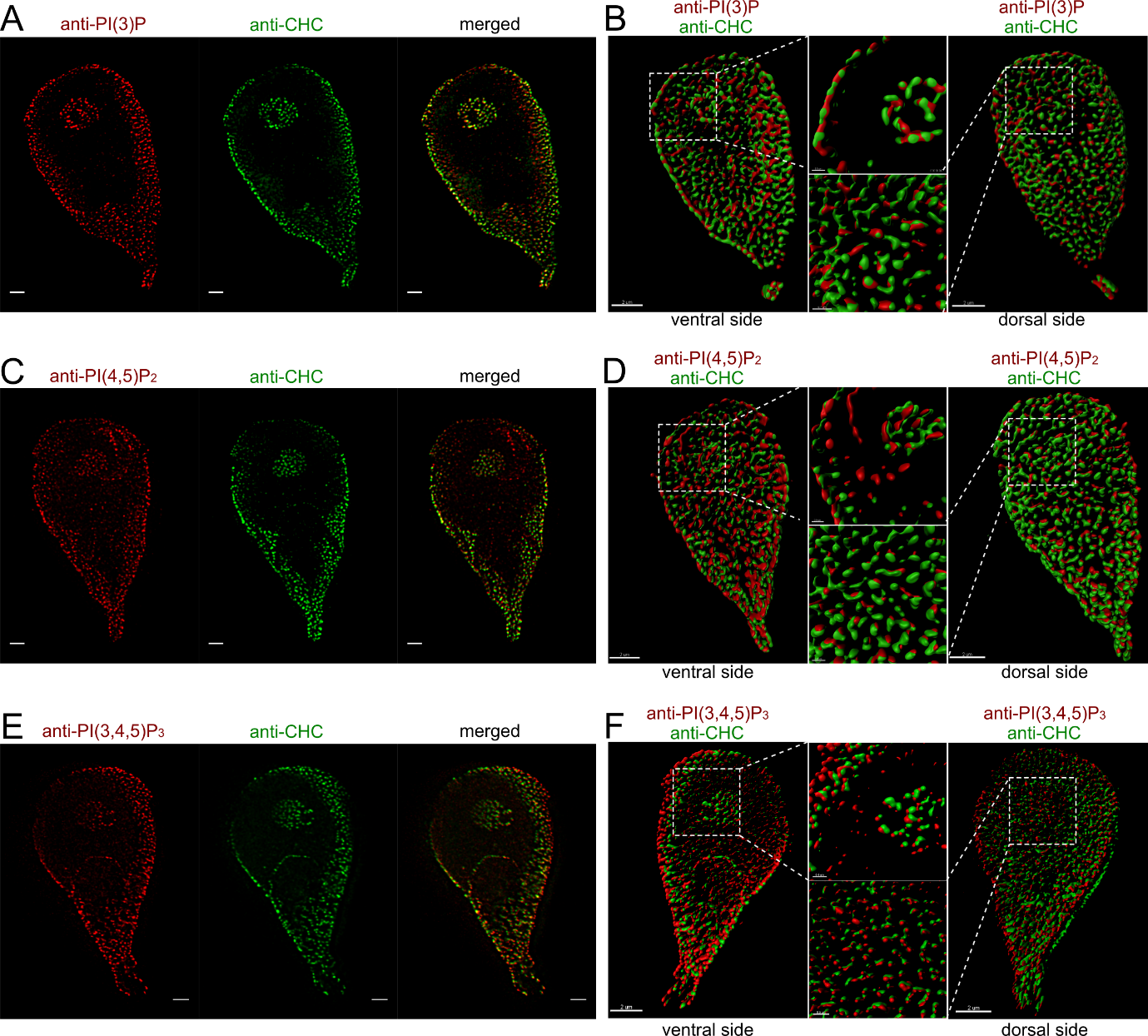

3D STED microscopy analysis followed by signal overlap and deconvolution of representative non-transgenic wild-type G. lamblia trophozoites co-labeled with anti-GlCHC (in green) antibody and either (A-B) anti-PI(3)P, (C-D) anti-PI(4,5)P2, or (E-F) anti-PI(3,4,5)P3 antibodies (in red). Dorsal and ventral sides are defined with respect to the ventral disk. Scale bar for (A, C, E): 1 µm. Scale bar for (B, D, F): 2 µm. Scale bar for insets in (B, D, F): 0.5 µm.
