## supplemental figure 4 for "Phosphoinositide-binding proteins mark, shape and functionally modulate highly-diverged endocytic compartments in the parasitic protist *Giardia lamblia*"

**Supplemental Figure 4: APEX-mediated electron microscopy analysis of *Gl*NECAP1 subcellular deposition.**

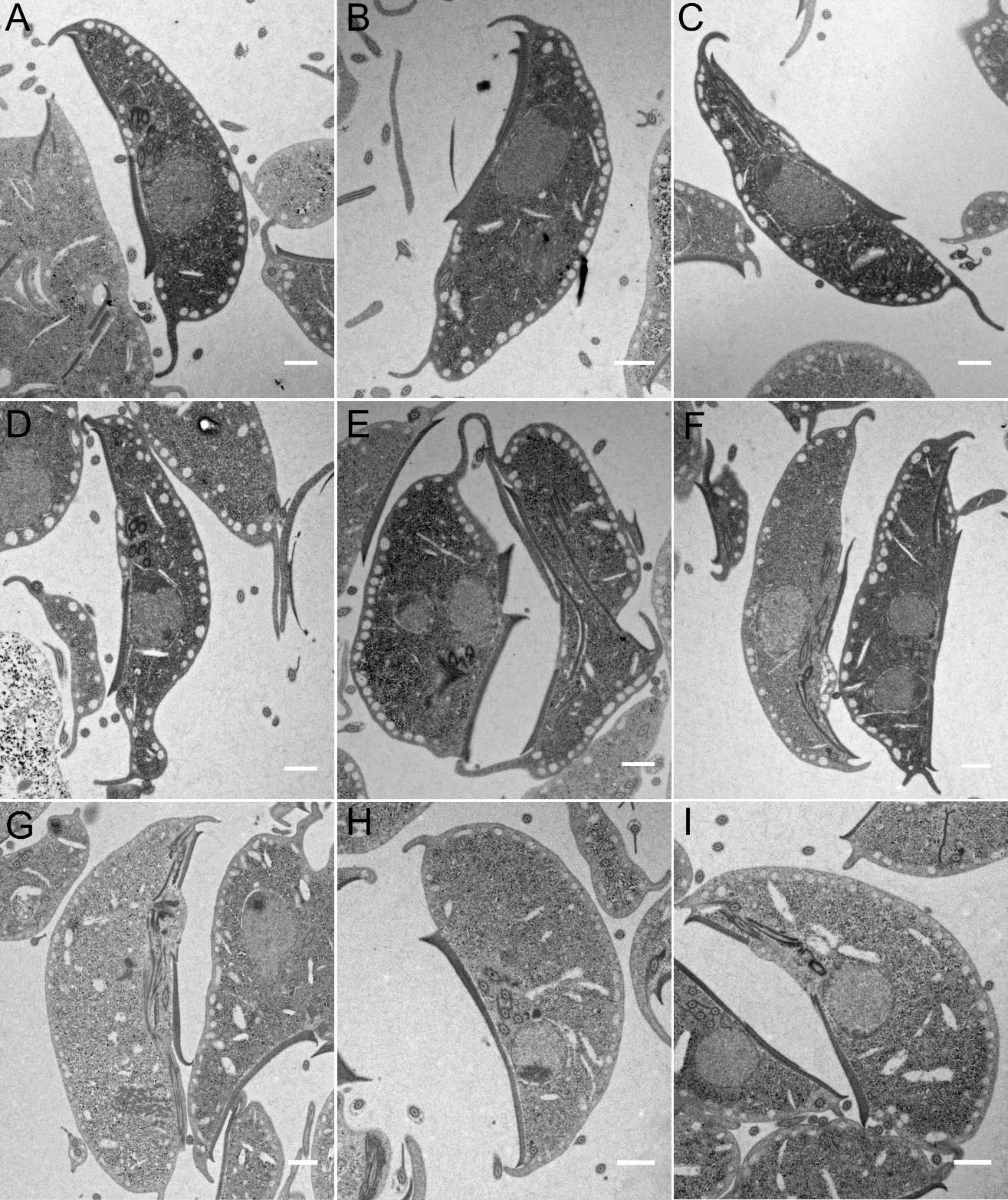

(**A-F**) Representative images of transgenic *G. lamblia* cells expressing construct pE*Gl*NECAP1::APEX2-2HA showing enlarged PVs and a diffused APEX-dependent cell staining signal. (**G-I)** Non-transgenic control cells. Scale bar: 1µm.
