## supplemental figure 5 for "Phosphoinositide-binding proteins mark, shape and functionally modulate highly-diverged endocytic compartments in the parasitic protist *Giardia lamblia*"

**Supplemental Figure 5: Overview of core protein interactomes determined from co-IP analyses**

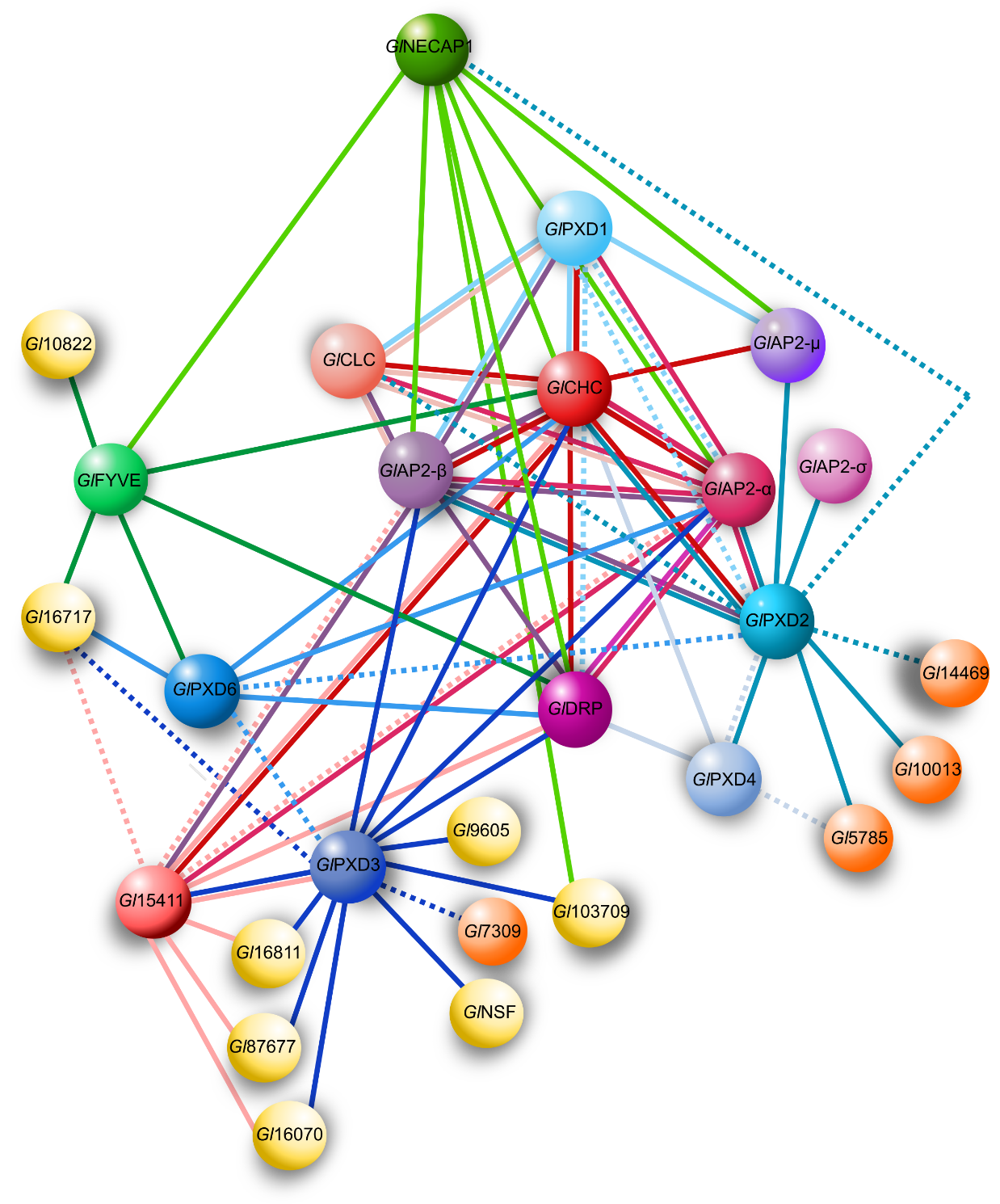

Interactomes for *Gl*FYVE, *Gl*NECAP1 and *Gl*PXD3, 4 and 6 defined by co-IP analysis were integrated with previously published data [19] for core clathrin assembly, *Gl*PXD1, *Gl*PXD2 and *Gl*FYVE interactomes. Solid lines: interactions detected at high stringency. Dashed lines: interactions detected at low stringency. Yellow partners are currently annotated on GDB as “hypothetical protein” *i.e.* proteins of unknown function.
