## supplemental figure 6 for "Phosphoinositide-binding proteins mark, shape and functionally modulate highly-diverged endocytic compartments in the parasitic protist *Giardia lamblia*"

**Supplemental Figure 6: Phylogenetic analysis and tree reconstruction for the predicted GTPase domain of the novel dynamin-like protein *Gl*9605.**

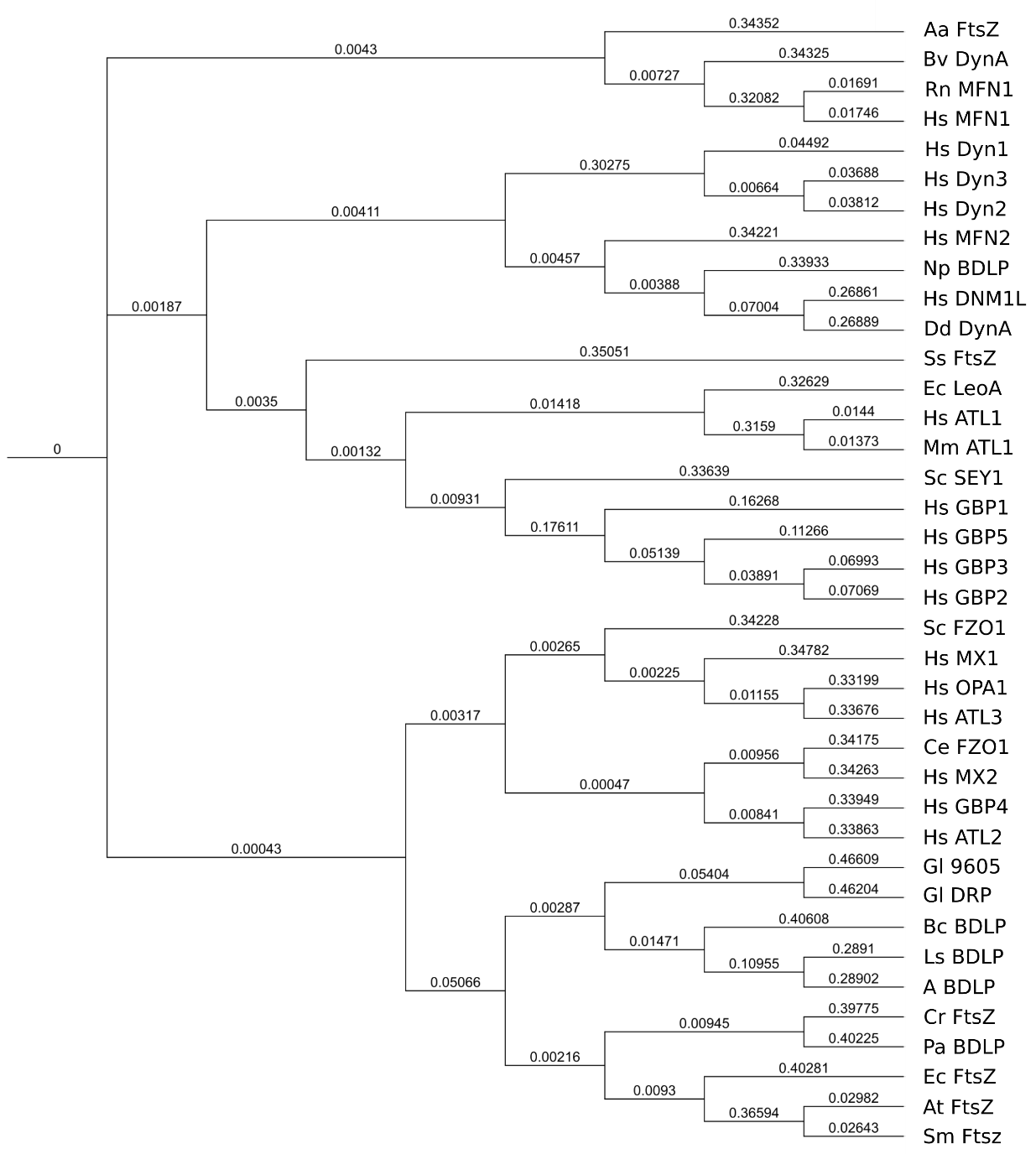

Phylogenetic analysis of predicted GTPase domains from the following prokaryotic and eukaryotic species used to compute the tree shown in the figure, including branch lengths as a measure of evolutionary distance: Aa - *Aquifex aeolicus*, Bv - *Bacillus velezensis*, Rn - *Rattus norvegicus*, Hm - *Homo sapiens*, Np - *Nostoc punctiforme,* Dd - *Dictyostelium discoideum*, Ss - *Synechocystis sp.*, Ec - *Escherichia coli*, Mm - *Mus musculus*, Sc - *Saccharomyces cerevisiae*, Ce - *Caenorhabditis elegans*, Gl - *Giardia lamblia*, Bc - *Bacillus cereus*, Ls - *Lysinibacillus saudimassiliensis*, A - *Anoxybacillus sp*., Cr - *Chlamydomonas reinhardtii*, Pa - *Pseudomonas aeruginosa*, At - *Agrobacterium tumefacies*, Sm - *Sinorhizobium meliloti*.
