## supplemental tables 1-7 for "Phosphoinositide-binding proteins mark, shape and functionally modulate highly-diverged endocytic compartments in the parasitic protist *Giardia lamblia*"

Supplemental Tables 1-7: Proteins identified in the interactomes of GlPXD1-4 and 6, GlFYVE and GlNECAP1

| Numbers Sheet Name | Numbers Table Name | Excel Worksheet Name |
|---|---|---|
| Table S1: GlPXD1 | | |
| | Table S1. Proteins identified in the interactome of GlPXD1 | Table S1_ GlPXD1 - Table S1. Pr |
| Table S2: GlPXD2 | | |
| | Table S2. Proteins identified in the interactome of GlPXD2 | Table S2_ GlPXD2 - Table S2. Pr |
| Table S3: GlPXD3 | | |
| | Table S3. Proteins identified in the interactome of GlPXD3 | Table S3_ GlPXD3 - Table S3. Pr |
| Table S4: GlPXD4 | | |
| | Table S4. Proteins identified in the interactome of GlPXD4 | Table S4_ GlPXD4 - Table S4. Pr |
| Table S5: GlPXD6 | | |
| | Table S5. Proteins identified in the interactome of GlPXD6 | Table S5_ GlPXD6 - Table S5. Pr |
| Table S6: GlFYVE | | |
| | Table S6. Proteins identified in the interactome of GlFYVE | Table S6_ GlFYVE - Table S6. Pr |
| Table S7: GlNECAP1 | | |
| | Table S7. Proteins identified in the interactome of GlNECAP1 | Table S7_ GlNECAP1 - Table S7. |

| Color Code |
|---|
| Bait |
| Clathrin assemblies |
| hypothetical protein |
| PIP-binding proteins |
| SNARE-related protein |
| Rab |
| Giardin |
| Tubulin |
| Proteasome-related protein |
| Enzyme |
| Kinase |
| Ribosomal protein |
| Protein 21.1 |
| Uncategorised protein |

### Supplementary Table 1 GL50803_7723 *Gl* PXD1

| | | bait specific spectral counts at given stringency parameters | | bait specific exclusive spectral counts at given stringency parameters | | InterProScan prediction | GO | HHPRED best hit (Probability E-value) |
|---|---|---|---|---|---|---|---|---|
| | | 95_2_95 | 95_2_50 | 95_2_95 | 95_2_50 | | | |
| ORF number | Annotation | Gl50803_7723 | Gl50803_7723 | Gl50803_7723 | Gl50803_7723 | | | |
| GL50803_7723 | Hypothetical protein | 38 | 86 | 38 | 86 | | | |
| GL50803_102108 | Clathrin heavy chain | 69 | 111 | 69 | | | | |
| GL50803_4259 | hypothetical protein | 7 | 13 | 7 | 13 | | | |
| GL50803_21423 | Beta adaptin | 21 | 38 | 21 | 38 | | | |
| GL50803_17304 | Alpha Adaptin | | 7 | | 7 | | | |
| GL50803_8917 | Mu adaptin | 4 | 7 | 4 | 7 | | | |
| GL50803_14373 | Dynamin (dynamin) | | 3 | | 3 | | | |
| GL50803_16595 | Liver stage antigen-like protein | | 3 | | 3 | | | |
| GL50803_21628 | hypothetical protein | | 11 | | 11 | - | - | Antitoxin RelB3 3BPQ_A (80.35/4.7) |
| GL50803_115337 | hypothetical protein | 9 | 10 | 9 | 10 | Ubiquitin-like domain family | GO:0005515 protein binding | UV excision repair protein RAD23 1OQY_A (98.6/1.2e-9) |
| GL50803_9861 | hypothetical protein | | 8 | | | - | - | Antitoxin RelB3 3BPQ_A (82.91/3.2) |
| GL50803_8627 | hypothetical protein | | 4 | | 4 | - | - | RNA splicing 5XJC_V (94.57/0.016) |
| GL50803_15251 | hypothetical protein | | 4 | | 4 | - | - | Nuclear pore complex protein 5IJO_Y (96.1/0.07) |
| GL50803_16659 | hypothetical protein | 3 | 3 | 3 | 3 | Proteasome component domain (PCI) | - | 26 proteasome subunit 3JCK_B (100/7.8e-32) |
| GL50803_15454 | hypothetical protein | | 3 | | 3 | Non-cytoplasmic domain | - | 26 proteasome subunit 3JCK_A (96.97/0.0016) |
| GL50803_11434 | 20S proteasome alpha subunit 2 | | 3 | | 3 | | | |
| GL50803_4365 | 26S protease regulatory subunit 6A | 3 | 4 | 3 | 4 | | | |
| GL50803_86683 | 26S protease regulatory subunit 7 | | 4 | | 4 | | | |
| GL50803_21331 | 26S protease regulatory subunit 7 | | 3 | | 3 | | | |
| GL50803_17106 | 26S protease regulatory subunit 8 | 4 | 7 | 4 | 7 | | | |
| GL50803_91643 | 26S proteasome regulatory subunit, putative | | 3 | | 3 | | | |
| GL50803_7110 | Ubiquitin | | 9 | | 9 | | | |
| GL50803_9779 | UPL-1 | | 3 | | 3 | | | |
| GL50803_17230 | Gamma giardin | | 7 | | 7 | | | |
| GL50803_114787 | Alpha-7.3 giardin | | 5 | | 5 | | | |
| GL50803_4812 | Beta-giardin | | 4 | | | | | |
| GL50803_103676 | Alpha-tubulin | | 3 | | 3 | | | |
| GL50803_101291 | Beta tubulin | 3 | 5 | 3 | 5 | | | |
| GL50803_16867 | AAA family ATPase | 6 | 8 | 6 | 8 | | | |
| GL50803_93358 | Alcohol dehydrogenase | 20 | 12 | | | | | |
| GL50803_11043 | Fructose-bisphosphate aldolase | 4 | 3 | | | | | |
| GL50803_9115 | Glucose-6-phosphate isomerase | | 3 | | 3 | | | |
| GL50803_21942 | NADP-specific glutamate dehydrogenase | | 6 | | 6 | | | |
| GL50803_93548 | Phospholipase B | | 5 | | 5 | | | |
| GL50803_15983 | Prolyl-tRNA synthetase | 3 | 6 | 3 | 6 | | | |
| GL50803_9413 | Protein disulfide isomerase PDI2 | 3 | 4 | 3 | 4 | | | |
| GL50803_32312 | Protein phosphatase 2C | 3 | 3 | 3 | 3 | | | |
| GL50803_17063 | Pyruvate-flavodoxin oxidoreductase | | 3 | | 3 | | | |
| GL50803_7439 | Ser/Thr phosphatase 2A, 65kDa reg sub A | | 6 | | 6 | | | |
| GL50803_113553 | Kinase, NEK | | 3 | | 3 | | | |
| GL50803_17054 | Acidic ribosomal protein P0 | 5 | 7 | | | | | |
| GL50803_16086 | Ribosomal protein L2 | | 6 | | 6 | | | |
| GL50803_16525 | Ribosomal protein L3 | 7 | 11 | 7 | 11 | | | |
| GL50803_17547 | Ribosomal protein L4 | 22 | 21 | | | | | |
| GL50803_17395 | Ribosomal protein L5 | | 3 | | 3 | | | |
| GL50803_17121 | Bip (BiP) | 43 | 20 | | | | | |
| GL50803_13747 | C4 group specific protein | | 5 | | 5 | | | |
| GL50803_15148 | Chaperone protein dnaJ | 3 | 7 | 3 | 7 | | | |
| GL50803_9808 | Chaperone protein DnaJ subfamily A | | 4 | | 4 | | | |
| GL50803_17483 | Chaperone protein DnaJ subfamily B | | 3 | | 3 | | | |
| GL50803_15591 | Coiled-coil protein | | 3 | | 3 | | | |
| GL50803_88765 | Cytosolic HSP70 | 14 | 24 | 14 | 24 | | | |
| GL50803_112304 | Elongation factor 1-alpha | 45 | 22 | | | | | |
| GL50803_17570 | Elongation factor 2 | | 4 | | 4 | | | |
| GL50803_98054 | Heat shock protein HSP 90-alpha | 4 | 7 | 4 | 7 | | | |
| GL50803_21321 | High cysteine membrane protein Group 5 | 3 | 3 | 3 | 3 | | | |
| GL50803_13864 | HSP 90-alpha | 3 | 4 | 3 | 4 | | | |
| GL50803_15106 | Importin beta-3-subunit | 6 | 9 | 6 | 9 | | | |
| Gl50803_5795 | Leucine-rich repeat protein | 3 | | 3 | | | | |
| GL50803_10577 | Nucleolar protein NOP5 | | 4 | | 4 | | | |
| GL50803_17060 | Protein 21.1 | | 3 | | 3 | | | |
| GL50803_27925 | Protein 21.1 | | 4 | | 4 | | | |
| GL50803_32778 | Protein 21.1 | | 3 | | 3 | | | |
| GL50803_9030 | Protein 21.1 | | 8 | | | | | |
| GL50803_10255 | Translation initiation factor eIF-4A, putative | | 3 | | 3 | | | |

Supplementary Table 2 GL50803_16595 *Gl* PXD2

| | | bait specific spectral counts at given stringency parameters | | bait specific exclusive spectral counts at given stringency parameters | | InterProScan prediction | GO | HHPRED best hit (Probability E-value) |
|---|---|---|---|---|---|---|---|---|
| ORF number | Annotation | 95_2_95<br>Gl50803_16595 | 95_2_50<br>Gl50803_16595 | 95_2_95<br>Gl50803_16595 | 95_2_50<br>Gl50803_16595 | | | |
| GL50803_16595 | Liver stage antigen-like protein | 365 | 501 | 365 | 501 | | | |
| GL50803_102108 | Clathrin heavy chain | 65 | 92 | 65 | | | | |
| GL50803_4259 | hypothetical protein | | 4 | | 4 | | | |
| GL50803_21423 | Beta adaptin | 18 | 34 | 18 | 34 | | | |
| GL50803_17304 | Alpha Adaptin | 6 | 13 | 6 | 13 | | | |
| GL50803_8917 | Mu adaptin | 7 | 12 | 7 | 12 | | | |
| GL50803_5328 | Sigma adaptin | 3 | 9 | 3 | 9 | | | |
| GL50803_14373 | Dynamin (dynamin) | | 3 | | 3 | | | |
| GL50803_42357 | hypothetical protein | 8 | 8 | 8 | 8 | | | |
| GL50803_5785 | hypothetical protein | 5 | 5 | 5 | 5 | | | |
| GL50803_10013 | hypothetical protein | 4 | 7 | 4 | 7 | | | |
| GL50803_14469 | Synaptobrevin-like protein | 2 | 9 | 2 | 9 | | | |
| GL50803_17332 | hypothetical protein | 37 | 76 | | | WD40-repeat-containing domain superfamily | GO:0005515 protein binding | serine/threonine-protein kinase<br>5YZV_A (95.38/0.55) |
| GL50803_9861 | hypothetical protein | 11 | 11 | | | - | - | Antitoxin RelB3<br>3BPQ_A (82.91/3.2) |
| GL50803_16404 | hypothetical protein | | 4 | | 4 | Thioredoxin-like superfamily | - | Protein disulfide-isomerase A5<br>4I6X_A (99.64/1.4e-17) |
| GL50803_16507 | hypothetical protein | | 4 | | 4 | ClpP/crotonase-like domain superfamily | - | Protein CT_858<br>3DOR_A (99.94/3.1e-28) |
| GL50803_16588 | hypothetical protein | | 3 | | 3 | Ribosomal protein P1/P2, N-terminal domain | - | 60S Acidic Ribosomal Protein P1<br>4BEH_B (99.92/2.6e-28) |
| GL50803_21628 | hypothetical protein | | 3 | | 3 | - | - | Antitoxin RelB3<br>3BPQ_A (80.35/4.7) |
| GL50803_17195 | hypothetical protein | | 2 | | 2 | PH-like domain superfamily | GO:0006897 endocytosis<br>GO:0016020 membrane | NECAP1<br>1TQZ_A (100/4.3e-52) |
| GL50803_17244 | Ribosomal protein L7a | 19 | 23 | | | | | |
| GL50803_1345 | Ribosomal protein L10a | 15 | 20 | 15 | 20 | | | |
| GL50803_11247 | Ribosomal protein L13a | 10 | 18 | | | | | |
| GL50803_7999 | Ribosomal protein S3 | 8 | 14 | 8 | 14 | | | |
| GL50803_14827 | Ribosomal protein S11 | 8 | 13 | 8 | 13 | | | |
| GL50803_16387 | Ribosomal protein L18a | 6 | 16 | 6 | 16 | | | |
| GL50803_10428 | Ribosomal protein L10 | 6 | 13 | 6 | 13 | | | |
| GL50803_14938 | Ribosomal protein L12 | 5 | 12 | 5 | 12 | | | |
| GL50803_16431 | Ribosomal protein L19 | 5 | 8 | 5 | 8 | | | |
| GL50803_4547 | Ribosomal protein S9 | 4 | 5 | 4 | 5 | | | |
| GL50803_4652 | Ribosomal protein S16 | 4 | 10 | | | | | |
| GL50803_10091 | Ribosomal protein L13 | 4 | 6 | 4 | 6 | | | |
| GL50803_14622 | Ribosomal protein L23 | 4 | 6 | 4 | 6 | | | |
| GL50803_6133 | Ribosomal protein L35a | 3 | 5 | 3 | 5 | | | |
| GL50803_7878 | Ribosomal protein S14 | 3 | | | | | | |
| GL50803_12981 | Ribosomal protein S5 | 3 | 6 | 3 | 6 | | | |
| GL50803_14329 | Ribosomal protein S7 | 3 | 4 | 3 | 4 | | | |
| GL50803_15228 | Ribosomal protein S15A | 3 | 3 | 3 | 3 | | | |
| GL50803_16310 | Ribosomal protein L27a | 3 | 5 | 3 | 5 | | | |
| GL50803_16525 | Ribosomal protein L3 | 3 | 8 | 3 | 8 | | | |
| GL50803_15520 | Ribosomal protein L21 | | 4 | | 4 | | | |
| GL50803_15046 | Ribosomal protein L26 | | 3 | | 3 | | | |
| GL50803_5947 | Ribosomal protein L35a | | 5 | | 5 | | | |
| GL50803_17056 | Ribosomal protein L9 | | 6 | | 6 | | | |
| GL50803_16652 | Ribosomal protein S13 | | 5 | | 5 | | | |
| GL50803_15260 | Ribosomal protein S15 | | 3 | | 3 | | | |
| GL50803_6135 | Ribosomal protein S17 | | 4 | | 4 | | | |
| GL50803_15551 | Ribosomal protein S18 | | 4 | | 4 | | | |
| GL50803_14699 | Ribosomal protein S23 | | 4 | | 4 | | | |
| GL50803_10367 | Ribosomal protein S24 | | 3 | | 3 | | | |
| GL50803_17054 | Acidic ribosomal protein P0 | 3 | 7 | | | | | |
| GL50803_11950 | Ribosomal protein L18 | | 3 | | 3 | | | |
| GL50803_8741 | Dipeptidyl-peptidase I precursor | | 3 | | 3 | | | |
| GL50803_17163 | Peptidyl-prolyl cis-trans isomerase B precursor | 3 | 4 | 3 | 4 | | | |
| GL50803_93938 | Triosephosphate isomerase | 5 | 6 | 5 | 6 | | | |
| GL50803_17121 | Bip (BiP) | | 16 | | | | | |
| GL50803_15591 | Coiled-protein | 5 | 6 | 5 | 6 | | | |
| GL50803_88765 | Cytosolic HSP70 | 10 | 22 | 10 | 22 | | | |
| GL50803_111950 | Dynein heavy chain | | 4 | | 4 | | | |
| GL50803_121045 | Histone H2B | 4 | | | | | | |
| GL50803_14212 | Histone H3 | | 6 | | 6 | | | |
| GL50803_135001 | Histone H4 | 3 | 4 | 3 | 4 | | | |
| GL50803_5795 | Leucine-rich repeat protein | 7 | | | | | | |
| GL50803_14521 | Peroxiredoxin 1 | 20 | 25 | | | | | |
| GL50803_15383 | Peroxiredoxin 1 | | 3 | | 3 | | | |

### Supplementary Table 3 GL50803_16596 GlPXD3

| ORF number | Annotation | bait specific spectral counts at given stringency parameters<br>95_2_95<br>Gl50803_16596 | 95_2_50<br>Gl50803_16596 | bait specific exclusive spectral counts at given stringency parameters<br>95_2_95<br>Gl50803_16596 | 95_2_50<br>Gl50803_16596 | InterProScan prediction | GO | HHPRED best hit (Probability E-value) |
|---|---|---|---|---|---|---|---|---|
| GL50803_16596 | Hypothetical protein | 895 | 1115 | 895 | 1115 | | | |
| GL50803_102108 | Clathrin heavy chain | 9 | 26 | 9 | | | | |
| GL50803_21423 | Beta adaptin | 10 | 17 | 10 | 17 | | | |
| GL50803_17304 | Alpha Adaptin | 3 | 8 | 3 | 8 | | | |
| GL50803_8917 | Mu adaptin | | 3 | | 3 | | | |
| GL50803_14373 | Dynamin (dynamin) | 23 | 31 | 23 | 31 | | | |
| GL50803_16811 | ZipA | 241 | 326 | 241 | 326 | | | |
| GL50803_9605 | hypothetical protein | 46 | 59 | 46 | 59 | P-loop containing nucleoside triphosphate | - | Antitoxin RelB3 5GOF_A (99.29/1e-12) |
| GL50803_41212 | hypothetical protein | 24 | 46 | 24 | 46 | - | - | Myosin 2 heavy chain striated 3JBH_B (99.95/3.6e-18) |
| GL50803_21628 | hypothetical protein | 14 | 23 | 14 | 23 | - | - | Antitoxin RelB3 3BPQ_A (80.35/4.7) |
| GL50803_9861 | hypothetical protein | 11 | 21 | | | - | - | Antitoxin RelB3 3BPQ_A (82.91/3.2) |
| GL50803_101278 | hypothetical protein | 6 | 12 | 6 | 12 | - | - | Shikimate kinase 2PT5_B (50.2/19) |
| GL50803_10882 | hypothetical protein | 6 | 8 | 6 | 8 | - | - | Vacuolar protein sorting-associated protein 75 5YPS_F |
| GL50803_113133 | hypothetical protein | 5 | 11 | 5 | 11 | ClpP/crotonase-like domain superfamily | - | Protein CT_858 3DOR_A (99.94/1.5e-27) |
| GL50803_17119 | hypothetical protein | 5 | 6 | 5 | 6 | Putative zinc-RING | - | Protein RUFY3 2DWK_A (86.65/6) |
| GL50803_103709 | hypothetical protein | 4 | 7 | 4 | 7 | BRO1 domain superfamily | GO:0005515 protein binding | BRO1 domain-containing protein 3R9M_A (98.55/2.9e-8) |
| GL50803_11720 | hypothetical protein | 4 | 7 | 4 | 7 | Ankyrin repeat-containing domain superfamily | GO:0005515 protein binding | Ankyrin 1N11_A (100/1.2e-32) |
| GL50803_16504 | hypothetical protein | 4 | 6 | 4 | 6 | - | - | Exocyst complex component EXO84 2D2S_A (72.8/20) |
| GL50803_17400 | hypothetical protein | 4 | 4 | 4 | 4 | Cyclin PHO80-like | GO:0000079 regulation of cyclin-dependent protein serine/threonine kinase activity GO:0019901 protein kinase | Cyclin-dependent protein kinase PHO85 4KRD_B (99.91/2.3e-6) |
| GL50803_94463 | hypothetical protein | 4 | 5 | 4 | 5 | - | - | Dimethylsulphoniopropionate lyase Ddd 4LA2_A (52.32/60) |
| GL50803_15499 | hypothetical protein | 3 | 4 | 3 | 4 | Tropomyosin | - | Smooth muscle myosin heavy chain 1I84_V (97.93/0.0021) |
| GL50803_115478 | hypothetical protein | | 7 | | 7 | Ankyrin repeat-containing domain superfamily | GO:0005515 protein binding | Ankyrin 1N11_A (100/7.8e-40) |
| GL50803_13584 | hypothetical protein | | 3 | | 3 | - | - | Smooth muscle myosin heavy chain 1I84_V (97.92/0.024) |
| GL50803_15251 | hypothetical protein | | 5 | | 5 | - | - | Nuclear pore complex protein Nup155 5IJO_Y (96.1/0.07) |
| GL50803_16717 | hypothetical protein | | 3 | | 3 | START-like domain superfamily | GO:0008289 lipid binding | StAR-related lipid transfer protein 3 5I9J_A (99.89/2.5e-25) |
| GL50803_32999 | hypothetical protein | | 4 | | 4 | SMC5, SMC6 | - | Smooth muscle myosin heavy chain 1I84_V (97.93/0.0021) |
| GL50803_4266 | hypothetical protein | | 3 | | 3 | - | - | Eukaryotic translation initiation factor 3 5A5T_A (98.99/0.000002) |
| GL50803_9183 | hypothetical protein | | 3 | | 3 | - | - | Pre-mRNA-processing-splicing factor 8 5XJC_V (26.44/350) |
| GL50803_7309 | Hypothetical protein | 2 | 3 | 2 | 3 | | | |
| GL50803_114776 | NSF | 3 | 11 | 3 | 11 | | | |
| GL50803_16453 | Carbamate kinase | | 14 | | | | | |
| GL50803_16034 | Kinase, CAMK CAMKL | 3 | 4 | 3 | 4 | | | |
| GL50803_16235 | Kinase, CAMK CAMKL | 4 | 6 | 4 | 6 | | | |
| GL50803_112076 | Kinase, CDC7 | | 5 | | 5 | | | |
| GL50803_17625 | Kinase, CMGC GSK | 3 | 4 | 3 | 4 | | | |
| GL50803_22850 | Kinase, CMGC MAPK | 4 | 10 | 4 | 10 | | | |
| GL50803_113553 | Kinase, NEK | 5 | 10 | 5 | 10 | | | |
| GL50803_11390 | Kinase, NEK | | 12 | | 12 | | | |
| GL50803_13215 | Kinase, NEK | 3 | 4 | 3 | 4 | | | |
| GL50803_15409 | Kinase, NEK | | 22 | | | | | |
| GL50803_16824 | Kinase, NEK | 3 | 5 | 3 | 5 | | | |
| GL50803_26199 | Kinase, NEK | 3 | 3 | 3 | 3 | | | |
| GL50803_5346 | Kinase, NEK | 3 | 3 | 3 | 3 | | | |
| GL50803_8445 | Kinase, NEK | | 4 | | 4 | | | |
| GL50803_87677 | Kinase, NEK | 34 | 6 | 34 | 50 | | | |
| GL50803_9421 | Kinase, NEK | 3 | 4 | 3 | 4 | | | |
| GL50803_15411 | Kinase, NEK-frag | 17 | 21 | | | | | |
| GL50803_15035 | Kinase, NEK-like | | 4 | | 4 | | | |
| GL50803_8805 | Kinase, SCY1 | 13 | 27 | 13 | 27 | | | |
| GL50803_90872 | Phosphoglycerate kinase | 5 | 6 | 5 | 6 | | | |
| GL50803_17143 | Pyruvate kinase | 8 | 9 | 8 | 9 | | | |
| GL50803_3206 | Pyruvate kinase | | 5 | | 5 | | | |
| GL50803_9909 | Pyruvate, phosphate dikinase | | 27 | | | | | |
| GL50803_16549 | Uridine kinase | 5 | 6 | 5 | 6 | | | |
| GL50803_8217 | Uridine kinase | 8 | 11 | 8 | 11 | | | |
| GL50803_10698 | Phosphorylase B kinase gamma catalytic chain | 5 | 8 | 5 | 8 | | | |
| GL50803_101291 | Beta tubulin | 26 | 36 | 26 | 36 | | | |
| GL50803_103676 | Alpha-tubulin | 17 | 21 | 17 | 21 | | | |
| GL50803_15097 | Alpha-14 giardin | | 5 | | 5 | | | |
| GL50803_14551 | Alpha-6 giardin | | 10 | | | | | |
| GL50803_103373 | Alpha-7.1 giardin | 3 | 21 | | | | | |
| GL50803_114787 | Alpha-7.3 giardin | 7 | 21 | 7 | 21 | | | |
| GL50803_17230 | Gamma giardin | 17 | 21 | 17 | 21 | | | |
| GL50803_14460 | Ubiquitin carboxyl-terminal hydrolase 4 | 3 | 3 | 3 | 3 | | | |
| GL50803_10661 | Ubiquitin-conjugating enzyme E1 | | 4 | | 4 | | | |
| GL50803_9779 | UPL-1 | | 6 | | 6 | | | |
| GL50803_10311 | Ornithine carbamoyltransferase | 68 | 85 | | | | | |
| GL50803_112103 | Arginine deiminase | 20 | 31 | | | | | |
| GL50803_93358 | Alcohol dehydrogenase | 20 | 26 | | | | | |
| GL50803_16867 | AAA family ATPase | 13 | 20 | 13 | 20 | | | |
| GL50803_17327 | Xaa-Pro dipeptidase | 15 | 19 | 15 | 19 | | | |
| GL50803_13608 | Acetyl-CoA synthetase | 14 | 18 | 14 | 18 | | | |
| GL50803_114609 | Pyruvate-flavodoxin oxidoreductase | 13 | 17 | | | | | |
| GL50803_15983 | Prolyl-tRNA synthetase | 14 | 17 | 14 | 17 | | | |
| GL50803_11118 | Enolase | 8 | 15 | 8 | 15 | | | |
| GL50803_21942 | NADP-specific glutamate dehydrogenase | 10 | 15 | 10 | 15 | | | |
| GL50803_7439 | Ser/Thr phosphatase 2A, 65kDa reg sub A | 12 | 14 | 12 | 14 | | | |
| GL50803_6226 | Glycogen phosphorylase | 5 | 13 | | | | | |
| GL50803_11043 | Fructose-bisphosphate aldolase | 10 | 12 | | | | | |
| GL50803_10570 | FKBP-type peptidyl-prolyl cis-trans isomerase | 6 | 11 | 6 | 11 | | | |
| GL50803_17063 | Pyruvate-flavodoxin oxidoreductase | 5 | 10 | 5 | 10 | | | |
| GL50803_15832 | Aminoacyl-histidine dipeptidase | 8 | 9 | 8 | 9 | | | |
| GL50803_3331 | Malate dehydrogenase | | 9 | | | | | |
| GL50803_6687 | Glyceraldehyde 3-phosphate dehydrogenase | 5 | 8 | 5 | 8 | | | |
| GL50803_86511 | Acyl-CoA synthetase | 5 | 7 | 5 | 7 | | | |
| GL50803_7532 | Vacuolar ATP synthase catalytic subunit A | 5 | 7 | 5 | 7 | | | |
| GL50803_16376 | ATP-dependent RNA helicase p47, putative | 4 | 6 | 4 | 6 | | | |
| GL50803_21118 | Long chain fatty acid CoA ligase 5 | | 6 | | 6 | | | |
| GL50803_113892 | Long chain fatty acid CoA ligase, putative | | 6 | | | | | |
| GL50803_9413 | PDI2 - Protein disulfide isomerase | 4 | 6 | 4 | 6 | | | |
| GL50803_93548 | Phospholipase B | 4 | 6 | 4 | 6 | | | |
| GL50803_10870 | Aspartyl-tRNA synthetase | 4 | 5 | 4 | 5 | | | |
| GL50803_9115 | Glucose-6-phosphate isomerase | 3 | 5 | 3 | 5 | | | |
| GL50803_16766 | Lysyl-tRNA synthetase | 3 | 5 | 3 | 5 | | | |
| GL50803_17254 | Phosphomannomutase-2 | 3 | 5 | 3 | 5 | | | |
| GL50803_16443 | Protein phosphatase 2A B' regulatory subunit Wdb1 | 3 | 5 | 3 | 5 | | | |
| GL50803_15214 | Ser/Thr phosphatase | 4 | 5 | 4 | 5 | | | |
| GL50803_101501 | Seryl-tRNA synthetase | 3 | 5 | 3 | 5 | | | |
| GL50803_12216 | Vacuolar ATP synthase subunit B | 5 | 5 | 5 | 5 | | | |
| GL50803_14961 | Vacuolar ATP synthase subunit H | | 5 | | 5 | | | |
| GL50803_8822 | 2,3-bisphosphoglycerate-independen phosphoglycerate mutas | | 4 | | 4 | | | |
| GL50803_9011 | Glycyl-tRNA synthetase | 4 | 4 | 4 | 4 | | | |
| GL50803_104173 | Isoleucyl-tRNA synthetase | 3 | 4 | 3 | 4 | | | |
| GL50803_8621 | Leucyl-tRNA synthetase | 3 | 4 | 3 | 4 | | | |
| GL50803_103713 | PDI4 - Protein disulfide isomerase | 3 | 4 | 3 | 4 | | | |
| GL50803_15215 | Phosphatase | 3 | 4 | 3 | 4 | | | |
| GL50803_10623 | Phosphoenolpyruvate carboxykinase | | 4 | | 4 | | | |
| GL50803_3643 | 70 kDa peptidylprolyl isomerase, putative | | 3 | | 3 | | | |
| GL50803_96460 | Alanyl-tRNA synthetase | | 3 | | 3 | | | |
| GL50803_3861 | Alcohol dehydrogenase 3 lateral transfer candidate | | 3 | | | | | |
| GL50803_3593 | Alcohol dehydrogenase lateral transfer candidate | | 3 | | 3 | | | |
| GL50803_14375 | Asparaginyl-tRNA synthetase | | 3 | | 3 | | | |
| GL50803_8826 | Glucokinase | 3 | 3 | 3 | 3 | | | |
| GL50803_16125 | Glycerol-3-phosphate dehydrogenase | | 3 | | | | | |
| GL50803_14285 | Malic enzyme | | 3 | | 3 | | | |
| GL50803_32312 | Protein phosphatase 2C | | 3 | | 3 | | | |
| GL50803_16567 | Protein phosphatase PP2A regulatory subunit A | 3 | 3 | 3 | 3 | | | |
| GL50803_14993 | Pyrophosphate-fructose 6-phosphate 1 phosphotransferase alpha subun | 3 | 3 | 3 | 3 | | | |
| GL50803_16795 | Topoisomerase II (VPS4b) | | 5 | | 5 | | | |
| GL50803_9704 | Transketolase | 6 | 7 | 6 | 7 | | | |
| GL50803_9508 | Metalloprotease, insulinase family | | 3 | | 3 | | | |
| GL50803_33769 | NADH oxidase lateral transfer candidate | 3 | 5 | 3 | 5 | | | |
| GL50803_21942 | NADP-specific glutamate dehydrogenase | | 15 | | | | | |
| GL50803_16525 | Ribosomal protein L3 | 7 | 11 | 7 | 11 | | | |
| GL50803_17547 | Ribosomal protein L4 | 22 | 34 | 22 | | | | |
| GL50803_17395 | Ribosomal protein L5 | 5 | 10 | 5 | 10 | | | |
| GL50803_19436 | Ribosomal protein L7 | | 3 | | | | | |
| GL50803_10367 | Ribosomal protein S24 | 3 | 3 | 3 | 3 | | | |
| GL50803_16265 | Ribosomal protein S3a | | 3 | | | | | |
| GL50803_14620 | Ribosomal protein S6 | | 6 | | | | | |
| GL50803_102813 | Protein 21.1 | | 3 | | 3 | | | |
| GL50803_12139 | Protein 21.1 | | 5 | | 5 | | | |
| GL50803_14859 | Protein 21.1 | 11 | 19 | 11 | 19 | | | |
| GL50803_16220 | Protein 21.1 | 4 | 9 | 4 | 9 | | | |
| GL50803_17060 | Protein 21.1 | 23 | 30 | 23 | 30 | | | |
| GL50803_17568 | Protein 21.1 | 3 | 3 | 3 | 3 | | | |
| GL50803_17586 | Protein 21.1 | | 3 | | 3 | | | |
| GL50803_27925 | Protein 21.1 | 30 | 46 | 30 | 46 | | | |
| GL50803_32778 | Protein 21.1 | 4 | 8 | 4 | 8 | | | |
| GL50803_4383 | Protein 21.1 | | 5 | | 5 | | | |
| GL50803_9030 | Protein 21.1 | 22 | 26 | | | | | |
| GL50803_17054 | Acidic ribosomal protein P0 | 18 | 25 | | | | | |
| GL50803_10370 | ATP/GTP binding protein, putative | | 5 | | 5 | | | |
| GL50803_137716 | Axoneme-associated protein GASP-180 | 82 | 118 | | | | | |
| GL50803_16745 | Axoneme-associated protein GASP-180 | 18 | 24 | | | | | |
| GL50803_17121 | Bip (BiP) | | 55 | | | | | |
| GL50803_9808 | Chaperone protein DnaJ subfamily A | 5 | 9 | 5 | 9 | | | |
| GL50803_15148 | Chaperone protein DnaJ subfamily C | 3 | 4 | 3 | 4 | | | |
| GL50803_10167 | Coiled-coil protein | 8 | 13 | 8 | 13 | | | |
| GL50803_113677 | Coiled-coil protein | 9 | 13 | 9 | 13 | | | |
| GL50803_15591 | Coiled-coil protein | | 3 | | 3 | | | |
| GL50803_17249 | Coiled-coil protein | 15 | 27 | | | | | |
| GL50803_88765 | Cytosolic HSP70 | 63 | 87 | 63 | 87 | | | |
| GL50803_16200 | Developmentally regulated GTP-binding protein 1 | | 3 | | 3 | | | |
| GL50803_103059 | Dynein heavy chain | | 6 | | 6 | | | |
| GL50803_94440 | Dynein heavy chain | | 3 | | | | | |
| GL50803_112304 | Elongation factor 1-alpha | | 61 | | | | | |
| GL50803_12102 | Elongation factor 1-gamma | 7 | 15 | 7 | 15 | | | |
| GL50803_17570 | Elongation factor 2 | 30 | 42 | 30 | 42 | | | |
| GL50803_97219 | Fibrillarin-like pre-rRNA processing protein Narcis et al | 3 | 5 | 3 | 5 | | | |
| GL50803_13864 | Heat shock protein HSP 90-alpha | 10 | 19 | 10 | 19 | | | |
| GL50803_98054 | Heat shock protein HSP 90-alpha | 17 | 28 | 17 | 28 | | | |
| GL50803_16412 | Heat-shock protein, putative | | 3 | | 3 | | | |
| GL50803_21321 | High cysteine membrane protein Group 5 | 3 | 5 | 3 | 5 | | | |
| GL50803_15106 | Importin beta-3 subunit | 5 | 8 | 5 | 8 | | | |
| GL50803_15054 | Kelch repeat-containing protein | 3 | 3 | 3 | 3 | | | |
| GL50803_102101 | Kinesin-3 | | 6 | | | | | |
| GL50803_112846 | Kinesin-3 | | 6 | | 6 | | | |
| GL50803_16343 | Median body protein | 5 | 8 | 5 | 8 | | | |
| GL50803_10577 | Nucleolar protein NOP5 | 3 | 4 | 3 | 4 | | | |
| GL50803_10969 | Nucleotide-binding protein 1 | 4 | 5 | 4 | 5 | | | |
| GL50803_4197 | Phosphatidylinositol transfer protein alpha isoform | 3 | 4 | 3 | 4 | | | |
| GL50803_5942 | Polyadenylate-binding protein, putative | 7 | 10 | 7 | 10 | | | |
| GL50803_4410 | SALP-1 | | 3 | | 3 | | | |
| GL50803_21444 | Spindle pole protein, putative | 4 | 5 | 4 | 5 | | | |
| GL50803_24537 | Spindle pole protein, putative | 3 | 3 | 3 | 3 | | | |
| GL50803_33660 | Spindle pole protein, putative | | 3 | | 3 | | | |
| GL50803_7031 | Spindle pole protein, putative | 4 | 7 | 4 | 7 | | | |
| GL50803_17565 | TBP-interacting protein TIP49 | 3 | 4 | 3 | 4 | | | |
| GL50803_9825 | TBP-interacting protein TIP49 | 3 | 4 | 3 | 4 | | | |
| GL50803_11397 | TCP-1 chaperonin subunit beta | 4 | 6 | | | | | |
| GL50803_11992 | TCP-1 chaperonin subunit epsilon | 5 | 5 | 5 | 5 | | | |
| GL50803_16124 | TCP-1 chaperonin subunit eta | 6 | 7 | 6 | 7 | | | |
| GL50803_13500 | TCP-1 chaperonin subunit theta | 8 | 9 | 8 | 9 | | | |
| GL50803_10231 | TCP-1 chaperonin subunit zeta | 3 | 3 | | | | | |
| GL50803_10255 | Translation initiation factor eIF-4A, putative | 13 | 17 | 13 | 17 | | | |
| GL50803_23833 | Vacuolar protein sorting 35 (VPS35) | | 3 | | 3 | | | |

| Supplementary Table 4 GL50803_42357 *Gl* PXD4 | | | | | | | | |
|---|---|---|---|---|---|---|---|---|
| | | bait specific spectral counts at given stringency parameters | | bait specific exclusive spectral counts at given stringency parameters | | InterProScan prediction | GO | HHPRED best hit (Probability E-value) |
| ORF number | Annotation | 95_2_95 Gl50803_42357 | 95_2_50 Gl50803_42357 | 95_2_95 Gl50803_42357 | 95_2_50 Gl50803_42357 | | | |
| GL50803_42357 | hypothetical protein | 37 | 78 | 37 | 78 | | | |
| GL50803_102108 | Clathrin heavy chain | 6 | 24 | 6 | | | | |
| GL50803_21423 | Beta Adaptin | | 4 | | 4 | | | |
| GL50803_14373 | Dynamin | 4 | 5 | 4 | 5 | | | |
| GL50803_16595 | Liver stage antigen-like protein | 2 | 4 | 2 | 4 | | | |
| GL50803_17037 | hypothetical protein | 10 | 17 | 10 | 17 | - | - | Nuclear pore complex protein Nup155 5IJN_T (75.53/96) |
| GL50803_9861 | hypothetical protein | | 7 | | | - | - | Antitoxin RelB3 3BPQ_A (82.91/3.2) |
| GL50803_41212 | hypothetical protein | | 3 | | 3 | - | - | Myosin 2 heavy chain 3JBH_B (99.95/3.6e-18) |
| GL50803_9183 | hypothetical protein | | 3 | | 3 | - | - | Pre-mRNA-processing-splicing factor 8 5XJC_V (26.44/350) |
| GL50803_11654 | Alpha-1 giardin | | 34 | | | | | |
| GL50803_7796 | Alpha-2 giardin | 38 | 46 | | | | | |
| GL50803_14551 | Alpha-6 giardin | | 7 | | | | | |
| GL50803_114787 | Alpha-7.3 giardin | 5 | | 5 | | | | |
| GL50803_101291 | Beta tubulin | | 4 | | 4 | | | |
| GL50803_17230 | Gamma giardin | | 6 | | 6 | | | |
| GL50803_112103 | Arginine deiminase | | 31 | | | | | |
| GL50803_114609 | Pyruvate-flavodoxin oxidoreductase | | 10 | | | | | |
| GL50803_17063 | Pyruvate-flavodoxin oxidoreductase | | 6 | | 6 | | | |
| GL50803_15215 | Phosphatase | | 4 | | 4 | | | |
| GL50803_7999 | Ribosomal protein S3 | 11 | 16 | 11 | 16 | | | |
| GL50803_14622 | Ribosomal protein L13 | 10 | 11 | 10 | 11 | | | |
| GL50803_1345 | Ribosomal protein L10a | 9 | 12 | 9 | 12 | | | |
| GL50803_16431 | Ribosomal protein L19 | 9 | 13 | 9 | 13 | | | |
| GL50803_10428 | Ribosomal protein L10 | 7 | 12 | 7 | 12 | | | |
| GL50803_16387 | Ribosomal protein L18a | 6 | 11 | 6 | 11 | | | |
| GL50803_16086 | Ribosomal protein L2 | 6 | 8 | 6 | 8 | | | |
| GL50803_16525 | Ribosomal protein L3 | 5 | 7 | 5 | 7 | | | |
| GL50803_17395 | Ribosomal protein L5 | 5 | 11 | 5 | 11 | | | |
| GL50803_14827 | Ribosomal protein S11 | 5 | 7 | 5 | 7 | | | |
| GL50803_14329 | Ribosomal protein S7 | 5 | 8 | 5 | 8 | | | |
| GL50803_7766 | Ribosomal protein SA | 4 | 5 | 4 | 5 | | | |
| GL50803_14938 | Ribosomal protein L12 | 3 | 6 | 3 | 6 | | | |
| GL50803_11247 | Ribosomal protein L13a | | 18 | | | | | |
| GL50803_8001 | Ribosomal protein L15 | | 3 | | 3 | | | |
| GL50803_11950 | Ribosomal protein L18 | | 3 | | 3 | | | |
| GL50803_6133 | Ribosomal protein L35 | | 3 | | 3 | | | |
| GL50803_17547 | Ribosomal protein L4 | | 17 | | | | | |
| GL50803_19436 | Ribosomal protein L7 | | 27 | | | | | |
| GL50803_17244 | Ribosomal protein L7a | | 24 | | | | | |
| GL50803_17056 | Ribosomal protein L9 | | 3 | | 3 | | | |
| GL50803_17054 | Acidic ribosomal protein P0 | | 5 | | | | | |
| GL50803_137716 | Axoneme-associated protein GASP-180 | | 33 | | | | | |
| GL50803_113677 | Coiled-coil protein | 7 | 19 | 7 | 19 | | | |
| GL50803_15591 | Coiled-coil protein | 40 | 90 | 40 | 90 | | | |
| GL50803_88765 | Cytosolic HSP70 | 3 | 6 | 3 | 6 | | | |
| GL50803_17265 | Dynein heavy chain | 3 | 5 | 3 | 5 | | | |
| GL50803_111950 | Dynein heavy chain | | 3 | | 3 | | | |
| GL50803_112304 | Elongation factor 1-alpha | 7 | 37 | | | | | |
| GL50803_15427 | Giardia trophozoite antigen GTA-2 | 3 | 5 | 3 | 5 | | | |
| GL50803_14521 | Peroxiredoxin 1 | 3 | | | | | | |

### Supplementary Table 5 GL50803_24488 *Gl* PXD6

| | | bait specific spectral counts at given stringency parameters | | bait specific exclusive spectral counts at given stringency parameters | | InterProScan prediction | GO | HHPRED best hit (Probability E-value) |
|---|---|---|---|---|---|---|---|---|
| ORF number | Annotation | 95_2_95 GL50803_24488 | 95_2_50 Gl50803_24488 | 95_2_95 Gl50803_24488 | 95_2_50 Gl50803_24488 | | | |
| GL50803_24488 | hypothetical protein | 30 | 57 | 30 | 57 | | | |
| GL50803_102108 | Clathrin heavy chain | 8 | 13 | 8 | | | | |
| GL50803_21423 | Beta Adaptin | 6 | 8 | 6 | 8 | | | |
| GL50803_14373 | Dynamin (dynamin) | 8 | 16 | 8 | 16 | | | |
| GL50803_16596 | hypothetical protein | | 4 | | 4 | | | |
| GL50803_17332 | hypothetical protein | 22 | 33 | | | WD40-repeat-containing domain superfamily | GO:0005515 protein binding | serine/threonine-protein kinase 5YZV_A (95.38/0.55) |
| GL50803_41212 | hypothetical protein | 7 | 16 | 7 | 16 | - | - | Myosin 2 heavy chain 3JBH_B (99.95/3.6e-18) |
| GL50803_21628 | hypothetical protein | 5 | 13 | 5 | 13 | - | - | Antitoxin RelB3 3BPQ_A (80.35/4.7) |
| GL50803_9861 | hypothetical protein | 5 | 10 | | | - | - | Antitoxin RelB3 3BPQ_A (82.91/3.2) |
| GL50803_4266 | hypothetical protein | 4 | 4 | 4 | 4 | - | - | Eukaryotic translation initiation factor 3 5A5T_A (98.99/0.000002) |
| GL50803_113133 | hypothetical protein | | 3 | | 3 | ClpP/crotonase-like domain superfamily | - | Protein CT_858 3DOR_A (99.94/1.5e-27) |
| GL50803_11720 | hypothetical protein | | 3 | | 3 | Ankyrin repeat-containing domain superfamily | GO:0005515 protein binding | Ankyrin 1N11_A (100/1.2e-32) |
| GL50803_16653 | hypothetical protein | | 3 | | 3 | Zinc finger, RING/FYVE/PHD-type | GO:0046872 metal ion binding | Zinc finger FYVE domain-containing protein |
| GL50803_16717 | hypothetical protein | 3 | 3 | 3 | 3 | START-like domain superfamily | GO:0008289 lipid binding | StAR-related lipid transfer protein 3 5I9J_A (99.89/2.5e-25) |
| GL50803_7204 | hypothetical protein | | 3 | | 3 | RNA-binding superdomain | O:0003676 nucleic acid binding | KIAA0430 protein 2DIU_A (96.43/0.00018) |
| GL50803_114776 | NSF | | 11 | | 3 | | | |
| GL50803_103676 | Alpha-tubulin | | 5 | | 5 | | | |
| GL50803_101291 | Beta tubulin | 4 | 7 | 4 | 7 | | | |
| GL50803_11654 | Alpha-1 giardin | | 16 | | | | | |
| GL50803_114787 | Alpha-7.3 giardin | | 4 | | | | | |
| GL50803_16867 | AAA family ATPase | 7 | 7 | 7 | 7 | | | |
| GL50803_13608 | Acetyl-CoA synthetase | 7 | 9 | 7 | 9 | | | |
| GL50803_86511 | Acyl-CoA synthetase | | 3 | | 3 | | | |
| GL50803_93358 | Alcohol dehydrogenase | 11 | 17 | | | | | |
| GL50803_112103 | Arginine deiminase | 8 | 13 | | | | | |
| GL50803_10521 | Arginyl-tRNA synthetase | | 3 | | 3 | | | |
| GL50803_11118 | Enolase | 3 | 9 | 3 | 9 | | | |
| GL50803_8682 | Glucose-6-phosphate 1-dehydrogenase | | 3 | | 3 | | | |
| GL50803_6226 | Glycogen phosphorylase | 3 | | | | | | |
| GL50803_14285 | Malic enzyme | | 3 | | 3 | | | |
| GL50803_9719 | NADH oxidase | | 3 | | 3 | | | |
| GL50803_33769 | NADH oxidase lateral transfer candidate | | 4 | | | | | |
| GL50803_21942 | NADP-specific glutamate dehydrogenase | 4 | 9 | 4 | 9 | | | |
| GL50803_9413 | PDI2 - Protein disulfide isomerase | | 5 | | 5 | | | |
| GL50803_8064 | PDI5 - Protein disulfide isomerase | | 3 | | 3 | | | |
| GL50803_93548 | Phospholipase B | | 3 | | 3 | | | |
| GL50803_17254 | Phosphomannomutase-2 | | 3 | | 3 | | | |
| GL50803_10698 | Phosphorylase B kinase gamma catalytic chain | | 3 | | 3 | | | |
| GL50803_15983 | Prolyl-tRNA synthetase | 5 | 5 | 5 | 5 | | | |
| GL50803_114609 | Pyruvate-flavodoxin oxidoreductase | 4 | 5 | | | | | |
| GL50803_17063 | Pyruvate-flavodoxin oxidoreductase | 4 | 7 | 4 | 7 | | | |
| GL50803_7439 | Ser/Thr phosphatase 2A, 65kDa reg sub A | 5 | 6 | 5 | 6 | | | |
| GL50803_9704 | Transketolase | 4 | 6 | 4 | 6 | | | |
| GL50803_7532 | Vacuolar ATP synthase catalytic subunit A | | 6 | | 6 | | | |
| GL50803_17327 | Xaa-Pro dipeptidase | 3 | 8 | 3 | 8 | | | |
| GL50803_16034 | Kinase, CAMK CAMKL | 3 | 3 | 3 | 3 | | | |
| GL50803_15409 | Kinase, NEK | 11 | 12 | | | | | |
| GL50803_15411 | Kinase, NEK-frag | 8 | 12 | | | | | |
| GL50803_102034 | Kinase, NEK-frag | | 3 | | 3 | | | |
| GL50803_10609 | Kinase, STE STE20 | | 3 | | 3 | | | |
| GL50803_90872 | Phosphoglycerate kinase | | 3 | | 3 | | | |
| GL50803_17143 | Pyruvate kinase | | 5 | | 5 | | | |
| GL50803_1345 | Ribosomal protein L10a | 3 | 3 | 3 | 3 | | | |
| GL50803_16387 | Ribosomal protein L18a | | 4 | | 4 | | | |
| GL50803_10091 | Ribosomal protein L23 | 3 | 3 | 3 | 3 | | | |
| GL50803_7870 | Ribosomal protein L23A | 3 | 5 | 3 | 5 | | | |
| GL50803_16310 | Ribosomal protein L27a | 3 | 4 | 3 | 4 | | | |
| GL50803_16525 | Ribosomal protein L3 | 3 | 4 | 3 | 4 | | | |
| GL50803_6133 | Ribosomal protein L35 | 3 | 3 | 3 | 3 | | | |
| GL50803_7878 | Ribosomal protein S14 | 6 | 6 | | | | | |
| GL50803_15260 | Ribosomal protein S15 | | 5 | | 5 | | | |
| GL50803_6135 | Ribosomal protein S17 | | 3 | | 3 | | | |
| GL50803_15551 | Ribosomal protein S18 | | 3 | | 3 | | | |
| GL50803_10367 | Ribosomal protein S24 | 4 | 5 | 4 | 5 | | | |
| GL50803_12981 | Ribosomal protein S5 | | 5 | | 5 | | | |
| GL50803_14329 | Ribosomal protein S7 | | 3 | | 3 | | | |
| GL50803_4547 | Ribosomal protein S9 | | 3 | | 3 | | | |
| GL50803_27925 | Protein 21.1 | 10 | 23 | 10 | 23 | | | |
| GL50803_17060 | Protein 21.1 | 6 | 12 | 6 | 12 | | | |
| GL50803_14859 | Protein 21.1 | 7 | 9 | 7 | 9 | | | |
| GL50803_21505 | Protein 21.1 | 4 | 4 | 4 | 4 | | | |
| GL50803_9030 | Protein 21.1 | | 11 | | | | | |
| GL50803_32778 | Protein 21.1 | | 3 | | 3 | | | |
| GL50803_17054 | Acidic ribosomal protein P0 | 8 | 12 | | | | | |
| GL50803_137716 | Axoneme-associated protein GASP-180 | 3 | 33 | | | | | |
| GL50803_17121 | Bip (BiP) | 29 | 36 | | | | | |
| GL50803_9808 | Chaperone protein DnaJ subfamily A | | 3 | | 3 | | | |
| GL50803_15148 | Chaperone protein DnaJ subfamily C | | 3 | | 3 | | | |
| GL50803_10167 | Coiled-coil protein | 8 | 12 | 8 | 12 | | | |
| GL50803_15591 | Coiled-coil protein | | 3 | | 3 | | | |
| GL50803_17249 | Coiled-coil protein | 11 | 15 | | | | | |
| GL50803_88765 | Cytosolic HSP70 | 21 | 35 | 21 | 35 | | | |
| GL50803_112304 | Elongation factor 1-alpha | 30 | 37 | | | | | |
| GL50803_12102 | Elongation factor 1-gamma | 3 | 4 | 3 | 4 | | | |
| GL50803_17570 | Elongation factor 2 | 8 | 19 | 8 | 19 | | | |
| GL50803_97219 | Fibrillarin-like pre-rRNA processing protein Narcisi et al | | 3 | | 3 | | | |
| GL50803_17090 | Giardia trophozoite antigen GTA-1 | | 3 | | 3 | | | |
| GL50803_98054 | Heat shock protein HSP 90-alpha | 7 | 10 | 7 | 10 | | | |
| GL50803_13864 | Heat shock protein HSP 90-alpha | 3 | 4 | 3 | 4 | | | |
| GL50803_21321 | High cysteine membrane protein Group 5 | | 4 | | 4 | | | |
| GL50803_121045 | Histone H2B | | 5 | | | | | |
| GL50803_135231 | Histone H3 | | 3 | | 3 | | | |
| GL50803_15106 | Importin beta-3 subunit | | 3 | | 3 | | | |
| GL50803_5795 | Leucine-rich repeat protein 1 virus receptor protein | 3 | | | | | | |
| GL50803_33769 | NADH oxidase lateral transfer candidate | 3 | 4 | 3 | 4 | | | |
| GL50803_17163 | Peptidyl-prolyl cis-trans isomerase B precursor | | 4 | | 4 | | | |
| GL50803_14521 | Peroxiredoxin 1 | 10 | | | | | | |
| GL50803_7031 | Spindle pole protein, putative | | 5 | | 5 | | | |
| GL50803_17565 | TBP-interacting protein TIP49 | 3 | 3 | 3 | 3 | | | |
| GL50803_11992 | TCP-1 chaperonin subunit epsilon | | 5 | | 5 | | | |
| GL50803_16124 | TCP-1 chaperonin subunit eta | 3 | 3 | 3 | 3 | | | |
| GL50803_13500 | TCP-1 chaperonin subunit theta | | 3 | | 3 | | | |
| GL50803_10255 | Translation initiation factor eIF-4A, putative | 3 | 4 | 3 | 4 | | | |
| GL50803_23833 | Vacuolar protein sorting 35 (VPS35) | | 7 | | 7 | | | |

| Supplementary Table 6 GL50803_16653 *Gl* FYVE | | | | | | | | |
|---|---|---|---|---|---|---|---|---|
| | | bait specific spectral counts at given stringency parameters | | bait specific exclusive spectral counts at given stringency parameters | | InterProScan prediction | GO | HHPRED best hit (Probability E-value) |
| ORF number | Annotation | 95_2_95 Gl50803_16653 | 95_2_50 Gl50803_16653 | 95_2_95 Gl50803_16653 | 95_2_50 Gl50803_16653 | | | |
| GL50803_16653 | hypothetical protein | 17 | 29 | 17 | 29 | | | |
| GL50803_102108 | Clathrin heavy chain | 4 | 8 | 4 | | | | |
| GL50803_14373 | Dynamin | 10 | 21 | 10 | 21 | | | |
| GL50803_17332 | hypothetical protein | 4 | 8 | | | WD40-repeat-containing domain superfamily | GO:0005515 protein binding | serine/threonine-protein kinase 5YZV_A (95.38/0.55) |
| GL50803_4595 | hypothetical protein | 4 | 4 | 4 | 4 | - | - | Pleurotolysin A; beta-sandwich fold 4OEB_C(51.13/26) |
| GL50803_23447 | hypothetical protein | 3 | 7 | 3 | 7 | - | - | CARDB domain, PFAM 07705 2M8X_ A (50.46/56) |
| GL50803_16717 | hypothetical protein | 3 | 3 | 3 | 3 | START-like domain superfamily | GO:0008289 lipid binding | StAR-related lipid transfer protein 3 5I9J_A (99.89/2.5e-25) |
| GL50803_10778 | hypothetical protein | | 6 | | 6 | - | - | DNA-directed RNA polymerase II subunit 3H0G_M (77.74/41) |
| GL50803_9183 | hypothetical protein | | 3 | | 3 | - | - | Pre-mRNA-processing-splicing factor 8 5XJC_V (26.44/350) |
| GL50803_112811 | hypothetical protein | | 3 | | 3 | Protein-tyrosine phosphatase-like/Myotubularin family | - | Myotubularin-related protein 2 (E.C.3.1.3.64) 1LW3_A (100/4.1e-87) |
| GL50803_10822 | WD-40 repeat protein family | **5** | 7 | 5 | 7 | | | |
| GL50803_114776 | NSF | 3 | 3 | 3 | 3 | | | |
| GL50803_11654 | Alpha-1 giardin | 7 | 10 | | | | | |
| GL50803_4812 | Beta-giardin | 7 | 11 | | | | | |
| GL50803_16867 | AAA family ATPase | 3 | 5 | 3 | 5 | | | |
| GL50803_13608 | Acetyl-CoA synthetase | | 3 | | 3 | | | |
| GL50803_86511 | Acyl-CoA synthetase | | 3 | | | | | |
| GL50803_93358 | Alcohol dehydrogenase | 4 | 5 | | | | | |
| GL50803_112103 | Arginine deiminase | | 5 | | | | | |
| GL50803_11118 | Enolase | | 4 | | 4 | | | |
| GL50803_11043 | Fructose-bisphosphate aldolase | | 6 | | | | | |
| GL50803_10311 | Ornithine carbamoyltransferase | 5 | 7 | | | | | |
| GL50803_17163 | Peptidyl-prolyl cis-trans isomerase B precursor | | 5 | | 5 | | | |
| GL50803_13875 | Phosphatase | 4 | 4 | 4 | 4 | | | |
| GL50803_93548 | Phospholipase B | | 3 | | 3 | | | |
| GL50803_10698 | Phosphorylase B kinase gamma catalytic chain | | 3 | | 3 | | | |
| GL50803_16443 | Protein phosphatase 2A B' regulatory subunit Wdb1 | | 3 | | 3 | | | |
| GL50803_17063 | Pyruvate-flavodoxin oxidoreductase | | 3 | | 3 | | | |
| GL50803_114609 | Pyruvate-flavodoxin oxidoreductase | | 3 | | | | | |
| GL50803_9909 | Pyruvate, phosphate dikinase | 3 | 7 | | | | | |
| GL50803_9704 | Transketolase | 3 | 4 | 3 | 4 | | | |
| GL50803_7532 | Vacuolar ATP synthase catalytic subunit A | 3 | 4 | 3 | 4 | | | |
| GL50803_17327 | Xaa-Pro peptidase | | 6 | | 6 | | | |
| GL50803_11390 | Kinase, NEK | 7 | 14 | 7 | 14 | | | |
| GL50803_15409 | Kinase, NEK | | 4 | | | | | |
| GL50803_8805 | Kinase, SCY1 | 3 | 7 | 3 | 7 | | | |
| GL50803_17244 | Ribosomal protein L7a | 3 | 4 | | | | | |
| GL50803_9030 | Protein 21.1 | 3 | 5 | | | | | |
| GL50803_21505 | Protein 21.1 | | 4 | | 4 | | | |
| GL50803_17060 | Protein 21.1 | | 3 | | 3 | | | |
| GL50803_6430 | 14-3-3 protein | 10 | 13 | 10 | 13 | | | |
| GL50803_137716 | Axoneme-associated protein GASP-180 | | 3 | | | | | |
| GL50803_17121 | Bip | 4 | 5 | | | | | |
| GL50803_9808 | Chaperone protein DnaJ | | 3 | | 3 | | | |
| GL50803_113677 | Coiled-coil protein | | 3 | | 3 | | | |
| GL50803_88765 | Cytosolic HSP70 | 3 | 4 | 3 | 4 | | | |
| GL50803_112304 | Elongation factor 1-alpha | | 6 | | | | | |
| GL50803_17570 | Elongation factor 2 | 3 | 3 | 3 | 3 | | | |
| GL50803_14521 | Peroxiredoxin 1 | 7 | 8 | | | | | |
| GL50803_103855 | RNase L inhibitor | | 4 | | 4 | | | |
| GL50803_9376 | Sec 23 | | 3 | | 3 | | | |
| GL50803_23833 | Vacuolar protein sorting 35 | 9 | 15 | 9 | 15 | | | |

| Supplementary Table 7 GL50803_17195G/NECAP1 | | | | | | | | |
|---|---|---|---|---|---|---|---|---|
| | | bait specific spectral counts at given stringency parameters | | bait specific exclusive spectral counts at given stringency parameters | | InterProScan prediction | GO | HHPRED best hit (Probability E-value) |
| ORF number | Annotation | 95_2_95 GI50803_17195 | 95_2_50 GI50803_17195 | 95_2_95 GI50803_17195 | 95_2_50 GI50803_17195 | | | |
| GL50803_17195 | hypothetical protein | 93 | 175 | 93 | 175 | | | |
| GL50803_102108 | Clathrin heavy chain | 26 | 35 | 26 | | | | |
| GL50803_21423 | Beta adaptin | 31 | 40 | 31 | 40 | | | |
| GL50803_17304 | Alpha Adaptin | 14 | 19 | 14 | 19 | | | |
| GL50803_8917 | Mu adaptin | 8 | 12 | 8 | 12 | | | |
| GL50803_14373 | Dynamin (dynamin) | 27 | 42 | 27 | 42 | | | |
| GL50803_16653 | hypothetical protein | 7 | 19 | 7 | 19 | | | |
| GL50803_101278 | Hypothetical protein | 31 | 34 | 31 | 34 | - | - | Shikimate kinase 2PT5_B (50.2/19) |
| GL50803_17332 | Hypothetical protein | 28 | 34 | | | WD40-repeat-containing domain superfamily | GO:0005515 protein binding | serine/threonine-protein kinase 5YZV_A (95.38/0.55) |
| GL50803_94463 | Hypothetical protein | 25 | 32 | 25 | 32 | - | - | Dimethylsulphoniopropionate lyase DddQ 4LA2_A (52.32/60) |
| GL50803_103709 | Hypothetical protein | 21 | 26 | 21 | 26 | BRO1 domain superfamily | GO:0005515 protein binding | BRO1 domain-containing protein 3R9M_A (98.55/2.9e-8) |
| GL50803_9183 | Hypothetical protein | 19 | 29 | 19 | 29 | - | - | Pre-mRNA-processing-splicing factor 8 5XJC_V (26.44/350) |
| GL50803_16504 | Hypothetical protein | 16 | 18 | 16 | 18 | - | - | Exocyst complex component EXO84 2D2S_A (72.8/20) |
| GL50803_8528 | Hypothetical protein | 11 | 19 | 11 | 19 | - | - | Replication protein A 14 kDa 3KDF_C (79.97/17) |
| GL50803_41212 | Hypothetical protein | 9 | 12 | 9 | 12 | - | - | Myosin 2 heavy chain 3JBH_B (99.95/3.6e-18) |
| GL50803_11354 | Hypothetical protein | 8 | 9 | 8 | 9 | - | - | Toxic extracellular endopeptidase (E.C.3.4.24.39) 2X3C_A (94.97/0.07) |
| GL50803_17119 | Hypothetical protein | 8 | 12 | 8 | 12 | Putative zinc-RING | - | Protein RUFY3 2DWK_A (86.65/6) |
| GL50803_11380 | Hypothetical protein | 8 | 11 | 8 | 11 | - | - | Pre-mRNA-splicing factor 8 6BK8_A (93.9/0.04) |
| GL50803_8405 | Hypothetical protein | 8 | 10 | 8 | 10 | Zinc finger C2HC domain-containing protein | - | Transcriptional repressor CTCF/DNA Complex 5YEL_A (99.24/6.2e-14) |
| GL50803_5810 | Hypothetical protein | 6 | 7 | 6 | 7 | FMN-binding split barrel, Pyridoxamine 5' phosphate oxidase, putative | GO:0048037 cofactor binding | putative general stress protein 2FHQ_A (99.85/2.3e-22) |
| GL50803_6464 | Hypothetical protein | 5 | 5 | 5 | 5 | Myosin heavy related/Crossover suppressor | - | Tyrosine-protein phosphatase non-receptor type 23 5MK2_B (98.25/0.000002) |
| GL50803_6933 | hypothetical protein | 5 | 6 | 5 | 6 | Armadillo-like helical | - | Exportin-5/13-mer peptide/GTP-binding nuclear protein Ran/RNA 3A6P_F (99.04/3.7e-7) |
| GL50803_10881 | Hypothetical protein | 5 | 8 | 5 | 8 | WD40/YVTN repeat-like-containing domain superfamily | GO:00005515 protei binding | Guanine nucleotide-binding protein G(i), alpha-1 5KDO_B (98.34/0.000011) |
| GL50803_10708 | Hypothetical protein | 4 | 9 | 4 | 9 | DHS-like NAD/FAD-binding domain superfamily/ Sirtuin family | GO:0070403 NAD+ binding | NAD- dependent protein deacetylase srtuin-2. (E.C.3.5.1.-) 5FYQ_A (00.9e-51) |
| GL50803_94117 | Hypothetical protein | 4 | 6 | 4 | 6 | - | - | Zinc finger homeobox protein 4 2YRK_A (85.51/.76) |
| GL50803_16390 | hypothetical protein | 4 | 4 | 4 | 4 | Ankyrin repeat-containing domain | GO:0005515 protein binding | Transcription factor p65 1OY3_D (97.26/0.00002) |
| GL50803_17172 | Hypothetical protein | 4 | 4 | 4 | 4 | - | - | Pre-mRNA-splicing factor 8, Pre-mRNA-splicing factor 6BK8_L (17.74/1600) |
| GL50803_17400 | Hypothetical protein | 4 | 4 | 4 | 4 | Cyclin PHO80-like | GO:0000079 regulation of cyclin-dependent protein serine/threonine kinase activity GO:0019901 protein kinase binding | Cyclin-dependent protein kinase PHO85 4KRD_B (99.91/2.3e-6) |
| GL50803_90434 | Hypothetical protein | 3 | 4 | 3 | 4 | - | - | Eukaryotic initiation factor 4A-III (E.C.3.6.4.13) 4C9B_B (93.79/.6) |
| GL50803_4266 | Hypothetical protein | 3 | 6 | 3 | 6 | - | - | Eukaryotic translation initiation factor 3 5A5T_A (98.99/0.000002) |
| GL50803_114776 | NSF | 20 | 25 | 20 | 25 | | | |
| GL50803_9558 | Rab1a | 12 | 17 | 12 | 17 | | | |
| GL50803_1695 | Rab11 | 10 | 10 | 10 | 10 | | | |
| GL50803_103676 | Alpha-tubulin | 42 | 56 | 42 | 56 | | | |
| GL50803_101291 | Beta tubulin | 51 | 67 | 51 | 67 | | | |
| GL50803_11654 | Alpha-1 giardin | 18 | 27 | | | | | |
| GL50803_7796 | Alpha-2 giardin | 9 | 22 | | | | | |
| GL50803_11683 | Alpha-3 giardin | 14 | 20 | 14 | 20 | | | |
| GL50803_103373 | Alpha-7.1 giardin | 14 | 15 | | | | | |
| GL50803_114787 | Alpha-giardin | 29 | 40 | | | | | |
| GL50803_4812 | Beta-giardin | 17 | 19 | | | | | |
| GL50803_11434 | 20S proteasome alpha subunit 2 | 3 | 3 | 3 | 3 | | | |
| GL50803_86683 | 26S protease regulatory subunit 7 | 4 | 4 | 4 | 4 | | | |
| GL50803_91643 | 26S proteasome non-ATPase regulatory subunit 2 | 10 | 10 | 10 | 10 | | | |
| GL50803_33166 | 26S proteasome regulatory subunit, putative | 12 | 10 | 12 | 10 | | | |
| GL50803_7110 | Ubiquitin | 7 | 13 | 7 | 13 | | | |
| GL50803_10661 | Ubiquitin-conjugating enzyme E1 | 16 | 18 | 16 | 18 | | | |
| GL50803_9779 | UPL-1 | 48 | 48 | 48 | 48 | | | |
| GL50803_8822 | 2,3-bisphosphoglycerate-independent phosphoglycerate mutase | 4 | 5 | 4 | 5 | | | |
| GL50803_8978 | 4-alpha-glucanotransferase | 16 | 16 | 16 | 16 | | | |
| GL50803_3643 | 70 kDa peptidylprolyl isomerase, putative | 17 | 19 | 17 | 19 | | | |
| GL50803_13608 | Acetyl-CoA synthetase | 40 | 53 | 40 | 53 | | | |
| GL50803_86511 | Acyl-CoA synthetase | 62 | 84 | 62 | 84 | | | |
| GL50803_16667 | Acyl-CoA synthetase | 41 | 41 | 41 | 41 | | | |
| GL50803_12150 | Alanine aminotransferase, putative | 3 | 8 | 3 | 8 | | | |
| GL50803_96460 | Alanyl-tRNA synthetase | 10 | 17 | 10 | 17 | | | |
| GL50803_93358 | Alcohol dehydrogenase | 39 | 42 | | | | | |
| GL50803_15832 | Aminoacyl-histidine dipeptidase | 6 | 11 | 6 | 11 | | | |
| GL50803_8407 | Aminoacyl-histidine dipeptidase | 5 | 7 | 5 | 7 | | | |
| GL50803_112103 | Arginine deiminase | 55 | 78 | | | | | |
| GL50803_10521 | Arginyl-tRNA synthetase | 12 | 13 | 12 | 13 | | | |
| GL50803_10870 | Aspartyl-tRNA synthetase | 12 | 19 | 12 | 19 | | | |
| GL50803_16376 | ATP-dependent RNA helicase p47, putative | 4 | 5 | 4 | 5 | | | |
| GL50803_33689 | Dipeptidyl-peptidase III | 8 | 9 | 8 | 9 | | | |
| GL50803_89347 | DNA-directed RNA polymerase II largest subunit RPB1 | 3 | 5 | 3 | 5 | | | |
| GL50803_11118 | Enolase | 12 | 28 | 12 | 28 | | | |
| GL50803_6304 | Fe-hydrogenase-1 | 5 | 7 | 5 | 7 | | | |
| GL50803_10570 | FKBP-type peptidyl-prolyl cis-trans isomerase | 3 | 4 | 3 | 4 | | | |
| GL50803_8682 | Glucose-6-phosphate 1-dehydrogenase | 6 | 9 | 6 | 9 | | | |
| GL50803_9115 | Glucose-6-phosphate isomerase | 4 | 13 | 4 | 13 | | | |
| GL50803_7195 | Glutamate synthase | 12 | 19 | 12 | 19 | | | |
| GL50803_9348 | Glutaminyl-tRNA synthetase | 9 | 10 | 9 | 10 | | | |
| GL50803_6687 | Glyceraldehyde 3-phosphate dehydrogenase | 26 | 33 | 26 | 33 | | | |
| GL50803_16125 | Glycerol-3-phosphate dehydrogenase | 29 | 37 | | | | | |
| GL50803_6226 | Glycogen phosphorylase | 22 | 31 | | | | | |
| GL50803_9011 | Glycyl-tRNA synthetase | 7 | 9 | 7 | 9 | | | |
| GL50803_9077 | Inositol 5-phosphatase 4 | 4 | 4 | 4 | 4 | | | |
| GL50803_104173 | Isoleucyl-tRNA synthetase | 11 | 14 | 11 | 14 | | | |
| GL50803_24662 | L-serine dehydratase | 6 | 8 | 6 | 8 | | | |
| GL50803_8621 | Leucyl-tRNA synthetase | 27 | 30 | 27 | 30 | | | |
| GL50803_9062 | Long chain fatty acid CoA ligase 5 | 3 | 3 | 3 | 3 | | | |
| GL50803_113892 | Long chain fatty acid CoA ligase, putative | 10 | 18 | | | | | |
| GL50803_16766 | Lysyl-tRNA synthetase | 14 | 16 | 14 | 16 | | | |
| GL50803_14285 | Malic enzyme | 8 | 13 | 8 | 13 | | | |
| GL50803_9508 | Metalloprotease, insulinase family | 5 | 6 | 5 | 6 | | | |
| GL50803_93551 | Metalloprotease, insulinase family | 3 | 4 | 3 | 4 | | | |
| GL50803_9719 | NADH oxidase | 6 | 7 | 6 | 7 | | | |
| GL50803_33769 | NADH oxidase lateral transfer candidate | 23 | 27 | | | | | |
| GL50803_21942 | NADP-specific glutamate dehydrogenase | 22 | 42 | | | | | |
| GL50803_10311 | Ornithine carbamoyltransferase | 55 | 59 | | | | | |
| GL50803_17163 | Peptidyl-prolyl cis-trans isomerase B precursor | 11 | 14 | 11 | 14 | | | |
| GL50803_14404 | Phosphatase | 7 | 10 | 7 | 10 | | | |
| GL50803_13875 | Phosphatase | 3 | 3 | 3 | 3 | | | |
| GL50803_10698 | Phosphorylase B kinase gamma catalytic chain | 8 | 13 | 8 | 13 | | | |
| GL50803_15983 | Prolyl-tRNA synthetase | 17 | 18 | 17 | 18 | | | |
| GL50803_9894 | Protein phosphatase 2A regulatory subunit, putative | 8 | 10 | 8 | 10 | | | |
| GL50803_32312 | Protein phosphatase 2C | 4 | 4 | 4 | 4 | | | |
| GL50803_16567 | Protein phosphatase PP2A regulatory subunit A | 4 | 7 | 4 | 7 | | | |
| GL50803_4079 | Protein phosphatase PP2A regulatory subunit B | 6 | 8 | 6 | 8 | | | |
| GL50803_14993 | Pyrophosphate-fructose 6-phosphate 1-phosphotransferase alpha subunit | 14 | 19 | 14 | 19 | | | |
| GL50803_114609 | Pyruvate-flavodoxin oxidoreductase | 64 | 86 | | | | | |
| GL50803_15214 | Ser/Thr phosphatase | 5 | 6 | 5 | 6 | | | |
| GL50803_7439 | Ser/Thr phosphatase 2A, 65kDa reg sub A | 11 | 20 | 11 | 20 | | | |
| GL50803_11740 | Ser/Thr phosphatase 2C, putative | 6 | 8 | 6 | 8 | | | |
| GL50803_7588 | Serine/threonine protein phosphatase 7 | 10 | 12 | 10 | 12 | | | |
| GL50803_14311 | Serine/threonine protein phosphatase PP-X isozyme 2 | 6 | 12 | 6 | 12 | | | |
| GL50803_12108 | Threonine dehydratase lateral transfer candidate | 12 | 18 | 12 | 18 | | | |
| GL50803_9704 | Transketolase | 32 | 40 | 32 | 40 | | | |
| GL50803_93938 | Triosephosphate isomerase, cytosolic | 3 | 3 | 3 | 3 | | | |
| GL50803_102710 | Ubiquitin carboxyl-terminal hydrolase 14 | 3 | 6 | 3 | 6 | | | |
| GL50803_14460 | Ubiquitin carboxyl-terminal hydrolase 4 | 5 | 9 | 5 | 9 | | | |
| GL50803_29307 | UTP-glucose-1-phosphate uridylyltransferase | 4 | 11 | 4 | 11 | | | |
| GL50803_7532 | Vacuolar ATP synthase catalytic subunit A | 19 | 21 | 19 | 21 | | | |
| GL50803_12216 | Vacuolar ATP synthase subunit B | 7 | 16 | 7 | 16 | | | |
| GL50803_87058 | Vacuolar ATP synthase subunit C | 11 | 16 | 11 | 16 | | | |
| GL50803_14961 | Vacuolar ATP synthase subunit H | 4 | 8 | 4 | 8 | | | |
| GL50803_17327 | Xaa-Pro dipeptidase | 4 | 12 | 4 | 12 | | | |
| GL50803_16453 | Carbamate kinase | 22 | 31 | | | | | |
| GL50803_16034 | Kinase, CAMK CAMKL | 6 | 6 | 6 | 6 | | | |
| GL50803_14364 | Kinase, CAMK CAMKL | 5 | 6 | 5 | 6 | | | |
| GL50803_112076 | Kinase, CDC7 | 3 | 3 | 3 | 3 | | | |
| GL50803_22850 | Kinase, CMGC MAPK | 3 | 5 | 3 | 5 | | | |
| GL50803_12223 | Kinase, NAK | 12 | 16 | 12 | 16 | | | |
| GL50803_16824 | Kinase, NEK | 41 | 51 | 41 | 51 | | | |
| GL50803_113553 | Kinase, NEK | 15 | 31 | 15 | 31 | | | |
| GL50803_11390 | Kinase, NEK | 12 | 19 | 12 | 19 | | | |
| GL50803_9421 | Kinase, NEK | 8 | 12 | 8 | 12 | | | |
| GL50803_16967 | Kinase, NEK | 7 | 14 | 7 | 14 | | | |
| GL50803_101534 | Kinase, NEK | 6 | 6 | 6 | 6 | | | |
| GL50803_26199 | Kinase, NEK | 4 | 4 | 4 | 4 | | | |
| GL50803_8445 | Kinase, NEK | 3 | 8 | 3 | 8 | | | |
| GL50803_2483 | Kinase, NEK | 3 | 3 | 3 | 3 | | | |
| GL50803_17231 | Kinase, NEK | 3 | 4 | 3 | 4 | | | |
| GL50803_15411 | Kinase, NEK-frag | 25 | 30 | | | | | |
| GL50803_8805 | Kinase, SCY1 | 15 | 24 | 15 | 24 | | | |
| GL50803_10609 | Kinase, STE STE20 | 8 | 11 | 8 | 11 | | | |
| GL50803_15514 | Kinase, STE STE20 | 7 | 8 | 7 | 8 | | | |
| GL50803_22165 | Kinase, STE STE20 | 5 | 6 | 5 | 6 | | | |
| GL50803_11301 | Nucleoside diphosphate kinase | 6 | 6 | 6 | 6 | | | |
| GL50803_90872 | Phosphoglycerate kinase | 40 | 53 | 40 | 53 | | | |
| GL50803_3206 | Pyruvate kinase | 24 | 31 | 24 | 31 | | | |
| GL50803_17143 | Pyruvate kinase | 15 | 25 | 15 | 25 | | | |
| GL50803_15410 | Ser/Thr protein kinase | 5 | 14 | 5 | 14 | | | |
| GL50803_1345 | Ribosomal protein L10a | 4 | 5 | 4 | 5 | | | |
| GL50803_16525 | Ribosomal protein L11 | 4 | 16 | 4 | 16 | | | |
| GL50803_14622 | Ribosomal protein L13 | 4 | 8 | 4 | 8 | | | |
| GL50803_8001 | Ribosomal protein L15 | 6 | 6 | 6 | 6 | | | |
| GL50803_19436 | Ribosomal protein L18a | 6 | 10 | | | | | |
| GL50803_14329 | Ribosomal protein L19 | 3 | 7 | 3 | 7 | | | |
| GL50803_10091 | Ribosomal protein L23 | 16 | 17 | 16 | 17 | | | |
| GL50803_10367 | Ribosomal protein L27 | 4 | 3 | | | | | |
| GL50803_14321 | Ribosomal protein L3 | 11 | 11 | 11 | 11 | | | |
| GL50803_8001 | Ribosomal protein L30 | 6 | 6 | 6 | 6 | | | |
| GL50803_17547 | Ribosomal protein L4 | 17 | 33 | | | | | |
| GL50803_5593 | Ribosomal protein L7 | 6 | 4 | 6 | 4 | | | |
| GL50803_17244 | Ribosomal protein L7a | 15 | 21 | | | | | |
| GL50803_7878 | Ribosomal protein S14 | 9 | 11 | | | | | |
| GL50803_16387 | Ribosomal protein S2 | 4 | 31 | 4 | 31 | | | |
| GL50803_16431 | Ribosomal protein S24 | 3 | 7 | 3 | 7 | | | |
| GL50803_16265 | Ribosomal protein S3a | 15 | 26 | | | | | |
| GL50803_8462 | Ribosomal protein S6 | 4 | 4 | 4 | 4 | | | |
| GL50803_8118 | Ribosomal protein S7 | 3 | 9 | 3 | 9 | | | |
| GL50803_14620 | Protein 21.1 | 14 | 6 | | | | | |
| GL50803_27925 | Protein 21.1 | 20 | 19 | 20 | 19 | | | |
| GL50803_21505 | Protein 21.1 | 16 | 28 | 16 | 28 | | | |
| GL50803_32778 | Protein 21.1 | 6 | 27 | 6 | 27 | | | |
| GL50803_14254 | Protein 21.1 | 7 | 11 | 7 | 11 | | | |
| GL50803_24842 | Protein 21.1 | 6 | 9 | 6 | 9 | | | |
| GL50803_16220 | Protein 21.1 | 9 | 6 | 9 | 6 | | | |
| GL50803_102813 | Protein 21.1 | 8 | 15 | 8 | 15 | | | |
| GL50803_17568 | Protein 21.1 | 4 | 19 | 4 | 19 | | | |
| GL50803_8174 | Protein 21.1 | 3 | 4 | 3 | 4 | | | |
| GL50803_17060 | Protein 21.1 | 19 | 28 | 19 | 28 | | | |
| GL50803_6430 | 14-3-3 protein | 34 | 52 | 34 | 52 | | | |
| GL50803_10358 | A-type flavoprotein lateral transfer candidate | 11 | 12 | 11 | 12 | | | |
| GL50803_15048 | ATP-dependent RNA helicase-like protein | 5 | 12 | 5 | 12 | | | |
| GL50803_16202 | Axoneme central apparatus protein | 4 | 5 | 4 | 5 | | | |
| GL50803_137716 | Axoneme-associated protein GASP-180 | 23 | 26 | | | | | |
| GL50803_16745 | Axoneme-associated protein GASP-180 | 5 | 6 | | | | | |
| GL50803_17121 | Bip | 30 | 37 | | | | | |
| GL50803_15148 | Chaperone protein dnaJ | 17 | 21 | 17 | 21 | | | |
| GL50803_9808 | Chaperone protein DnaJ | 10 | 29 | 10 | 29 | | | |
| GL50803_11953 | Coatomer alpha subunit | 7 | 9 | 7 | 9 | | | |
| GL50803_113677 | Coiled-coil protein | 9 | 11 | 9 | 11 | | | |
| GL50803_17105 | Coiled-coil protein | 5 | 3 | 5 | 3 | | | |
| GL50803_88765 | Cytosolic HSP70 | 90 | 120 | 90 | 120 | | | |
| GL50803_7662 | Dynein heavy chain | 3 | 4 | 3 | 4 | | | |
| GL50803_33218 | Dynein intermediate chain | 7 | 7 | 7 | 7 | | | |
| GL50803_15106 | Elongation factor 1-alpha | 99 | 8 | 99 | 8 | | | |
| GL50803_12102 | Elongation factor 1-gamma | 22 | 42 | 22 | 42 | | | |
| GL50803_17570 | Elongation factor 2 | 84 | 109 | 84 | 109 | | | |
| GL50803_14614 | Eukaryotic translation initiation factor 5A | 7 | 8 | 7 | 8 | | | |
| GL50803_15869 | GTP-binding nuclear protein RAN/TC4 | 7 | 24 | 7 | 24 | | | |
| GL50803_98054 | Heat shock protein HSP 90-alpha | 25 | 36 | 25 | 36 | | | |
| GL50803_13864 | Heat shock protein HSP 90-alpha | 17 | 32 | 17 | 32 | | | |
| GL50803_16412 | Heat-shock protein, putative | 12 | 17 | 12 | 17 | | | |
| GL50803_121045 | Histone H2B | 4 | 4 | | | | | |
| GL50803_103059 | IFT complex B | 3 | 3 | 3 | 3 | | | |
| GL50803_13500 | Importin beta-3 subunit | 8 | 17 | 8 | 17 | | | |
| GL50803_102101 | Kinesin-3 | 23 | 23 | | | | | |
| GL50803_102101 | Kinesin-3 | 6 | 32 | 6 | 32 | | | |
| GL50803_112846 | Kinesin-3 | 3 | 32 | 3 | 32 | | | |
| GL50803_14521 | Peroxiredoxin 1 | 34 | 45 | | | | | |
| GL50803_10167 | Polyadenylate-binding protein, putative | 7 | 6 | 7 | 6 | | | |
| GL50803_13452 | Regulator of nonsense transcripts 1-like protein | 6 | 7 | 6 | 7 | | | |
| GL50803_103855 | RNase L inhibitor | 4 | 5 | 4 | 5 | | | |
| GL50803_2562 | Sec31 | 6 | 6 | 6 | 6 | | | |
| GL50803_7031 | Spindle pole protein, putative | 4 | 12 | 4 | 12 | | | |
| GL50803_17055 | Spindle pole protein, putative | 4 | 7 | 4 | 7 | | | |
| GL50803_27310 | Stress-induced-phosphoprotein 1 | 10 | 14 | 10 | 14 | | | |
| GL50803_9825 | TBP-interacting protein TIP49 | 6 | 6 | 6 | 6 | | | |
| GL50803_11992 | TCP-1 chaperonin subunit epsilon | 3 | 4 | 3 | 4 | | | |
| GL50803_16124 | TCP-1 chaperonin subunit eta | 11 | 13 | 11 | 13 | | | |
| GL50803_5942 | TCP-1 chaperonin subunit theta | 10 | 10 | 10 | 10 | | | |
| GL50803_16568 | Transcription factor, putative | 3 | 3 | 3 | 3 | | | |
| GL50803_13561 | Translation elongation factor | 26 | 27 | 26 | 27 | | | |
| GL50803_10255 | Translation initiation factor eIF-4A, putative | 18 | 27 | 18 | 27 | | | |
| GL50803_137703 | Vacuolar protein sorting 29 | 4 | 5 | 4 | 5 | | | |
| GL50803_23833 | Vacuolar protein sorting 35 | 13 | 15 | 13 | 15 | | | |
