## supplemental table 8 for "Phosphoinositide-binding proteins mark, shape and functionally modulate highly-diverged endocytic compartments in the parasitic protist *Giardia lamblia*"

Supplemental Table 8: List of oligonucleotide names and sequences for construct synthesis

| Primer pair | Construct | Forward primer (5’ to 3’ direction) | | Reverse primer (5’ to 3’ direction) | |
|---|---|---|---|---|---|
| 1 | **Pendo-HA-GIPXD3 (Promoter+HA fragment)** | | | pE_HA_16596_prom_Avr_as | GCCCTAGGCGCGTAGTCTGGGACATCGTATGGGTACATAAAGTTTTACTTACATATTTAATCGAAC |
| 2 | **Pendo-HA-GIPXD3 (ORF fragment)** | pE_HA_16596_orf_Avr_s | GCCCTAGGTCTAATAGGTTTACCACCCAGACG | pE_HA_16596_orf_Pac_as | GCTTAATTAATCACTCTTTAAGAGCATAAGTGATCGGC |
| 3 | **Pendo-HA-GIPXD4 (Promoter+HA fragment)** | pE_HA_42357_prom_XbaI_s | 5'-GCT CTA GAG TCA ATT CGG TCG TTA GCA TAC CGA CTC GTC TCA GTG AAC CGT G-3' | pE_HA_42357_prom_NotI_as | GCGCGGCCGCCGCGTAGTCTGGGACATCGTATGGGTACATTTTTTTAAGATACAGAGGCAAAAGGC |
| 4 | **Pendo-HA-GIPXD4 (ORF fragment)** | pE_HA_42357_orf_NotI_s | GCGCGCGGCCGCAGACGGTGATCAGGAGCAGG | pE_HA_42357_orf_Pac_as | GCTTAATTAATCAGCAGCGAGCTGCCTTTAAGAA |
| 5 | **Pendo-HA-GIPXD5 (Promoter+HA fragment)** | | | pE_HA_16548_prom_Avr_as | GCCCTAGGCGCGTAGTCTGGGACATCGTATGGGTACATTCACACCAAGAAACCTAGACC |
| 6 | **Pendo-HA-GIPXD5 (ORF fragment)** | pE_HA_16548_orf_Avr_s | GCCCTAGGATAAAAAACGTACTCATGTTTGCAGCAG | pE_HA_16548_orf_Pac_as | GCTTAATTAACTATGTGTATCCTTCCTCAATAGAAC |
| 7 | **Pendo-HA-GIPXD6 (Promoter+HA fragment)** | pE_HA_24488_prom_Xba_s | GCTCTAGACAGGAGGATGCTTAAATCCTTTATT | pE_HA_24488_prom_Avr_as | GCCCTAGGCGCGTAGTCTGGGACATCGTATGGGTACATCATTTTTAAATAGCTTCCCTGCGC |
| 8 | **Pendo-HA-GIPXD6 (ORF fragment)** | pE_HA_24488_orf_Avr_s | GCCCTAGGACGCACGTTGTTTCGACTGCAT | pE_HA_24488_orf_Pac_as | GCTTAATTAATCACCTCAAGGTGTAGCTCTC |
| 9 | **Pendo-GINECAP1-HA** | pE17195Xba_F | GCTCTAGAGTGTACCGACAGGAATGATACGCTC | pE17195HAPac_B | GCTTAATTAATTACGCGTAGTCTGGGACATCGTATGGGTACTTGGGAGGCGCCCCAAAATTG |
| 10 | **Pendo-GIBAR1-HA** | 15847_xbaI_fw | GCtctagaGTGTACGTCACAAATATTTACTGTAATCAGATCAAGCATTAG | 15847_pacI_HA_rv | CGTTAATTAACTACGCGTAGTCTGGGACATCGTATGGGTAGACAATATTAGCACGGCAGCGGTCTTTTTCTG |
| 11 | **Pendo-HA-GIFERM (Promoter+HA fragment)** | P115468_Spe _for | GC ACT AGT CTG TCT GAA ATG CAT TCA TAG TTA TG | P115468_Sph_ATGHA_rev | GC GCATGC CGC GTA GTC TGG GAC ATC GTA TGG GTA CAT TGA ATC A ATC AAC TCT TTT AAA GTT CAT TG |
| 12 | **Pendo-HA-GIFERM (ORF fragment)** | Sph_115468_for | GC GCATGC GAT CTT GCA CGC TTT GCC CAC | 115468_Pac_rev | CG TTAATTAA TCA TA GAA TAT GCT CCA GTG CAC |
| 13 | **Pendo-GIPROP2-HA** | P16957_Xba_for | GC TCTAGA CGA TGA TCC TTT TTC ATG TCA GTC AG | 16957_HA_Pac_rev | GC TTAATTAA TCA CG CGT AGT CTG GGA CAT CGT ATG GGT A CT TAA GAC TCC AAA CGC TTT GCA G |
| 14 | **Pendo-GI16801-HA** | P16801_Xba_for | GC tctaga ATG CTC TAC GCC CAC CTC GAG | 16801_V5s_Avr_rev | GC cctagg TTA atcaagtccaagaagaggattaggaat GACAGCTTCGGAGATCTCTGTG |
| 15 | **MBP::HA::*GI* PXD1** | MBP-PX-7723 fw ecoR1 HA | gc GAATTC ATGTACCCATACGATGTCCCAGACTACGCGACCATACGCATACAGGTATCTG | MBP-PX7723 as Xba | gcTCTAGACTAGTGGCGATGCATCCCCT |
| 16 | **MBP::HA::*GI* PXD2** | MBP-PX16595 fw ecoR1 HA | gc GAATTC ATGTACCCATACGATGTCCCAGACTACGCGATCGAGGCCCCTGTGCGAC | MBP-PX16595 as Xba | gcTCTAGATCACCGGTGATATACAAGGATAT |
| 17 | **MBP::HA::*GI* PXD3** | MBP-PX16596 fw ecoR1 HA | gc GAATTC ATGTACCCATACGATGTCCCAGACTACGCGCAGGTCAAAGACATAAAAAGAAC | MBP-PX16596 as xba | gcTCTAGACTACTCTTTAAGAGCATAAGTGA |
| 18 | **MBP::HA::*GI* PXD4** | MBP-PX42357 fw ecoR1 HA | gc GAATTC ATGTACCCATACGATGTCCCAGACTACGCGCATGGAGCTGCATCGTCCAT | MBP-PX42357 as xba | gcTCTAGATCAGCAGCGAGCTGCCTTTA |
| 19 | **MBP::HA::*GI* PXD5** | MBP-PX16548 fw ecoR1 HA | gc GAATTC ATGTACCCATACGATGTCCCAGACTACGCGGCCGAAGAGACGGGGGCG | MBP-PX16548 as xba | gcTCTAGACTATGTGTATCCTTCCTCAATA |
| 20 | **MBP::HA::*GI* PXD6** | MBP-PX24488 fw ecoR1 HA | gc GAATTC ATGTACCCATACGATGTCCCAGACTACGCGGTCACACCTGCGAGGCTTAG | MBP-PX24488 as Xba | gcTCTAGACTACCTCAAGGTGTAGCTCTC |
| 21 | **MBP::HA::*GI* FYVE** | 16653MBP_fw | 5GCG AAT TCT ACC CAT ACG ATG TCC CAG ACT ACG CGC TCT CCA TCG CAA TTC CTC ACA AGA C-3 | 16653MBP_bw | GCTCTAGATCAGGGAGGCCCGTCCACCAC |
| 22 | **MBP::HA::*GI* NECAP1** | 17195MBP_fw | gcGAATTCTACCCATACGATGTCCCAGACTACGCGCTCTCCATCGCAATTCCTCACAAGAC | 17195MBP_bw | GCTCTAGATTACTTGGGAGGCGCCCCAAAATTG |
| 23 | **pCWP1-2xFYVE::GFP** | pSec2xFyveGFp_s | GCACTAGTATGGATGGTACCGAGCTCGAAAG | pSec2xFyveGFp_as | GCTTAATTAATTACTTGTACAGCTCGTCCATG |
| 24 | **pCWP1-GFP::P4C** | pSecGFP4CSidC_s | GCACTAGTATGCAGGTGAGCAAGGGCGAG | pSecGFP4CSidC_as | GCTTAATTAACTAAAAGAATTCAATTGCTTCATC |
| 25 | **Pendo-16811-HA** | pE16811HA_SpeF | GCACTAGTGTGAAGAGTATGTCATGCGTG | pE16811_HAPacIB | GCTTAATTAACTACGCGTAGTCTGGGACATCGTATGGGTACTTTTTTCCATTTTGATCTGTAGAG |
| 26 | **Pendo-103709-HA** | pE103709fwAVR2 | GCCCTAGGCTGACGACCTCGACGATAATT | pE103709revHApac | gcttaattaaCTACGCGTAGTCTGGGACATCGTATGGGTACTGTGTGGTGATTGTGATGGG |
| 27 | **Pendo-NSF-HA (part A)** | pE114776.NheI.f | CG GCTAGC AAGGCGAGGCCAACTTGAAGAG | 114776_partA_rv | CG GCATGC A CAG GAG GAT CAC GCC CTG AG |
| 28 | **Pendo-NSF-HA (part B)** | 114776_partB_fw | CG GCATGC CAACGTTTCATAGAACAAAGTGGGCAG | pE114776HAPac.b | GCTTAATTAATCACGCGTAGTCTGGGACATCGTATGGGTAGGATAGCGTTGCATTCAATGTCCTG |
| 29 | **Pendo-HA-5785 (Promoter+HA fragment)** | 5785prom.AvrII.f | 5'-GCT CTA GAC ATT TGA GGA CTT GTG GGG AAT TCA CTA AAC GTG CCA G-3' | 5785prom.SphI.b | GCGCATGCCCAGCCTTTCTTTTTTAATGTTTTTGAAC |
| 30 | **Pendo-HA-5785 (ORF fragment)** | pE5785orf.SphI.f | GCGCATGCATGTACCCATACGATGTCCCAGACTACGCGGAAGAGATAGAATGTTCACTCATGG | pE5785orf.PacI.b | GCTTAATTAATCAATATCTGATCTCTGAGGGCTC |
| 31 | **Pendo-HA-7309 (Promoter+HA fragment)** | 7309prom.AvrII.f | GCCCTAGGGTATGTGCTCCATCATAGTTCTCAG | 7309prom.Xho.b | GCCTCGAGCGGTGAGCAATGCCAATTACCG |
| 32 | **Pendo-HA-7309 (ORF fragment)** | pE7309orf.XhoI.f | GCCTCGAGATGTACCCATACGATGTCCCAGACTACGCGGAGAACATGTATGACGACTTTATAGAC | pE7309orf.PacI.b | GCTTAATTAATTACTTAAAGATTGTGCTCAGGACG |
| 33 | **Pendo-HA-10013 (Promoter+HA fragment)** | 10013prom.XbaI.f | 5'-GCT CTA GAA ACC TGA GAG CCC AGG CTT TCC CAG AAG TGA AGA TTC ATT ATC GTC-3' | 10013prom.SphI.b | GCGCATGCTTTTATGTTTTTTATTTTCTTCTACTCGTC |
| 34 | **Pendo-HA-10013 (ORF fragment)** | pE10013orf.SphI.f | GCGCATGCATGTACCCATACGATGTCCCAGACTACGCGGACGATTCTGGTAACGATATTGGAG | pE10013orf.PacI.b | GCTTAATTAATTAAGGACACTGCCACGTGAATC |
| 35 | **Pendo-HA-14469 (Promoter+HA fragment)** | pE14469.fl.Xba.fw | GCTCTAGAAAATAGACACACAAGCAAGGGAGAAGAG | pE14469HA.AvrII.bw | GCCCTAGGCGCGTAGTCTGGGACATCGTATGGGTACATAATTTCTCTTTTGGGTTTTGCCACACTAATG |
| 36 | **Pendo-HA-14469 (ORF fragment)** | pE14469orf.AvrII.fw | GCCCTAGGAGTGCCGGGAACATTTATGCGCTC | 14469.fl.pac.bw | GCTTAATTAA TTAAGAAAACTTGATCGCGATGGGAACG |
| 37 | **Pendo-HA-9605 (Promoter+HA fragment)** | pEHA9605AvrII.f | GCCCTAGGCATATGGAGCGCCCCCTAC | pEHA9605prom.SpeI.bw | GCACTAGTTGATTCTGAGATTGACTATGGAATCAATC |
| 38 | **Pendo-HA-9605 (ORF fragment)** | pEHA9605orfSpeI.fw | GCACTAGTATGTACCCATACGATGTCCCAGACTACGCGATGATTGGATTAAAAAACCAAAAAGATCTTTTAAAC | pEHA9605orfPac.bw | GCTTAATTAATCACCTGCTGCGCTTTTTCTTAG |
| 39 | **PCWP1-HA-9605-S74N** | 9605_S74N_s | AACACCTTCGTCAACGGATTCATTAAC | 9605_ S74N_as | CTTCCCGACCCCGGTTTC |
| 40 | **PCWP1-HA-9605-K73E** | 9605_K73E_s | 5'-GAA ACC GGG GTC GGG GAG TCC ACC TTC GTC-3' | 9605_ K73E_as | GAC GAA GGT GGA CTC CCC GAC CCC GGT TTC-3' |
| 41 | **Pendo-HA-16717 (Promoter+HA fragment)** | 16717prom.XbaI.f | GCTCTAGAACCTTCCTGAGCTCGAAGGAC | 16717prom.SpeI.b | GCACTAGTCCTGCAGAATATTCATGTTGGGCTG |
| 42 | **Pendo-HA-16717 (ORF fragment)** | pE16717orf.SpeI.f | GCACTAGTATGTACCCATACGATGTCCCAGACTACGCGCCCATCACAGAGGCCGTTATC | pE16717orf.PacI.b | GCTTAATTAATCACTTCTGGACCTGCCCGG |
| 43 | **Pendo-GINECAP-APEX2::2xHA (ORF fragment)** | pE17195Xba_F | GCTCTAGAGTGTACCGACAGGAATGATACGCTC | 17195.AvrIIbw | gcCCTAGGCTTGGGAGGCGCCCCAAAATTG |
| 44 | **Pendo-GINECAP-APEX2::2xHA (APEX fragment)** | APEX2HA fw AVR2 | gcCCTAGGGGCGGATCAGGCTCTGGAAA | APEX2HA rev pac | gcttaattaaTTAGGCGTAGTCAGGGACATCATA |
| 45 | **Pendo-GINECAP1ΔWVIF-HA** | ΔPH_fw | TGATTGGCCTGGTGAGCAGGGCAACCTCAGCAT | ΔPH_bw | ATGCTGAGGTTGCCCTGCTCACCAGGCCAATCA |
| 46 | **pCWP1-NT-GIFYVE::HA (ORF fragment)** | pC16653HA fw avr | GCCCTAGGATGTATTCGGGTCCTTACAATA | pC16653HA_NT_rev sph | GCGCATGCCCCGTCCACCACAAGCATGA |
| 47 | **pCWP1-CT-GIFYVE::HA (ORF fragment)** | pC16653HA_CT fw avr | GCCCTAGGATGCCTCCCAATCCCCTTGCTTAC | pE16653 rev Sph | GCGCATGCCGCCAGCTCCTCACTGTGAA |
