## supplemental table 9 for "Phosphoinositide-binding proteins mark, shape and functionally modulate highly-diverged endocytic compartments in the parasitic protist *Giardia lamblia*"

Supplemental Table 9: Amino acid sequences of lipid-binding modules used in vitro for protein lipid-overlay assay

| | Amino-acid position | Amino-acid sequence |
|---|---|---|
| ***Gl* PXD1** | 553-691 | TIRIQVSEHNALHLPRSLFFTPDGKYKNYVRYLDYRKVFDRFTHTQVLMFIFAVSSQERKLK<br>CISKTYEDFHVLHKELVGAFGYSAVPGLPSFRSMNMGGGRHEVIHDTQKRLTKYLTEINNV<br>QALRTSAIYRSFLKHQ |
| ***Gl* PXD2** | 1665-1801 | IEAPVRPGYESSLLSYVKVSSDPTTCHPTKFVARLTAHTWCIEQKSKPYVVYTFTISTEKGV<br>FEVAKRFSELRELDKALRRVFPTVTLPKYPSSMGGKFTDEQLEERKRLLGRYMSQLNQIAV<br>IRTSQVYKEFIQMK |
| ***Gl* PXD3** | 443-587 | QVKDIKRTKSPKSPKRSGASEKNDNSRKERKGTSEARVLSYEVCSTGSGHPFVVYKIELPS<br>QDGGESVVIHKRMKEIKAFHKRLQRNFPSITLPKLPQDRPWALGGNKLKFVQERSRLLEAY<br>CGDINHNAMIYKSRIYKDYINEQ |
| ***Gl* PXD4** | 883-1023 | HGAASSIVRSNKHRDSGRHVSLFVDTSSCSSFYVKVLSHRTVIISGSRPFISYYIELGMDGR<br>RLHISCRFSEVKEHAAISAAYPAATLPDIPPDRPYALGGWKPEFIRERARLLEAYLYSMHSIT<br>FVRTSNIYRQFLKAA |
| ***Gl* PXD5** | 90-233 | AEETGALSAIPSAFPPWGKPTVYDPQNPITHITAVIVDQPHTMMDRNHPYTQYMIFKSELPA<br>NKRYYMVYRRFNDVKRLAATLLDLKVGPLPSLPPQRSVFQSLYDPLFEDRRKRLAEYLQLL<br>SSTLSVVQTDAFQHFLTEH |
| ***Gl* PXD6** | 39-178 | VTPARLRNEGGCVTQTPIITETKESSDDPPPLSVCDNYTVCSNGGSHPFIVYDLLIPRKKHQ<br>VQVSKRFSTIKRLHADLLQEGLEHLPSFPRDRPWNLGGNRKGFVEERVREINNYFHRLLS<br>NKAVVNSNAFKYFLSEND |
| ***Gl* FYVE** | 218-300 | LSIAIPHKTLVPRTEWIQDKDVNNCQGCDTAFTLLNRRHHCRRCGRIFCNACCIKPIDILSDT<br>GTDRLCKLCKAVMLVVDGPP |
| ***Gl* NECAP1** | 1-178 | MEFEQVLATVTQVVAYKLPPTLNLSSFKCADWPGEWVIFQGNLSIISKGEACSVSLVAPDT<br>GAEAARFPIEYKGTVPVVEKASDSSRYFVIVVKDPTGAKMAFIGIGFQERDGAFAFQAALA<br>DHGKLLDRKAHPPAQIVVNQDFSLKSGEKIKIGLGKKTGGTPSVPPGGFNFGAPPK |
